# Spatial dynamics of mammalian brain development and neuroinflammation by multimodal tri-omics mapping

**DOI:** 10.1101/2024.07.28.605493

**Authors:** Di Zhang, Leslie A Rubio Rodríguez-Kirby, Yingxin Lin, Mengyi Song, Li Wang, Lijun Wang, Shigeaki Kanatani, Tony Jimenez-Beristain, Yonglong Dang, Mei Zhong, Petra Kukanja, Shaohui Wang, Xinyi Lisa Chen, Fu Gao, Dejiang Wang, Hang Xu, Xing Lou, Yang Liu, Jinmiao Chen, Nenad Sestan, Per Uhlén, Arnold Kriegstein, Hongyu Zhao, Gonçalo Castelo-Branco, Rong Fan

## Abstract

The ability to spatially map multiple layers of the omics information over different time points allows for exploring the mechanisms driving brain development, differentiation, arealization, and alterations in disease. Herein we developed and applied spatial tri-omic sequencing technologies, DBiT ARP-seq (spatial ATAC–RNA–Protein-seq) and DBiT CTRP-seq (spatial CUT&Tag– RNA–Protein-seq) together with multiplexed immunofluorescence imaging (CODEX) to map spatial dynamic remodeling in brain development and neuroinflammation. A spatiotemporal tri-omic atlas of the mouse brain was obtained at different stages from postnatal day P0 to P21, and compared to the regions of interest in the human developing brains. Specifically, in the cortical area, we discovered temporal persistence and spatial spreading of chromatin accessibility for the layer-defining transcription factors. In corpus callosum, we observed dynamic chromatin priming of myelin genes across the subregions. Together, it suggests a role for layer specific projection neurons to coordinate axonogenesis and myelination. We further mapped the brain of a lysolecithin (LPC) neuroinflammation mouse model and observed common molecular programs in development and neuroinflammation. Microglia, exhibiting both conserved and distinct programs for inflammation and resolution, are transiently activated not only at the core of the LPC lesion, but also at distal locations presumably through neuronal circuitry. Thus, this work unveiled common and differential mechanisms in brain development and neuroinflammation, resulting in a valuable data resource to investigate brain development, function and disease.

## MAIN TEXT

The development of the mammalian brain is a meticulously regulated and dynamic process that involves the genesis, differentiation, and maturation of diverse cellular lineages. Cellular heterogeneity within each cellular lineage, including neurons and glia, provides a broad landscape that balances cellular distinction and collectivity contributing to the complex architecture and multifunctional capabilities of the central nervous system (**CNS**) at large^1,2^. Neuroinflammation causes substantial remodeling of cellular architecture in the brain and may draw parallels and distinctions with development.

Mouse corticogenesis occurs through an inside-out mechanism in which cells of the deeper cortical layers such as corticothalamic (**CTPN**) and callosal projection neurons (**CPN**) (embryonic day (E)12.5, E12.5) in layer VI and subcerebral (**SCPN**) and CPN (E13.5) in layer V are defined earlier than those that populate the outer layers including pyramidal neurons (E14.5) in layer IV and CPNs (E15.5)^1^ in layer II/III. Postnatal development mainly involves further migration, maturation, axonal guidance (for upper layers), synaptogenesis and dendritic branching, representing spatially dynamic and diverse processes. Glial genesis occurs simultaneously, with oligodendrocytes and astrocytes being specified and populating the cerebral cortex (**CTX**) and the underlying white matter (**WM**), including the corpus callosum (**CC**). Oligodendrocytes are generated in three waves; the first wave begins at E12.5, the second at E15.5 and the final wave beginning as early as E17.5 and mainly occurring postnatally at P0 (postnatal day 0)^3,4^. During the postnatal timeframe, oligodendrocyte precursor cells (**OPCs**) are proliferating, migrating and differentiating into oligodendrocytes allowing for myelination to begin around P10 and reach completion by P21^3^. Astrocytes are born starting around E16 but only reach full regional heterogeneity via their postnatal development^5^. Understanding developmental programs assists in elucidating the mechanisms of disease states and how to promote repair.

Reparative mechanisms due to injury, demyelination or neurodegeneration recycle many of the developmental programs but can also differ depending on cell-cell interaction, regional context and the influence of the immune system. Disease-associated states of glia have been a primary focus in recent years along with the juxtaposition of health and development. This is chiefly due to the ability of large-scale sequencing strategies to delineate the nuance, transience and stability of cell states by interrogating the RNA and protein expression profiles and importantly the chromatin accessibility and architecture^6–13^.

Spatial omics technologies—including spatial epigenomics, transcriptomics and proteomics— leverage either next-generation sequencing (**NGS**) or imaging techniques to provide an unprecedented level of insight into the molecular architecture of biological systems. One such imaging-based spatial proteomics technology, Co-detection by indexing (**CODEX**), provides morphological information at single-cell resolution^14^. On the NGS front, to thoroughly understand gene regulation mechanisms, we developed a platform approach named deterministic barcoding in tissue (**DBiT**), which realized the spatial co-profiling of epigenome and transcriptome^15^, as well as spatial co-mapping of transcriptome and proteome^16^. It is desirable to further realize the co-profiling of epigenome, transcriptome, and proteome altogether from the same tissue section in a spatially resolved manner to investigate the molecular mechanisms across all the layers of the central dogma^17^. Herein, we developed and applied two DBiT-based spatial tri-omic technologies, including spatial <u>A</u>TAC–<u>R</u>NA–<u>P</u>rotein-seq (abbreviated as **DBiT ARP-seq**) and spatial <u>C</u>UT&<u>T</u>ag–<u>R</u>NA–<u>P</u>rotein-seq (abbreviated as **DBiT CTRP-seq**), for simultaneous profiling of genome-wide chromatin accessibility or histone modifications (H3K27me3), the entire transcriptome, and the proteome (∼150 proteins) within the same tissue section at cellular level.

To elucidate the cellular and molecular processes during the mammalian brain development and compare with the neuroinflammatory dynamics during demyelination and remyelination in the lysolecithin (**LPC**) mouse model of multiple sclerosis (**MS**), we applied both CODEX and DBiT-based spatial tri-omic sequencing to mouse brains from newborn to juvenile stages (P0, P2, P5, P10, P21), the human brain primary visual cortex (V1) from prenatal to postnatal stages (second trimester, third trimester, and infancy), the LPC mouse brain lesions at both peak demyelination (5 days post-lesion, 5 DPL) and peak remyelination (21 days post-lesion, 21 DPL) phases (**Fig. 1a and Extended Data Fig. 1**). The generated datasets are accessible as an online resource and can be explored within the spatial coordinates of the tissue at (https://spatial-omics.yale.edu/).

**Fig. 1:**
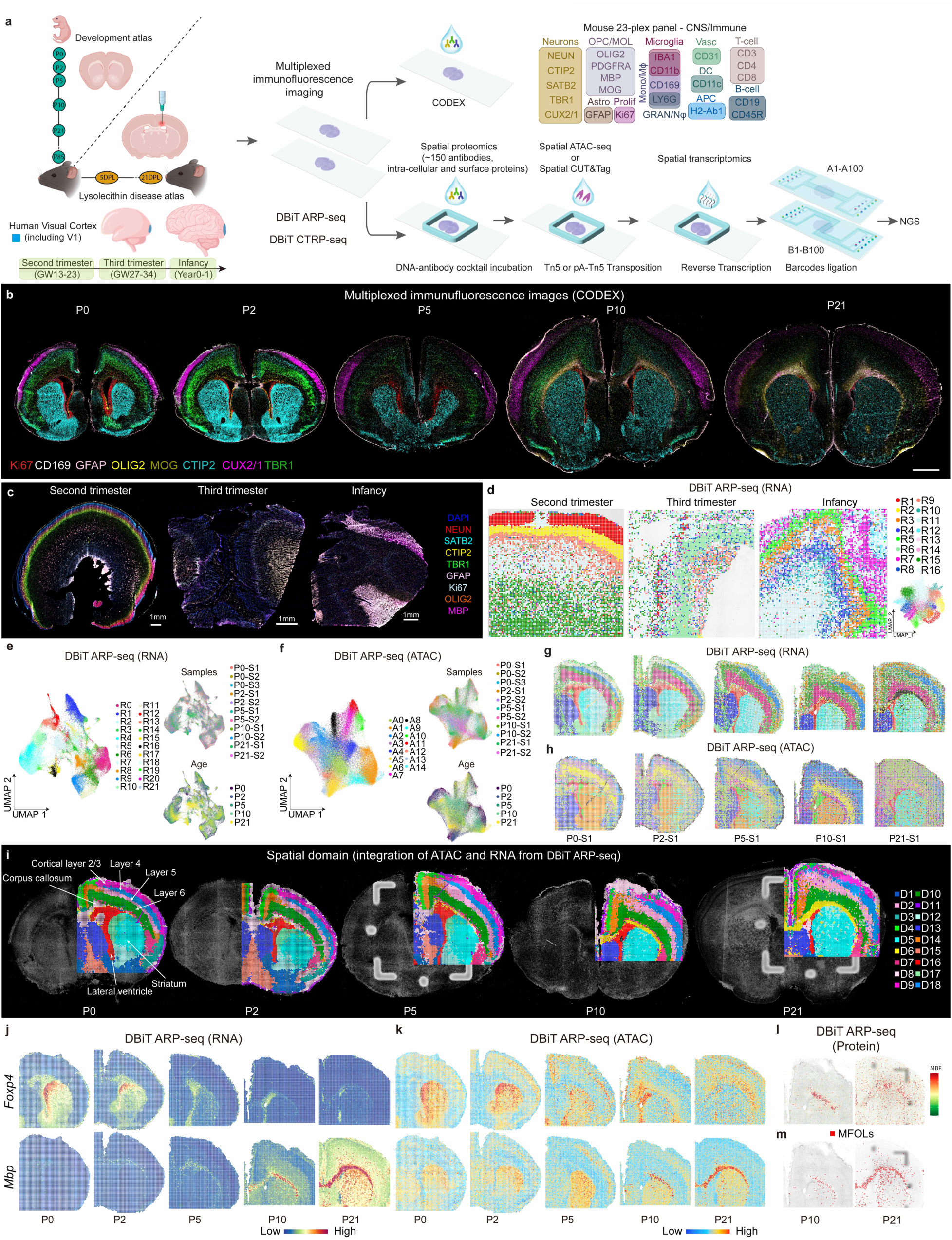
Spatial epigenome–transcriptome–proteome mapping of the developing mouse and human brains at the onset of myelination. **a**, Schematic workflow. **b,** CODEX images of whole mouse brains at P0 to P21. Scale bar, 1 mm. **c,** CODEX images of human brain V1 region at second trimester, third trimester, and infancy. Scale bar, 1 mm. **d,** UMAP and spatial distribution of spatial RNA clusters of the human brain samples at different stages. **e,** UMAPs of the mouse brain spatial RNA clusters at different ages from P0 to P21. **f,** UMAPs of the mouse brain spatial ATAC clusters at different ages from P0 to P21. **g,** Spatial distribution of RNA clusters in **e**. **h,** Spatial distribution of ATAC clusters in **f**. **i,** Spatial domain of the mouse brains generated by the integration of RNA and ATAC data in DBiT ARP-seq. **j-k,** Spatial mapping of gene expression (**j**) and gene activity score (GAS) (**k**) of *Foxp4* and *Mbp* for ATAC and RNA in DBiT ARP-seq. **l,** MBP expression from ADT protein data in DBiT ARP-seq. **m,** Label transfer of MFOLs from scRNA-seq^24–26^ to P10 and P21 mouse brains.

### Multiplex immunofluorescence imaging reveals contrasting spatial dynamics of layer-defining transcription factors and myelination

To generate the proteomic and cell type maps of mouse and human brains with single-cell resolution, we designed a 23-plex CODEX antibody panel for mouse (**Fig. 1a and Extended Data Fig. 2**) and a 19-plex panel for human (**Extended Data Fig. 3a**), which include the canonical markers for all major cell types in the developing brain. We imaged the mouse brain coronal sections (Bregma; 0.5 mm for P21 and closely analogous positions for P0-P10) from P0 to P21 as well as human brain V1 regions at second trimester (gestational week 13-23, GW13-23), third trimester (GW27-34), and infancy (Year 0-1). Segmentation, Seurat clustering, and cell typing revealed the expected arealization of the mouse brain at all analyzed stages with expected marker distribution, specificity and morphology (**Fig. 1b and Extended Data Fig. 2a-c**).

In the mouse samples, the ramified astrocytic marker GFAP was expressed near the meninges and medial ventricular zones at P0, starting to expand at P5 to specific areas in the parenchyma and corpus callosum, a region where it was particularly abundant at P21 (**Fig. 1b and Extended Data Fig. 2c**). To monitor cellular proliferation—a crucial indicator of development, immune response, and repair—we utilized Ki67, a common proliferation marker to visualize proliferating cells. Ki67+ cells were densely concentrated and showed prominent nuclear staining in the ventricular region, where neuroblasts, precursor cells and glia reside^18,19^. The Ki67 periventricular density contracted during later developmental stages along with its overall diffuse abundance in the corpus callosum and cortical layers indicating a shift from progenitor to terminally differentiated cell status. Nevertheless, a proliferative pool remained through development (**Fig. 1b and Extended Data Fig. 2c**).

All neuronal markers displayed morphology with positional patterning that populated the gray matter regions, with sparsity in WM structures like the corpus callosum and anterior commissure. NEUN stained mature neurons throughout the brain, excluding the corpus callosum, and each cortical layer showed specific staining, with layers II/III and IV expressing CUX2/1, layers II/III, V, and VI expressing TBR1, and CTIP2 expressed in layers V and VI, and more prominently in the striatum (**STR**) (**Figs. 1b and 2b,d**). Strikingly, while these layer-defining transcription factors (CUX2/1, TBR1, CTIP2, and SATB2) were expressed at P21, their protein expression was considerably reduced compared to early postnatal stages (**Figs. 1b and 2b,d**).

**Fig. 2:**
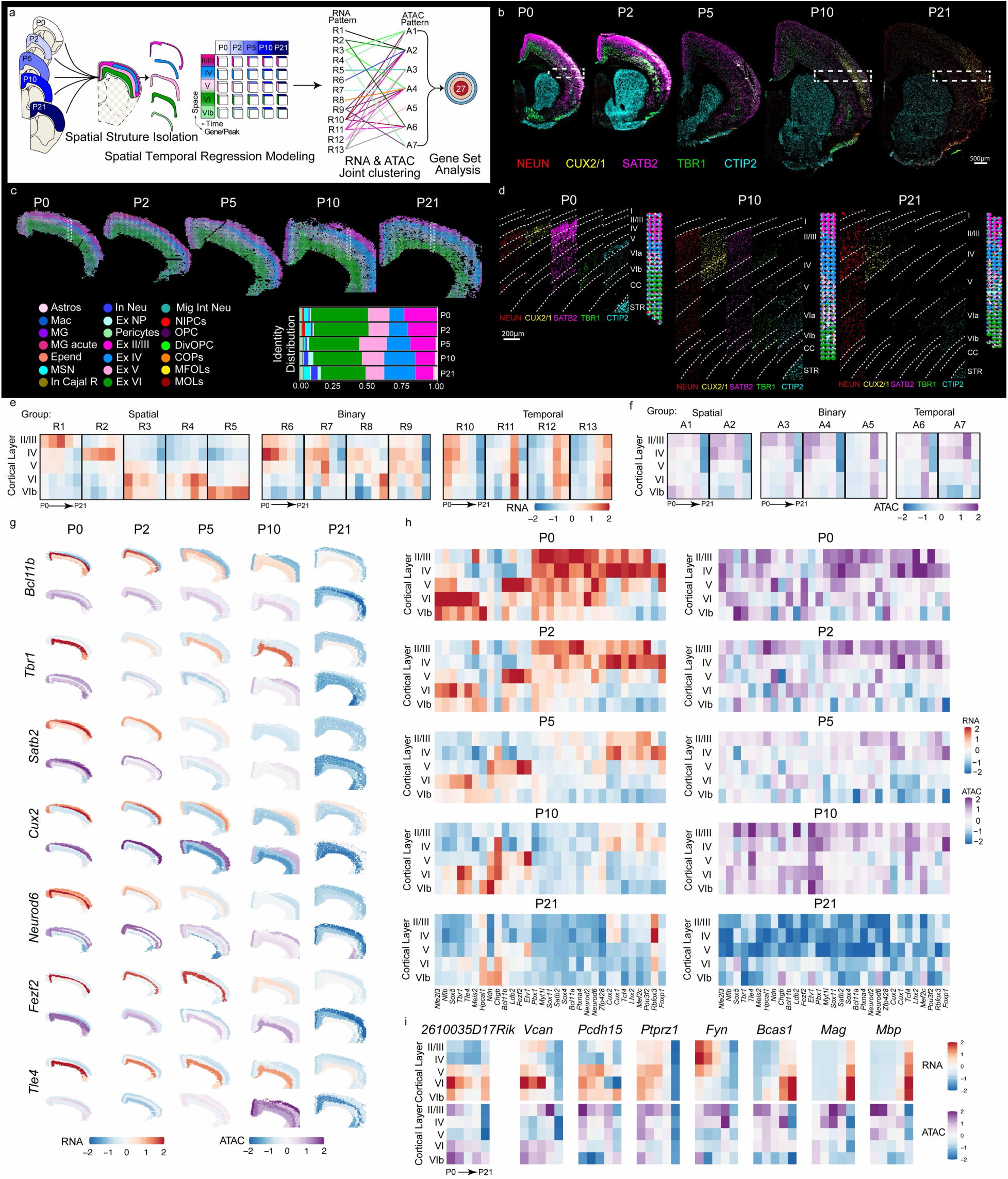
Spatial and temporal dynamics of the transcriptome and chromatin accessibility of the mouse brain cortical layers from P0 to P21. **a,** Schematic representation of spatial temporal regression model on mouse brain cortical layers. **b,** CODEX images of the mouse brain hemisphere from P0 to P21. Scale bar, 500 μm. **c,** Decomposition and pie chart visualization of different cell types in cortical layers. **d,** Delineation of cortical layers in the select region of interest indicated by dashed rectangles in **b** and **c**. **e,** Heatmaps of the 13 RNA clusters generated from the regression model. **f,** Heatmaps of the 7 ATAC clusters generated from the regression model. **g,** The RNA gene expression and ATAC GAS calculated on the basis of the regression model for specific genes. **h,** Heatmaps of the RNA gene expression (left) and ATAC GAS (right) calculated on the basis of the regression model for specific genes. **i,** Heatmaps of RNA gene expression (top) and ATAC GAS (bottom) for specific genes.

Delineation of the oligodendrocyte lineage was accomplished with OLIG2, the pan oligodendrocyte marker; in addition, with either PDGFRA to identify oligodendrocyte precursor cell (**OPC**) or myelin protein staining using MBP and MOG for mature oligodendrocyte (**MOL**). OLIG2 was, as expected, expressed in cells mainly in the corpus callosum but also throughout the brain. We found that the myelin components MOG were mostly absent from P0 to P5, only being robustly expressed at P10 (**Figs. 1b and 3b**), as previously reported^3^, which had opposing structural segregation to that of neuronal nuclei staining. Interestingly, however, the expression of MOG was limited to the lateral part of the corpus callosum at P10 and only spread throughout the entire corpus callosum at P21, suggesting a lateral to medial progression of myelination. Myelin in the cerebral cortex was also identified at P21 (**Fig. 3b and Extended Data Fig. 2b**).

**Fig. 3:**
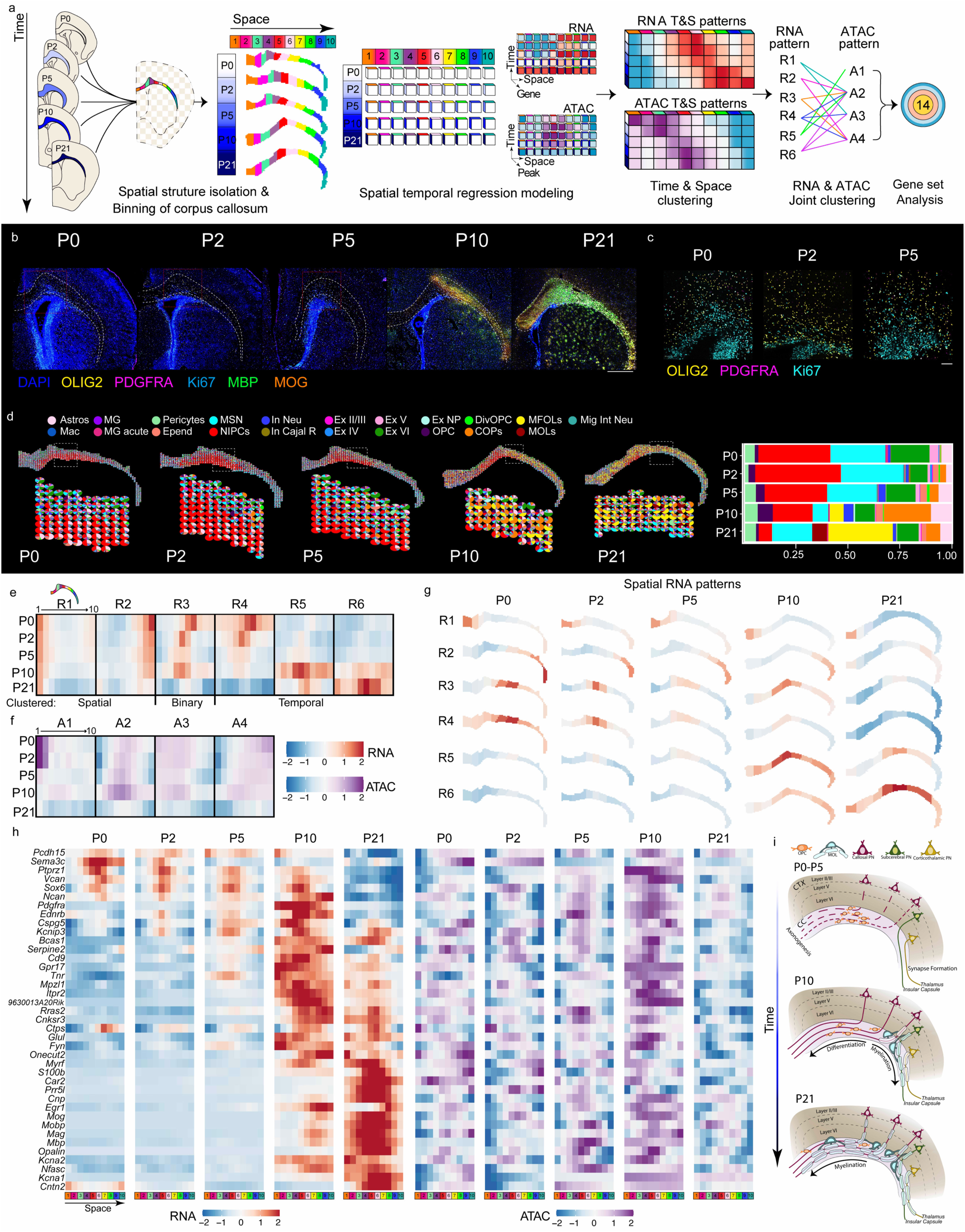
Spatiotemporal dynamics of corpus callosum during development and myelination. **a,** Schematic representation of spatial temporal regression model on mouse brain corpus callosum from P0 to P21. **b,** CODEX images of the mouse brain corpus callosum from P0 to P21. Scale bar, 1 mm. **c,** Magnified images of the regions shown as the red dashed rectangles in **b** from P0-P5, respectively, with staining of OLIG2, PDGFRA, and Ki67. **d,** Decomposition and pie chart visualization of cell types in corpus callosum. **e,** Heatmaps of the 6 RNA clusters generated from the regression model. **f,** Heatmaps of the 4 ATAC clusters generated from the regression model. **g,** Spatial RNA patterns for RNA clusters (R1-R6). **h,** The RNA gene expression calculated on the basis of the regression model for specific genes. **i,** Descriptive model summarizing how myelination progresses in the corpus callosum and cerebral cortex.

We also included markers for immune cells in CODEX; Innate immune cells, such as granulocytes/neutrophils (**GRAN/Nφ**) with LY6G; monocytes/macrophages (**Mono/Mϕ**) with IBA1, CD11b, and CD169; dendritic cells with CD11c and H2-Ab1 to more broadly identify cells with antigen presentation capacity. On the adaptive immune cell arm we incorporated CD3, CD4, and CD8, for pan T-cells, T-helper cells, and cytotoxic T-cells, respectively; and B cells using CD19 and CD45R (**Fig. 1a**). Overall, most immune cell subtypes were detected in development, but as expected were extremely sparse, being primarily localized to the meningeal compartment. CD169, however, was abundant in the meninges and present in the choroid plexus identifying a developmental counterpart to CNS resident CD169^+^ cells that increase after ischemia^20^ (**Extended Data Fig. 2c**).

The 19-plex CODEX panel for humans also included canonical markers for neurons, glia cell types, immune cell types, and proliferation (**Extended Data Fig. 3a**). CODEX imaging of the human brain V1 cortex revealed distinct arealization. Similar to our observations in the mouse postnatal brain, we observed a notable decrease in the expression of SATB2, CTIP2, and TBR1 at infancy compared to early developmental stages. Notably, myelination marked by MBP expression began to intensify during infancy, consistent with prior reports (**Extended Data Fig. 3b**)^21^. Overall, our CODEX profiling, by providing a comprehensive protein benchmark at single-cell resolution of developmental mouse and human brain at different ages, uncovered dynamic patterns of layer-defining transcription factors and myelination.

### DBiT-based spatial tri-omic sequencing to simultaneously co-profile epigenome, transcriptome, and protein in the brain tissue sections

To further investigate the molecular mechanisms across all the layers of the central dogma underlying the spatial dynamics unveiled by CODEX, we developed the all-encompassing spatial assay that allows co-profiling of epigenome, transcriptome, and a large panel of proteins simultaneously in the same tissue section. Spatial <u>A</u>TAC–<u>R</u>NA–<u>P</u>rotein-seq (**DBiT ARP-seq**) is shown schematically in **Fig. 1a** and **Extended Fig. 1a**. A frozen tissue section was fixed with formaldehyde, incubated with a cocktail of 136 antibody-derived DNA tags (ADTs) for mouse (BioLegend mouse panel with spiked intra-cellular proteins, **Extended Data Table 4**) or 163 ADTs for human (BioLegend human panel, **Extended Data Table 5**), and then treated with the Tn5 transposition complex pre-loaded with a DNA adapter containing a universal ligation linker, that can be inserted into the transposase accessible genomic DNA loci. The same tissue section was then incubated with a biotinylated DNA adapter, also containing a universal ligation linker and a poly-T sequence, which binds to the poly-A tail of mRNAs and ADTs to initiate reverse transcription (RT) directly in tissue. Subsequently, two microfluidic channel array chips with perpendicular channels were applied to the tissue section to sequentially introduce spatial barcodes Ai (i = 1−100) and Bj (j = 1−100), creating a 2D grid of spatially barcoded tissue pixels (pixel size 20 μm) each uniquely defined by combinations of barcodes Ai and Bj, totaling 10,000 barcoded pixels. The procedure was concluded with the release of barcoded cDNAs (from both mRNAs and ADTs) and genomic DNA (gDNA) fragments following reverse crosslinking, where cDNAs are enriched using streptavidin beads and gDNA fragments are collected in the supernatant. Separate libraries for gDNA and cDNAs were then constructed for next-generation sequencing.

The mouse ADT protein panel encompasses 119 antibodies targeting cell surface antigens, 9 isotype control antibodies specific to immune cells, and 8 self-conjugated antibodies for intracellular markers representative of canonical mouse brain cell types (**Extended Data Table 4**). Similarly, the human ADT panel includes 154 antibodies against unique cell surface antigens, covering key lineage antigens, complemented by 9 isotype control antibodies for immune cell profiling (**Extended Data Table 5**).

For the spatial ATAC modality, across all samples from mouse brains from development stages (P0-P21), the LPC model at 5 DPL and 21 DPL, and human brain V1 region from the second trimester, third trimester, and infancy, we observed a median of 14,489 unique fragments per pixel. Among these, 16% were enriched at transcription start sites (TSS) regions and 15% located in peaks (**Extended Data Figs. 4a, 5a**). For the RNA portion, a total of 23,824 genes were detected with an average of 1,108 genes and 2,066 unique molecular identifiers (UMIs) per pixel (**Extended Data Fig. 5a**). For the proteins, the average protein count per pixel is 82 and the protein UMI account per pixel is 863.

We also developed spatial tri-omic profiling that measures genome-wide histone modification in conjunction with transcriptome and proteins. Spatial <u>C</u>UT&<u>T</u>ag–<u>R</u>NA–<u>P</u>rotein-seq (**DBiT CTRP-seq**) was conducted similarly, by applying instead an antibody against H3K27me3 to the tissue section and then using protein A tethered Tn5-DNA complex to perform co-assay of cleavage under targets and tagmentation (**CUT&Tag**)^22,23^ (**Fig. 1a and Extended Data Fig. 1b**). We performed DBiT CTRP-seq for H3K27me3 on 5 DPL and 21 DPL mouse brains. We obtained a median of 9,102 unique fragments per pixel, of which 10% of fragments overlapped with TSS regions, and 12% were located in peaks (**Extended Data Fig. 4a,5a**). For the RNA data, a total of 23,397 genes were detected with an average of 1,318 genes per pixel and 2,392 UMIs per pixel (**Extended Data Figs. 4a, 5a**). For the proteins, the average protein count per pixel is 88 and the protein UMI account per pixel is 793.

The insert size distributions of chromatin accessibility (DBiT ARP-seq) and histone modification (DBiT CTRP-seq, H3K27me3) fragments were consistent with the captured nucleosomal fragments in all tissues (**Extended Data Fig. 4b**). The correlation analysis between replicates showed high reproducibility (r = 0.99 for ATAC, r=0.98 for RNA, and r=0.99 for protein in DBiT ARP-seq; r=0.98 for CUT&Tag (H3K27me3), r=0.95 for RNA, and r=0.99 for protein in DBiT CTRP-seq, r stands for Pearson correlation coefficient, **Extended Data Fig. 5b,c**).

### DBiT ARP-seq allows molecular deconvolution of the development of the mouse and human cortical layers and corpus callosum

To investigate the spatial dynamics of mammalian cortical and white matter development at the molecular level, we utilized DBiT ARP-seq on adjacent tissue sections processed with CODEX. After applying DBiT ARP-seq to postnatal mouse brain sections, we clustered RNA and ATAC data separately, integrating across all timepoints. All the samples integrated well, with the coronally sectioned mouse brains identified by 22 major RNA (R0-R21) and 15 major ATAC clusters (A0-A14) (**Fig. 1e-h**). Notably, after mapping cell states defined by scRNA-seq^24–26^ to our RNA data (**Extended Data Figs. 6 and 7**), we found the cluster R16 localized to the corpus callosum, identified as myelin-forming oligodendrocytes (MFOLs) (**Fig. 1m and Extended Data Fig. 6b**), was only represented in the P10 and P21 samples (**Fig. 1e-h**), which is consistent with our CODEX data **(Fig. 3b)**. Integrating single-cell ATAC-seq mouse brain atlas data^27^ with our P10 ATAC-seq data also identified all major cell types (**Extended Data Fig. 7**). Label transfer was then applied to assign cell types to specific spatial locations based on the epigenetic states that may govern cell type formation. All dominant cell types identified through ATAC-seq are consistent with the results from RNA-seq label transfer (**Extended Data Fig. 7b-e**). For better delineation of brain sections, we further integrated our RNA and ATAC data using SpatialGlue^28^, leading to 18 refined spatial domains (D1-D18) (**Fig. 1i and Extended Data Fig. 8a,b**). Most of the brain regions retained the same delineation during mouse brain development from P0 to P21 (**Fig. 1i**). Spatial distribution of these domains aligned with tissue histology from Allen brain atlas^29^ and provided better arealization when compared to CODEX (**Fig. 1b and Extended Data Fig. 2**).

Consistent with the CODEX analysis, our DBiT ARP-seq data also revealed that MBP and MOG proteins, and chromatin accessibility and RNA expression of *Mbp* and *Mog*, were induced at P10 and P21 but not at P0 (**Fig. 1j-l and Extended Data Figs. 6c-e and 8c,d**). The protein data from DBiT ARP-seq ADTs also showed general concordance in the positional signal for CTIP2, CUX2/1, NEUN, SATB2, TBR1 as compared with RNA gene expression, ATAC gene activity score (**GAS**), and the CODEX imaging (**Extended Data Figs. 7a and 9**). Nevertheless, in contrast to CODEX, the integrated spatial domains led to the division of cortical layers II/III and V, subdividing the cortical areas between primary and secondary motor and eventually anterior cingulate at each layer^29^, with medial D4 and D14 clusters segregating from clusters D9 (layer II/III) and D2 (layer V), respectively (**Fig. 1i**). While the striatum was characterized by markers of medium spiny neurons such as *Bcl11b* (clusters D5, D12, and D17) (**Extended Data Fig. 9**), during the developmental process, cluster D17 was gradually substituted with cluster D5, where we found that the expression and ATAC gene activity score of *Foxp4* (at cluster D17), a transcription factor known to regulate morphogenesis, disappeared gradually (**Fig. 1j,k and Extended Data Fig. 6d,e**)^30–32^. Furthermore, the expression change of *Foxp4* in the postnatal striatum is consistent with previous studies showing *Foxp1* and *Foxp2* retaining dominance in that region postnatally, both of which exhibit high expression in the striatum in our data set^32,33^ (**Fig. 1j and Extended Data Figs. 6d, 8c**). While our CODEX identified sparse immune cell labeling with a broader ADT panel, we were able to detect more immune cell markers for monocytes/macrophages/microglia (CD11b, CD169, CD68, CD86, and CD26), dendritic cells (CD11c) and NK cells (CD49b). The abundance of some immune populations was quite high indicating a possible redundancy of marker expression with resident central nervous system (CNS) cells (**Extended Data Fig. 7a**).

DBiT ARP-seq on human brain V1 region tissue sections, adjacent to those analyzed with CODEX, identified 16 RNA and 9 ATAC clusters (**Extended Data Fig. 3c,d**). This data suggests that transcriptional profiles prominently define cell types during the prenatal and neonatal stages of human brain development. The spatial regions identified demonstrated consistency with the MERFISH data obtained from adjacent tissue sections^34^. For all the samples, RNA clusters R1, R2, R9 (second trimester), R6, R9 (third trimester), and R5, R3, R4, R10 (infancy) represent cortical layers, with cortical neuron marker *NCAM1*, which is involved in neurogenesis. Additionally, the ADT protein expression of CD56 (protein for *NCAM1*) correlated with *Ncam1* RNA expression and chromatin accessibility. *RBFOX3* expression was primarily noted in the cortical layers, and *MBP* expression began to dominate starting from the third trimester. All these results are consistent with observations from CODEX (**Extended Data Fig. 3e,f**). The ATAC data also revealed unique clusters with marker genes enriched in cortical layers and WM, including *NCAM1*, *RBFOX3*, and *MBP* (**Extended Data Fig. 3e,f**). Thus, our spatial tri-omic DBiT ARP-seq and CODEX in the mouse and human brain yields a comprehensive dataset with the important features of spatial conservation and temporal evolution verified across all three layers of the central dogma.

### Temporal persistence of chromatin accessibility for cortical layer-defining transcription factors

Our CODEX and DBiT ARP-seq analysis allowed the identification of the cortical layer-defining transcription factor expression at the protein (CODEX and ADTs), RNA, and chromatin accessibility levels. To investigate the noted temporal and spatial variations (**Figs. 1b, 2b-d and 3b**) at the chromatin accessibility and transcriptome levels among cortical layers, we developed a computational framework to systematically examine RNA and ATAC patterns crossing three dimensions, i.e. space, time and modality (**Fig. 2a**, see Methods). The framework starts with applying generalized additive regression to model the spatial (cortical layers II/III, IV, V, VI, and VIb), temporal (P0-P21) effects and their interaction on gene activity score for ATAC and gene expression for RNA. We then undertake a two-step procedure to categorize patterns of genes that exhibit significant spatial/temporal changes in either RNA or ATAC. First, we perform joint clustering on the concatenated RNA and ATAC data to simultaneously capture the spatiotemporal patterns across both modalities. Subsequently, we summarize these patterns based on their similarities in RNA and ATAC profiles, respectively, and combine clusters that exhibit similar patterns. Using this framework, we identified 13 distinct RNA clusters and 7 ATAC patterns, resulting in 27 unique combinatorial clusters that display varied patterns of changes across time and space (**Fig. 2a,e,f, Extended Data Fig. 10 and Extended Data Table 7**).

Among the 13 RNA clusters identified, we categorized them into three major groups based on similar expression patterns across space and/or time (**Fig. 2e and Extended Data Fig. 10a**). R1-R5 had an expression that was predominantly related to space, and therefore we observed high RNA expression restrained to specific layers, in particular *Bcl11b* (R3) and *Tbr1* (R3) (**Fig. 2e,g,h, Extended Data Figs. 9-11 and Extended Data Table 7**). In contrast, for R10-R13 clusters, time was the main driver of RNA expression changes, with the temporal variation of expression being consistent across a subset of layers. *Satb2* (R10) presented such a pattern (**Fig. 2e,g,h, Extended Data 9-11**). The third group (R6-R9) was binary with the influence of both time and space. For instance, *Cux1*&*2* are present in the binary RNA cluster R6 (**Fig. 2e,g,h, Extended Data Figs. 9-11 and Extended Data Table 7**).

The ATAC gene activity score patterns exhibited a more diffuse spatial/temporal distribution, and the identified patterns did not always reflect the variations observed in the RNA clusters (**Fig. 2e,f**). While for a proportion of cortical-layer specific transcription factors there was an alignment between RNA expression and chromatin accessibility (*Cux1*, *Cux2*, *Foxp1*, *Lhx2*, *Mef2c*, and *Pou3f2*), we observed a misalignment for a subset of key transcription factors and the different layers (**Fig. 2g,h and Extended Data Fig. 11**). Specific subsets of transcription factors presented chromatin accessibility trailing in time and/or more diffused in space, when compared to their RNA expression. ATAC signal for *Bcl11a*, *Etv1*, *Meis2*, *Myt1l*, *Neurod2*, *Neurod6*, *Pbx1*, *Satb2*, *Sox4*, and *Sox11* trailed in time compared to RNA (**Fig. 2g,h and Extended Data Fig. 11**). Previous single-cell RNA-seq and ATAC-seq on projection neurons (**PN**), including subcerebral (**SCPN;** *Bcl11b*, *Fezf2*, *Ldb2*, *Sox5*), corticothalamic (**CTPN**; *Tle4*, *Sox5*, *Tbr1*, *Ndn*, *Tcf4*, *Hpcal1*, *Zfp428*, *Chgb*), and callosal projection (**CPN**; *Cux1*, *Cux2*, *Plxna4*, *Satb2*) during embryonic development revealed that chromatin accessibility precedes gene expression in certain cases^1^. We mapped known markers for PNs to determine if their spatial segregation was accurate. Each subclass of PN had marker enrichment in 1-4 RNA clusters. Higher RNA expression in the upper cortical layer clusters (R2, R6, and R10) was enriched for callosal CPN markers, with the most residing in cluster R10. The deeper layers, with a higher RNA expression, contained subcerebral SCPN markers populating almost exclusively to R3, and corticothalamic CTPN that had a little broader cluster representation in R3, R5, R9, and R13 (**Fig. 2e-h, Extended Data Fig. 11 and Extended Data Table 7**). In contrast to embryonic development^1^, we did not observe any cases of chromatin accessibility leading gene expression for all marker genes of SCPN, CTPN, and CPN in our postnatal dataset (**Fig. 2e-h and Extended Data Fig. 11**). Furthermore, we observed a cohort of transcription factors, such as *Neurod2* and *6,* for which the chromatin accessibility persisted longer than RNA (**Fig. 2g and Extended Data Fig. 11**). Thus, while epigenetic priming occurs during embryonic development upon establishment of specific expression of genes characteristic of different cortical layers^6^, our data suggests that in post-natal development we observe a decrease in the expression of these genes (both at RNA and protein level), which is not accompanied by reduction of ATAC signal, suggesting that these genes could retain epigenetic memory of the previous transcriptional state of an earlier stage, despite the marked decrease in transcriptional output. Nevertheless, this epigenetic trailing subsides at P21, when both RNA and protein expressions are significantly reduced.

### Spatial spreading of chromatin accessibility for cortical layer-defining transcription factors

Cortical-layer specific transcription factors such as *Bcl11b*, *Fezf2*, *Ldb2*, *Nfe2l3*, *Nfib*, *Rbfox3*, *Sox5*, *Tbr1*, and *Tle4* were found to present chromatin accessibility spreading across layers (**Fig. 2g,h and Extended Data Fig. 11**), suggesting the possibility of inherent epigenetic plasticity across different cortical layers. For example, *Fezf2* and *Tle4* are quite restricted at the RNA expression level to layer V and VI, respectively (**Fig. 2g and Extended Data Fig. 11a**). However, their gene accessibility score extends to layer VI and V, respectively (**Fig. 2g,h and Extended Data Fig. 11a**). The dynamics between these two factors has been recently documented in which the CTPNs in *Tle4* knockout mice are not distinct from their corticofugal counterpart, SCPN. Therefore, coverage of ATAC suggests plasticity even though *Tle4* is a transcriptional repressor of *Fezf2*^35^*. Tbr1* is another example of chromatin accessibility spreading across layers amidst defined RNA expression. Between P0-P21, the RNA expression for *Tbr1* is maintained in layer VI while the chromatin accessibility extends to layer II/III. The accessibility we observe for *Tbr1* makes sense in the context of its diverse functions across projection neurons and differential expression across time^36^. We have previously shown deposition of the repressive histone mark H3K27me3 at the TSS of genes not expressed in specific cortical layers, and its absence in genes expressed in those layers^15,37^. Thus, despite the spatial diffusion of chromatin accessibility at postnatal stages in cortical layers, Polycomb-mediated repression mechanisms might prevent ectopic expression of layer-specific genes, which warrants further investigation.

### Spatial and temporal patterns of axonal maturation and myelination in the postnatal cortical layers

In order to explore signaling pathway dynamics over space and time in the postnatal cortical layers, we performed gene ontology (GO) analysis on each of the 27 combined RNA/ATAC clustered patterns (**Extended Data Figs. 12, 13**). As expected, we found that general GO terms affiliated with neuronal biology terms across the clustered patterns are associated with callosal CPN, corticothalamic CTPN, and subcerebral SCPN. However, other terms were more specified to specific projection neuron subtypes. Glutamatergic signaling was found only in three clusters of R3-A4 (CTPN & SCPN associated), R5-A3 (CTPN associated), and R9-A7 (CTPN associated). Moreover, other neurotransmission terms (calcium ion-regulated exocytosis of neurotransmitter and regulation of neurotransmitter transport) were also identified in the corticothalamic CTPN-associated R13-A3. Callosal CPN-associated clusters had the terms semaphorin-plexin signaling, axonogenesis, and neuron migration in R10-A2/A4 (CPN associated), which was the predominant cluster for guidance cues. Taken together, all clusters were undergoing general developmental changes, callosal CPN seemed to be less mature due to the ongoing axonal guidance programs, while corticothalamic CTPN and subcerebral SCPN were more mature with features of exocytosis, and glutamatergic synaptic transmission already underway. The maturity regarding postnatal development is consistent with embryonic development order since corticothalamic CTPN are the first to emerge at E11.5-13.5, followed by subcerebral SCPN at E12.5-14.5 and finally the callosal CPN at approximately E15.5-E17.5^1^.

Interestingly, in addition to projection neuron patterning, we also identified clusters exhibiting GOs related to oligodendrogenesis (**Extended Data Figs. 12, 13 and Extended Table 7**). RNA expression linked to OPCs and MOLs was identified in R3 (**OPC/COP/NFOL**; *Vcan*, *Spon1*, *Pcdh15, 2610035D17Rik*), R9 (**OPC/COP**; *Ptprz1*, *Serpine2*, *Grin3a*), R10 (**OPC/COP/NFOL**; *Sox6*, *Fyn*, *Fam107b*, *Frmd4a*) and R12 (**OPC**; *Bcas1*, *Sept8,* **MOL**; *Nrxn3*, *Nrgn*, *S100b*)^13,38,39^ (**Extended Data Fig. 14 and Extended Table 7**). NMDA receptor subunits indicating primed OPCs were also segregated to R9 (*Grin2d*, *Grin3a)* and R12 (*Grin2a*)^40^ (**Extended Data Fig. 14 and Extended Table 7**). Since clusters R3 and R9 also represent corticothalamic CTPN and subcerebral SCPN, our data suggests an overlap between the time and space of OPCs and corticofugal projection neuron maturation. Myelin-associated genes identified across gene sets were entirely clustered to R13-A3 (*Mbp*, *Mag*, *Mobp*), which is the final temporal program to arise beginning at P10 and increasing at P21 and is associated with CTPN (**Fig. 2i and Extended Data Fig. 14b**). Interestingly, there was a greater intensity of the spatial segregation of myelin-associated gene expression, where layer VI had the highest levels for all the myelin genes (**Fig. 2i and Extended Data Fig. 14b**). The distinction between these clustered/patterned programs in time but not space provides a window of opportunity to unravel mechanisms involving preferential myelination and differential patterning in relation to the PN subsets. Thus, our data indicates that OPCs follow a similar spatial and temporal trajectory to that of corticothalamic CTPN and subcerebral SCPN in early postnatal development, while at P10 and P21, the spatial myelination takes over which is seen more heavily in the deep layers where the SCPN and CTPN heavily populate.

### Priming of chromatin accessibility of myelin genes across the corpus callosum during the myelination window

The timing of myelination in the developing mouse brain has been well documented to initiate around P10^3^. A possible lateral to medial progression of oligodendrocyte differentiation at these stages was previously observed by in situ sequencing^41^. The expression of MBP and MOG was limited to the lateral part of the corpus callosum at P10 and was only spreading throughout at P21, suggesting a lateral to medial progression of myelination (**Fig. 3b**). We utilized DBiT ARP-seq developmental data set (P0-P21) to further investigate at a spatial and molecular level postnatal myelination, by coupling the spatial framework of tissue architecture with our tri-omic sequencing approach (**Fig. 3a**). ADT protein data from DBiT ARP-seq confirmed that MBP and MOG proteins were expressed at P10 and P21 but not at P0, whereas OLIG2 was expressed at P0 (**Fig. 1l and Extended Data Figs. 6c, 7a**). *Mbp* and *Mog* presented chromatin accessibility and expression at the lateral part of the corpus callosum (cluster D6) at P10 and became abundant in the whole corpus callosum at P21 (**Fig. 1j,k and Extended Data Figs. 6d,e, 8c**). Before P10, the corpus callosum was characterized by markers of oligodendrocyte precursor cells (OPC) such as *Olig2* and *Pdgfra* (**Fig. 3d and Extended Data Fig. 8c**).

Interestingly, we observed not only concordance between the transcription of oligodendroglia genes and their chromatin accessibility, but also chromatin priming at P0-P5 in regions of the corpus callosum where RNA expression is observed to occur at P10 (**Fig. 3h**). At P21, while transcription persists for some myelin genes, the chromatin accessibility was already reduced (**Fig. 3h**). Thus, the spatial/temporal chromatin accessibility dynamics of myelination in the corpus callosum at postnatal stages is different from the dynamics for cortical layer specific transcription factors, which presents priming at embryonic stages^1^, but trailing at postnatal stages (**Fig. 2**).

### Spatial dynamics of oligodendroglia differentiation and myelination in the post-natal corpus callosum

To determine if there is indeed a lateral to medial progression in myelination, we isolated the corpus callosum (P0-P21) and computationally partitioned it into 10 separate regions spanning medial to lateral in their positional order (**Fig. 3a**). Like the separate cortical layer analysis in cortical layers, we took each segment, separately, to simultaneously determine the RNA expression and chromatin accessibility. Afterwards, we rebuilt the spatial architecture of the 10 segments to construct RNA and ATAC patterns and derive their respective gene set lists enabling us to map the changes over the entire space. In the final step, we curated the RNA and ATAC patterns. This approach enabled us to simultaneously analyze the corpus callosum (**CC**) development through space and time. We mapped 6 RNA and 4 ATAC clustered patterns, and 14 combined (**Fig. 3e,f and Extended Data Fig. 15**). As in the cortical layers, clusters could be categorized into spatial, binary, and temporal trends (**Fig. 3e**), and ATAC patterns were less defined and were influenced by both time and space (**Fig. 3f**). We observed clusters that exhibited high RNA expression early in either the medial (R1) or lateral (R2) region which was maintained from P0 to P21. Two other clusters had early high expression centrally, that subsequently favored a more medial (R4) or lateral (R3) region (**Fig. 3e,g and Extended Data Fig. 15**). Across all the time points, the medial-lateral domain border corresponded with the eventual early MOG protein staining that we observed in the CODEX and ADT at P10 (except for R1) (**Figs. 3b and Extended Data Fig. 6c, 15**). Therefore, our initial hypothesis of lateral to medial myelination seemed to perhaps contribute but not be the main factor regulating the myelination of axons.

The general landscape of the 14 different concerted RNA/ATAC programs and the regional differences were further identified by GO term analysis (**Extended Data Fig. 16**). We found that only one group (R6-A2) was identified with the terms associated with myelination. Myelin-associated genes and maturation markers were found in this set (*Cnp*, *Egr1*, *Mog*, *Mobp*, *Mag*, *Mbp*, and *Opalin*) (**Fig. 3h, Extended Data Figs. 16-18 and Extended Data Table 8**)^39^. This cluster was not only producing myelin but likely myelination was occurring since we identified genes involved in the node of Ranvier including neurofascin, Kv1.1/1.2, and contactin (*Nfasc*, *Kcna1*, *Kcna2*, and *Cntn2*)^42–45^ (**Fig. 3h, Extended Data Figs. 16-18 and Extended Data Table 8**). Interestingly, R3-A2 and R5-A2 presented GO terms of glial migration were classified (**Extended Data Figs. 16, 17**). These clusters presented stronger connections with specific oligodendroglia progenitors cell types (**R3-OPC/COP**; *Ptprz1, Vcan*, *Sox6, Ncan, Pcdh15, Sema3c* and **R5-OPC/COP**; *Pdgfra, Ednrb, Cspg5, Kcnip3, Bcas1, Serpine2, Cd9*, *Gpr17*) (**Fig. 3h, Extended Data Figs. 17-19 and Extended Data Table 8**). Shared genes between committed oligodendrocyte precursors (**COPs**), newly formed oligodendrocytes (**NFOLs**), myelin forming oligodendrocytes (**MFOLs**) were primarily in R5 (**COP/NFOL/MFOL**; *Tnr*, *Mpzl1*, *Itpr2*, *9630013A20Rik*, **NFOL**; *Rras2*, *Cnksr3*, and **MFOL/MOL**; *Ctps*, *Glul*) (**Fig. 3h and Extended Data Fig. 19**). Thus, R5 is likely to represent an overall more mature population of oligodendrocytes. Cluster R6-A4 also contained a mix of COPs and MOL specific subtype genes (*Fyn, Onecut2, Myrf, S100b, Car2*, and *Prr5l),* placing this population even more differentiated than R5 but less than R6-A2 (**Fig. 3h, Extended Data Figs. 17, 20 and Extended Data Table 8**). Thus, in contrast of the CODEX protein analysis, DBiT-ARP suggests a medial to lateral maturation hierarchy of R3-A2 >> R5-A2 >> R6-A4 and R6-A2.

R3 to R5 had observably high expression just below the cingulum bundle (regional space bin 3-6; central CC for the individual hemisphere) but then reduced expression in that region (**Fig. 3e,g**). By mapping this population to the cell type pie charts, we found that the cluster trend was likely due to remnant neural progenitors like cells and OPCs (**Fig. 3d**). There is a proportion of these cells in the P0 and P2 tissue sections that are co-labeled with OLIG2 and PDGFRA (**Fig. 3b,c**). Thus, our data suggests that an early response of OPCs/COPs begins at P0 that modestly favors the lateral CC (R3), then at P5-P10 the expression of NFOLs and MFOLs can be seen with a high lateral preference so that at P10 the myelination begins and moves medially by P21. Interestingly, a full medial shift (to the CC midline) of the OPCs/COPs begins between P5-P10 and ends by P21 with minimal expression for each gene, while the NFOLs occur between P10-P21 preceding the myelination in time and space (**Fig. 3h**). In summary we observe a timed and sequential bidirectional myelination process occurring from the central CC where the OPC/COPs enter from the postnatal wave to first migrate and differentiate laterally followed by the second phase of migration and differentiation in the medial direction (**Fig. 3i**).

### Layer specific projection neurons axonogenesis and synaptogenesis time may orchestrate myelination

Interestingly, while interrogating the GO terms for the corpus callosum, we found that the MOL and myelin-associated group (R6) was associated with many synaptic transmission terms, while the OPC/COP (R3) group was demarcated with earlier neuron programs like neuron migration, neuron projection regulation, and dendritic spine development (**Extended Data Figs. 16, 17**). The neuronal programs were segregated in space where the more mature had expression on the lateral end of the CC, and the developmental neurons had medial expression (**Fig. 2e,g**). Surprisingly, by examining these two clusters, we found cortical layer marker expression of *Bcl11b* (Layer V; SCPN) in R2, and *Cux1* and *Lhx2* (Layer II/III; CPN) in R4 (**Extended Data Fig. 20 and Extended Data Table 8**). These markers were consistent with cortical layer data and might originate from their RNA transport into the axonal tracts. Other genes with medial expression that have been linked to the callosal CPNs include *Foxg1, Plxna4* and *Epha4*, which are involved in the axonal guidance of this projection neuron subtype (**Extended Data Fig. 20**)^46,47^. Corticothalamic CTPNs that have been experimentally deleted for *Sema3e* have postnatal deficits with pathfinding^48^. Consistent with the axonal positioning for CTPNs, we found *Sema3e* expression constrained to the lateral end of the CC. Both *Plxna1* and *Plxnd1* have been associated with projection neurons and we see an interesting expression pattern in which it begins laterally and moves medially through development (**Extended Data Fig. 20**)^49^. Taken together, in the regions of the corpus callosum, where myelination initiates, we have subcerebral SCPN marker expression along with expression of genes associated with neural transmission. In the medial region that gets myelinated later, we see gene expression associated with corticothalamic CPN that are still undergoing axonogenesis. When considering the inside-out temporal development of PNs, subcerebral SCPNs are defined before corticothalamic CPNs. Furthermore, the establishment of their axons is also separated in time in which corticofugal PNs project embryonically starting around E14.5 and CPNs project postnatally and continue refining through P8^48,50,51^. Therefore, cues from the neurons, their maturation, and their directionality of their axon might orchestrate the OPC migration to the MOL myelination process (**Fig. 3i**).

### Spatial tri-omic mapping of neuroinflammation and remyelination using a focal lesion mouse model of MS

OPCs are thought to contribute to remyelination in the context of multiple sclerosis^52^. To study the process of remyelination yielding the ability to interrogate developmental and reparative programs simultaneously, we injected 1% lysolecithin (**LPC**) into the mouse corpus callosum (unilaterally and ventral to cingulum bundle) at 85 days and conducted CODEX, DBiT ARP-seq, and DBiT CTRP-seq (targeting H3K27me3, repressive loci) on coronal sections (**Fig. 4**).

**Fig. 4:**
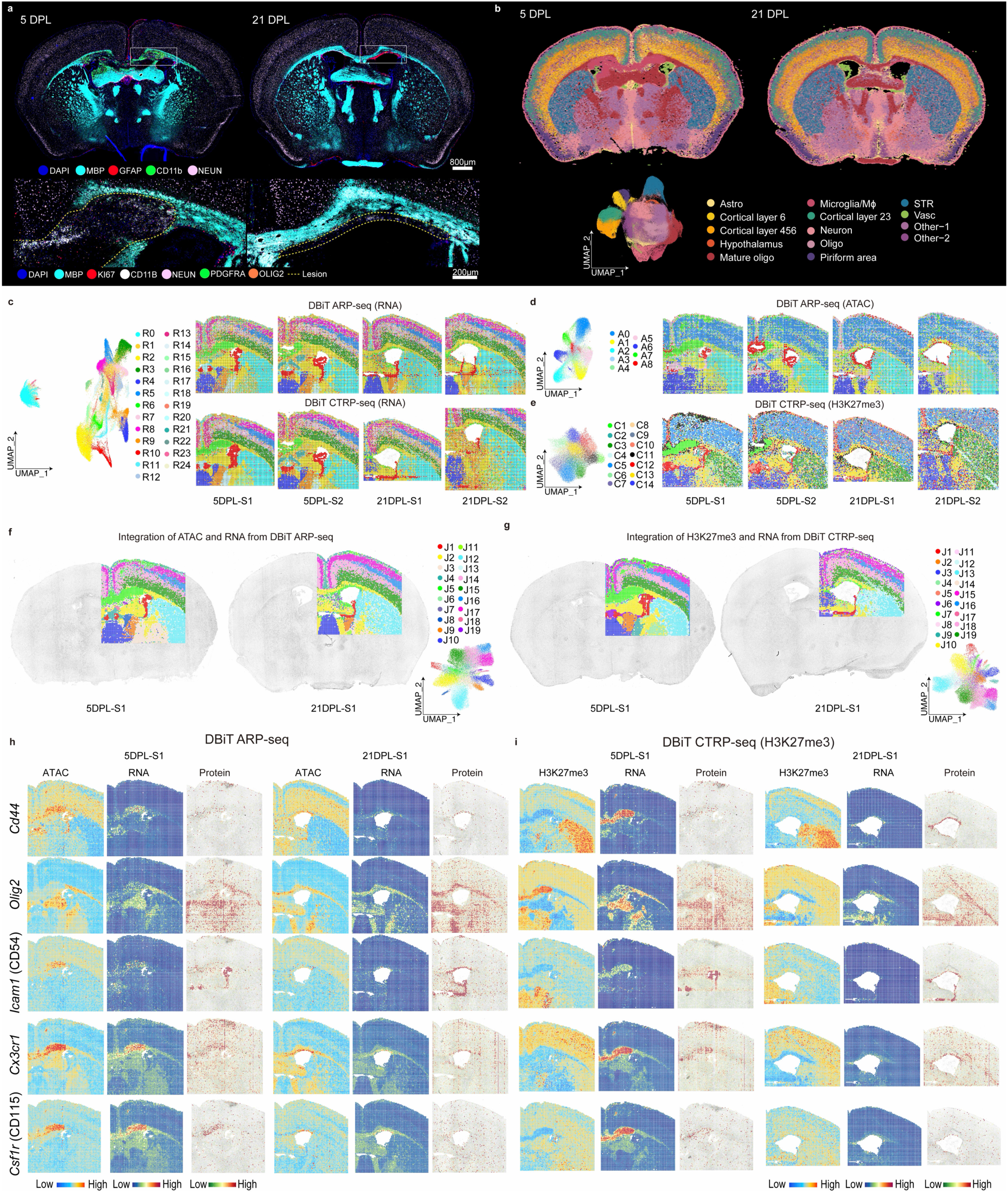
Spatial epigenome–transcriptome–proteome mapping of the LPC mouse model brains at 5 DPL and 21 DPL. **a,** CODEX images of whole mouse brains at 5 DPL and 21 DPL. The images at the bottom are magnified views of the regions indicated by the white dashed rectangles. **b,** Seurat clustering of the CODEX images in **a**. **c,** The UMAPs and spatial distribution of RNA clusters in both DBiT ARP-seq (upper) and DBiT CTRP-seq (H3K27me3) (bottom) for the LPC mouse samples at 5 DPL and 21 DPL (including the replicates). **d-e,** UMAPs and spatial distribution of ATAC clusters in DBiT ARP-seq (**d**), as well as H3K27me3 CUT&Tag clusters in DBiT CTRP-seq (**e**) for LPC mouse samples at 5 DPL and 21 DPL (including the replicates). **f,** Integration of RNA and ATAC data in DBiT ARP-seq. **g,** Integration of H3K27me3 and RNA data in DBiT CTRP-seq. **h-i,** Spatial mapping of gene expression, GAS, CSS, and ADT protein expression for *Cd44*, *Olig2*, *Icam1*, *Cx3cr1*, and *Csf1r* in both DBiT ARP-seq (**h**) and DBiT CTRP-seq (**i**).

CODEX provided a detailed single-cell resolution and morphological visualization of the lesion evolution at 5 days post-lesion (**DPL**) and repair at 21 DPL in the coronal brain sections (**Fig. 4a**). We performed unsupervised clustering of protein expression from 444,947 of cells across 4 coronal samples, and the cell types were assigned to each cluster, accordingly, based on the marker expression and the Allen Brain Atlas^29^ (**Fig. 4b and Extended Data Fig. 23**). While most of the cell types remained unaltered between 5 DPL and 21 DPL, MOLs and myelin were lost at the lesion site. Astrocytosis was prominent along the injection track and perilesional and there was observable macrophage/microglia accumulation in the lesion at 5 DPL. In contrast, by 21 DPL, the lesion site was much smaller, and successful remyelination was observed in the CC, with significant reduction of the astrocytic and macrophage/microglial presence (**Fig. 4a and Extended Data Fig. 23**). At 5 DPL there was already a response of OPCs to the insult (OLIG2 and PDGFRA), increased proliferative activity (Ki67), and the presence of granular cells (**Fig. 4a**), highlighting inflammation and the beginnings of the regenerative processes. At 21 DPL, the resolved lesion was marked with a reduction of CD11b+ macrophages/microglia and a corresponding resolution of the inflammatory response. The OLIG2 and PDGFRA positive cells regenerated and MOLs remyelinated most of the lesion area (MBP staining, **Fig. 4a**).

DBiT ARP-seq and DBiT CTRP-seq (H3K27me3) were performed on the adjacent tissue sections to that of CODEX. For both 5 DPL and 21 DPL, we identified ATAC and RNA specific clusters for DBiT ARP-seq (ATAC: 13 and RNA: 25) and DBiT CTRP-seq (H3K27me3: 9 and RNA: 25) (**Fig. 4c,d**). All modalities (RNA, ATAC, and CUT&Tag) could delineate the specific regions of mouse coronal brains and agreed well with anatomical annotation defined by Allen Brain Atlas^29^. We then conducted integration of ATAC and RNA, as well as H3K27me3 and RNA, respectively, for better delineation of the mouse brains (**Fig. 4f,g and Extended Data Fig. 21a,b**). For the integrated ATAC and RNA in DBiT ARP-seq, we delineated major regions of the mouse brain at both 5 DPL and 21 DPL. These include all six cortical layers (II/III; J17, IV; J16, V; J14 and VI; J15); fiber tracts (cluster J2) such as CC, fimbria, internal capsule, and stria (medullaris and terminalis); the choroid plexus (cluster J1) of the lateral and third ventricles; the caudoputamen (cluster J12); the thalamus (clusters J10, J9, J8); and importantly the clusters elicited by neuroinflammation (J5) (**Fig. 4f**). The marker genes for RNA and ATAC were calculated (**Extended Data Fig. 21c,e**), such as *Sox10* and *Neurod6*, which are markers for oligodendrocytes and cortical layer neurons, respectively, and identified 19,804 significant linkages between regulatory elements and their corresponding target genes (**Extended Data Fig. 22**). For the integrated H3K27me3 and RNA in DBiT CTRP-seq (H3K27me3), we also produced detailed spatial UMAPs for mouse brains at 5 DPL and 21 DPL, with high correspondence between our two methodologies (DBiT tri-omic mapping and CODEX) (**Fig. 4g**). Finally the cell types in mouse brains at 5 DPL and 21 DPL were assigned identities using cell2location^53^ to map scRNA-seq data^26,54^ onto our spatial RNA data from both DBiT ARP-seq and DBiT CTPR-seq (**Extended Data Fig. 21g,h**). The results align with those observed in the CODEX image (where there was overlap) and known marker expression previously reported in LPC lesions^55–58^. For instance, the presence of activated lymphocytes (*Cd44+)* was confirmed by DBiT ARP-seq and DBiT CTPR-seq in all three modalities, localized to the lesion core, the injection site, and in the meninges at the brain surface and falx cerebri (**Fig. 4h,i**).

### A transient population of microglia arise during myelination in development and remyelination in the LPC models of MS

We then evaluated gene expression, chromatin accessibility (by calculating gene activity score, GAS), along with histone modification (by calculating the chromatin silencing score (CSS), for H3K27me3), for marker genes related to oligodendrocytes, neurons, and neuroinflammation (**Fig. 4h,i and Extended Data Fig. 21c-f**). At 5 DPL, *Olig2* expression is reduced in the lesion area at the levels of ATAC, RNA, and protein, accompanied by increased CSS for H3K27me3 (**Fig. 4h,i**). By 21 DPL, after remyelination, *Olig2* repopulated the lesion area in the corpus callosum, except for a remaining demyelinated region dorsal to the ventricles, and H3K27me3 exhibited low CSS (**Fig. 4h,i**). At 5 DPL, the expression of *Icam1* (CD54), typically expressed on endothelial and immune system cells^59,60^, was noted in both the lesion and ventricular areas across ATAC, RNA, and protein modalities, with low CSS for H3K27me3, aligning with previous observations. By 21 DPL, *Icam1* remained localized to the periventricular regions across ATAC, RNA, and ADT modalities, with persistently low CSS for H3K27me3 (**Fig. 4h,i**). *CD9,* a marker of immune cells and OPCs^61^, had heightened RNA expression, chromatin accessibility, and low CSS in both lesion and corpus callosum with differences in the levels (for RNA/ATAC, very high in the lesion, lower in the CC; for H3K27me3, very low in the lesion and reduced in the CC) (**Extended Data Fig. 24**).

Macrophage/microglial markers such as *Cx3cr1*, *Cd86*, and *Csf1r*^62–68^, predominated the lesion region across ATAC, RNA, and protein modalities, with consistently low CSS for H3K27me3 at 5 DPL (**Fig. 4h,i and Extended Data Fig. 24**). By 21 DPL, *Cx3cr1*, *Cd86*, and *Csf1r* were no longer at the area of the previous core lesion, but rather remained dominant in discrete periventricular demyelinated regions, where the CSS for H3K27me3 for these genes was low (**Fig. 4h,i and Extended Data Fig. 24**). Also, the CD11c*+*/*Itgax* pixels were present at both 5 and 21 DPL and populated regions of the primary lesion (5 DPL) and the unmyelinated CC dorsal to the lateral ventricle (21 DPL) (**Fig. 5b,c and Extended Data Fig. 25b**). Since CD11c+ microglia have been described to be also transient in development, appearing postnatally (P0-P7) in WM tracts like the CC, and diminishing in adolescent mice^69^, we investigated this population in our developing mouse brain CODEX and DBiT ARP-seq datasets. Indeed, we were able to track CD11c+ (*Itgax*) microglia at P0, residing mostly medial to the lateral ventricles until P5 when they localized to the CC dorsal to ventricles (**Fig. 5a and Extended Data Fig. 25**). Interestingly, we observed a large divergence between the chromatin accessibility and RNA expression of *Itgax* and *Itgam,* in which the accessibility pattern showed a broader distribution compared to the RNA expression profile, except at P21 (**Extended Data Fig. 25a**). While their chromatin accessibility at 5 DPL was broad, cells in the lesion had greater chromatin accessibility, and this pattern was maintained at 21 DPL in the unmyelinated region of the CC (**Fig. 5c**).

**Fig. 5:**
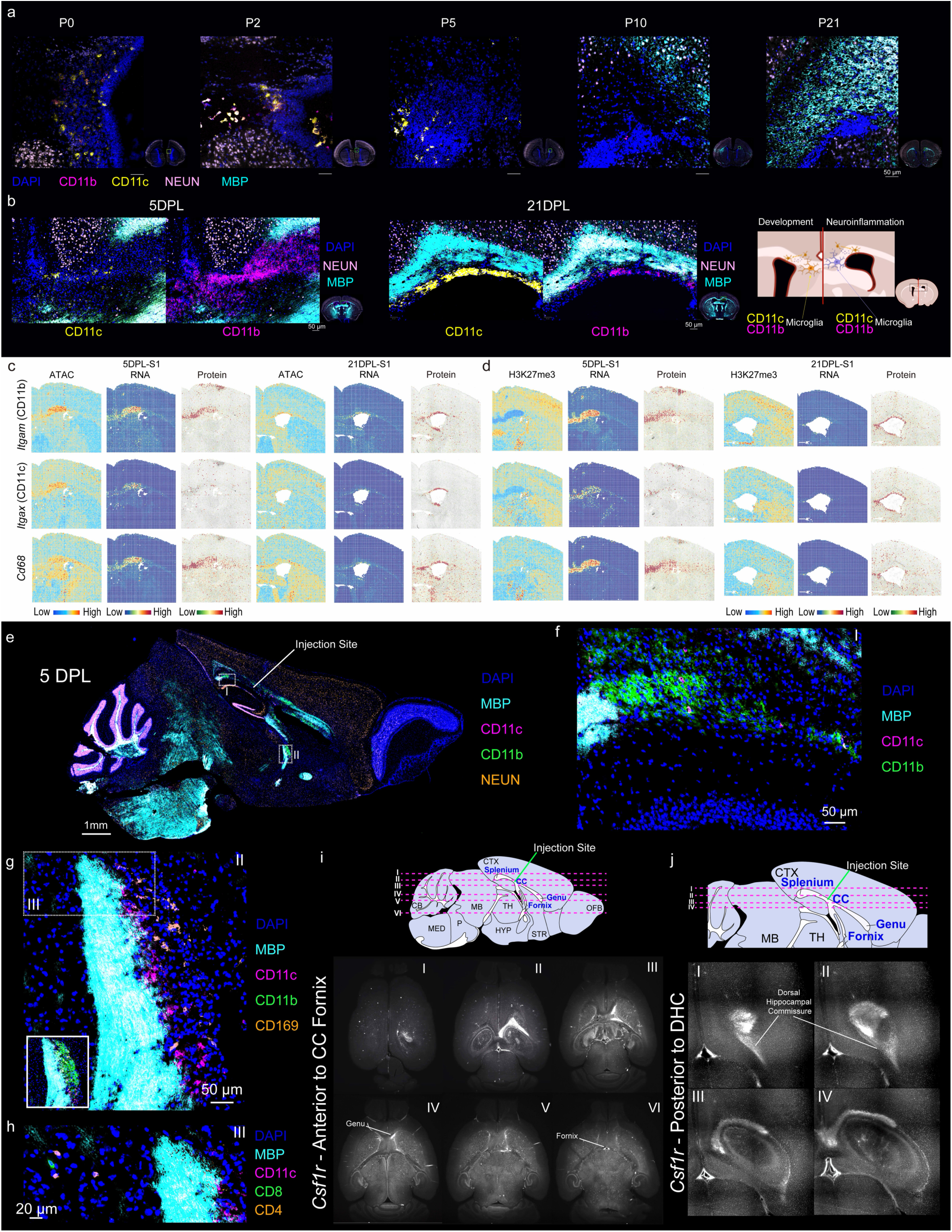
Microglia in both development and LPC mouse brains. **a,** CODEX images showing microglial subsets in the mouse brain from P0 to P21. **b,** CODEX images showing microglial cells in the LPC mouse brain at 5 DPL (left) and 21 DPL (middle). Right: schematic diagram depicting different types of microglia in development vs inflammation. **c-d,** Spatial mapping of gene expression, GAS, CSS, and ADT protein expression for *Itgam* (CD11b), *Itgax* (CD11c), and *Cd68* in both DBiT ARP-seq (**c**) and DBiT CTRP-seq (**d**). **e,** CODEX image of the LPC mouse brain on a sagittal tissue section at 5 DPL. **f,** Magnified image of the region of interest I indicated by white dashed rectangle I in **e**. **g,** Magnified image of the region of interest II indicated by the white dashed rectangle II in **e**. **h,** Magnified image of the region shown as the white dashed rectangle III in **g** with staining of CD4 and CD8. **i-j,** TRIC-DISCO results of *Csf1r* from anterior (**i**) and posterior (**j**) direction to injection.

We also observed a spatially broad distribution of protein expression and chromatin accessibility for *Cd68*, a marker for phagocytotic activity (**Fig. 5c**). *Cd68* is associated with more mature and functional microglia, which may represent sparse microglia activation in regions distant from the lesion.

### Distal microglia associated with WM tracks is observed far from the LPC lesion

We then performed CODEX on sagittal sections near the LPC injection site at 5 DPL. Surprisingly, we found CD11b+ cells located distally from the primary lesion but retained in WM tracks (**Fig. 5e-g**). The distal spread occurred in the directions anterior, posterior and ventral (**Fig. 5e I,II**) to the lesion (**Fig. 5e-g**). While mostly encompassing distal parts of the CC like the splenium and the bottom of the fornix, we also observed CD11b+ cells in the dorsal hippocampal commissure. Thus, our data suggests that a focal demyelination lesion event might trigger microglial activation in distant sites within the CNS.

To investigate whether distal microglia activation could occur in other regions not covered in our CODEX, we performed Tris-mediated retention of *in situ* hybridization signal during clearing DISCO (TRIC-DISCO), a probe-based strategy for cleared brain 3D imaging of specific RNA transcripts^70^. We performed a clearing, labeling, and imaging protocol for the microglial marker *Csf1r* on whole brain tissue extracted from mice 5 DPL (**Fig. 5i,j and Extended Data Videos 1-3**). Recapitulating and expanding on our CODEX imaging, we saw spreading to both hemispheres with a higher signal intensity on the side ipsilateral to the injection. In particular, we observed *Csf1r* spreading (anterior to the injection) through to the genu of the CC and ventrally to the fornix (**Fig. 5i**). In the posterior direction we saw *Csf1r* intensity to the splenium of the CC and along the dorsal hippocampal commissure (**Fig. 5j**). A closer look at the distal population showed CD11c+ microglia that co-stained for CD11b but not CD169 (**Fig. 5g**). The distal cluster along the fornix were found to be in close to CD4+ and CD8+ T-cells (**Fig. 5h**).

Thus, our CODEX and TRIC-DISCO analysis further indicates that induction of a focal demyelinating lesion in one area of the brain can lead to microglial activation distally.

## Discussion

Understanding the complexity of the multicellular programs that occur simultaneously in the CNS in development and disease has been a long-time fascination of neuroscientists. While the inherent diversity and heterogeneity were first appreciated by the initial drawings of neuronal and glial morphology^71^, the single-cell and spatial technologies of the last decade have enabled the exploration of the brain and spinal cord with unprecedented resolution. Spatial omics technologies provide unparalleled opportunities to further decipher the brain. Here, we provide a spatial multiomic compendium of the cellular and molecular processes underlying: (1) the development of the cerebral cortex and associated white matter (corpus callosum), benchmarked against human cortical development, and (2) the progression from focal demyelination (using LPC) to remyelination of the corpus callosum. Specially, the obtained spatiotemporal dynamics multiomics atlas revealed temporal persistence and spatial spreading of chromatin accessibility for cortical layer-defining transcription factors, as well as priming of chromatin accessibility of myelin genes across the corpus callosum during the myelination window, suggesting an orchestration between layer-specific projection neuron axonogenesis and myelination. Application of our technologies to diseases unveiled transient activation of microglia at the core of the LPC lesion and at distal locations.

Chromatin priming at the level of chromatin accessibility has been shown to occur in the context of the embryonic development of cortical layers^1^ and transition to disease-associated states^13^. Interestingly, while we could observe such priming at a temporal and spatial level at the corpus callosum in term of oligodendroglia lineage progression and myelination, we find instead epigenetic trailing of cortical layer-defining transcription factors. Given that some of these factors still exhibit transcription, albeit reduced with the progression of time, this might reflect a fading epigenetic memory of the developmental states.

Our analysis also indicates an association with the cortical layer neurons maturation and the differentiation of oligodendroglia and myelination at the cortex and, perhaps more unexpectedly, at the corpus callosum. We were able to track the position of projection neurons and their postnatal development successfully segregating the processes of axonogenesis and synaptogenesis in space and time. Callosal CPNs populating layer II/III were mainly defined by axonal guidance cues and axonogenesis that decreased over time. Subcerebral SCPNs and corticothalamic CTPNs were regionally defined to layers V-VI and had elements of a more mature state synaptogenesis, glutamate secretion, and dendrite morphogenesis. Mature oligodendrocytes and myelination expression had a gradient of expression from low (II/III) to high (VI) at P10-P21, with OPCs, COPs, and NFOLs present in each layer and reduced over time. Strikingly, we observed RNA hallmarks of different cortical neuronal populations in specific regions of the corpus callosum, which might be explained by RNA transport of mRNAs through the developing axons to the synapse compartments, where they might exert specific functions^70,72^. By following the projections and tracking the oligodendrocytes to the underlying white matter, we were able observe preferential myelination across the CC that again was associated with the state of neuronal maturation. We propose that MOLs first myelinate the corticofugal PNs that run along the lateral region of the CC and then move laterally to myelinate the CPN tracts once they are further along their developmental program (**Fig. 3i**).

We incorporated a focal lesion-based disease model to study demyelination, repair and immune mediated disease-associated states of neurons and glia in an analogous tissue location. Promoting OPC differentiation by the initiation of developmental programs or the removal of the breaks of differentiation associated with disease is one mechanism thought to be reparative in MS^52^. Since these processes revolve around not only the OPC/OL but also the dynamic processes of other cell types, the maintenance of the cellular context is critical. The simultaneous utilization of the developmental and disease DBiT ARP-seq datasets provides a way to benchmark each system through space and time. Similar to previous reports^69^, we observed a population of microglia that expressed a canonical dendritic cell marker, *Itgax*/CD11c. We were able to segregate this population in every layer of our data analysis. CD11c+ microglia have been described as integral to developmental myelin, furthermore they are thought to be reparative in disease models^69,73–78^. This is the first time this population has been identified in the LPC model. In aging and disease states (ischemia, stroke, injury AD, ALS, MS) the percentage of CD11c+ microglia is variable depending on location and disease state. In development and MS models (EAE and cuprizone) the CD11c+ microglia have been linked to myelination-promoting developmental and repair mechanisms, one such mechanism is the secretion of insulin growth factor (IGF1)^79–81^. Consistent with CD11c+ microglia being pro-repair we see more at 21 DPL localized to the only region of the CC that had not yet been remyelinated.

Through multiplex and whole tissue imaging strategies like CODEX and DISCO, respectively, we were able to track distal macrophages/microglia that were populated along WM tracts both anterior and posterior to the lesion. We found that a proportion of the distal CD11b+ macrophages/microglia were also CD11c+. Future studies will be aimed to elucidate how and why there is distal spread of macrophages/microglia. While we do not know the underlying mechanism of the spread, there are several potential drivers at play. It is possible that the spread of microglia is propagated through ventricular transport of inflammatory signaling cues, inflammatory cells nearby could set up the inflammation and finally perhaps the inflammatory signal is propagated along the axonal tract. While it is difficult to know if the T-cells are driving the distal changes, it is unlikely since we do not observe T-cells at high amounts in every area we have microglia expansion. It is more likely that the microglia influence the environment and can promote low levels of inflammation. The mechanism of ventricular spread makes sense considering some of the regions we observe the expansion enter. However, it is unlikely that the ventricles are the only mechanism since we see expansion in areas that are not periventricular. Finally, there is some evidence to suggest the axonal propagation of inflammation could occur especially in the visual pathway and optic nerve^82–86^. Since the spread appears to be propagated along WM tracts, a leading hypothesis is that neurons can transmit inflammatory cues to distal locations. Indeed, Karadottir and colleagues show that this is the case for the caudal cerebellar peduncle track, where focal demyelination induced by ethidium bromide also leads to distal microglia activation, which can in part modified by neuronal activity (De Faria Junior et al, companion paper).

By employing multiplexed immunofluorescence imaging (CODEX) and our newly developed spatial tri-omic technologies, DBiT-ARP (i.e. spatial ATAC–RNA–Protein-seq) and DBiT-CTRP (spatial CUT&Tag–RNA–Protein-seq), as well as TRIC-DISCO to the developing cortex and corpus callosum, and to a mouse model of de/remyelination, we show how multimodal spatial omics can unveil novel aspects of brain development and disease. These technologies can be further utilized to explore additional regions of CNS during development, as well as to investigate the mechanisms of neuroinflammatory mediated damage and repair.

## METHODS

### Animals

The present study followed some applicable aspects of the PREPARE^87^ planning guidelines checklist such as the formulation of the *in vivo* study, dialogue between scientists and the animal facility, and quality control of the *in vivo* components in the study. All animals were born, bred, and housed at Karolinska Institutet, Comparative Medicine Biomedicum animal facility (KMB). Mouse brain tissues (postnatal days P0, P2, P5, P10, and P21) were obtained from a mouse line generated by crossing *Sox10:Cre* animals (The Jackson Laboratory mouse strain 025807) on a C57BL/6j genetic background with *RCE:loxP* (eGFP) animals (The Jackson Laboratory mouse strain 32037-JAX) on a C57BL/6xCD1 mixed genetic background. Females with a hemizygous Cre allele were mated with males lacking the Cre allele, while the reporter allele was kept in hemizygosity in both females and males. In the resulting *Sox10:Cre-RCE:LoxP* (eGFP) progeny the entire OL lineage was labelled with eGFP.

None of the experimental animals in this study were subjected to previous procedures before enrollment in the study. All animals were free from mouse viral pathogens, ectoparasites endoparasites, and mouse bacteria pathogens. Mice were kept with the following light/dark cycle: dawn 6:00-7:00, daylight 7:00-18:00, dusk 18:00-19:00, night 19:00-6:00 and housing to a maximum number of 5 per cage in individually ventilated cages (IVC Sealsafe plus GM500, Tecniplast, Italy). General housing parameters such as relative humidity, temperature, and ventilation follow the European Convention for the Protection of Vertebrate Animals used for experimental and other scientific purposes treaty ETS 123, Strasbourg 18.03.1996/01.01.1991. Briefly, consistent relative air humidity of 55%±10, 22 °C and the air quality is controlled with the use of stand-alone air handling units supplemented with HEPA filtrated air. Monitoring of husbandry parameters is done using ScanClime® (Scanbur) units. Cages contained hardwood bedding (TAPVEI, Estonia), nesting material, shredded paper, gnawing sticks and card box shelter (Scanbur). The mice received a regular chow diet (CRM(P) SDS, United Kingdom and CRM(P), SAFE, Augy, France). Water was provided by using a water bottle, which was changed weekly. Cages were changed every other week. Cage changes were done in a laminar air-flow cabinet (NinoSafe MCCU mobile cage changing unit) provided with a HEPA H14 EN1822 filter (0.3 um particle size). Facility personnel wore dedicated scrubs, socks and shoes. Respiratory masks were used when working outside of the laminar air-flow cabinet. Both sexes were included in the study.

All experimental procedures on animals were performed following the European directive 2010/63/EU, local Swedish directive L150/SJVFS/2019:9, Saknr L150, Karolinska Institutet complementary guidelines for procurement and use of laboratory animals, Dnr. 1937/03-640 and Karolinska Institutet Comparative Medicine veterinary guidelines and plans (version 2020/12/18). The procedures described were approved by the regional committee for ethical experiments on laboratory animals in Sweden (Stockholms Norra Djurförsöksetiska Nämnd, Lic. nr. 1995/2019 and 7029/2020).

### Tissue and slide preparation

For all postnatal collection points except P21, we performed a decapitation and extracted the brain, immediately placing the tissue in Tissue Tek O.C.T compound over a bath of dry ice and 70% ethanol. For all mice older than P21 we anesthetized and then performed a transcardiac perfusion with gassed, ice-cold artificial cerebral spinal fluid. After, we decapitated and removed the brain following the same embedding procedure as the neonates.

All tissues were stored at –80℃ until further usage. Fresh 50×75 mm (Ted Pella Inc, no. NC1811932) 0.01% poly-L-lysine (Sigma, no. P1524-100) coated slides were prepared and stored at 4 ℃ and used within one week for mounting tissue. For brain tissue sectioning, the tissue was incubated for 30 minutes in the cryostat chamber at –20 ℃. Slides were simultaneously precooled (poly-L-lysine coated for DBiT and charged Superfrost Plus (Eprendia, no. 4951PLUS-001) for CODEX) and kept in the cryostat for the entire procedure. Sections were cut using an antiroll plate, to a thickness of 10 µm. Tissue collection was done by placing the cold slide in the correct position on top of the tissue then placing over a gloved backhand until the tissue was fully bound. The slide was immediately placed back in the cryostat chamber until sectioning was complete at which point all slides were stored at –80 ℃.

### Lysolecithin mouse model (1% LPC injection)

*Sox10:Cre-RCE:LoxP* (eGFP) mice aged 10-12 weeks were used in the LPC study. Briefly, under an aseptic technique, mice received an intraperitoneal injection of a non-steroidal anti-inflammatory analgesic (Carprofen; 5mg kg^−1^), then deeply anaesthetized with isoflurane, an ophthalmic ointment was applied in both eyes to prevent corneal desiccation, then the animal was placed over a warm pad for the entire procedure to prevent hypothermia. The site of the incision was shaved and sanitized with 2% chlorhexidine before cutting a 0.5 cm incision with a scalpel to expose the cranium. Using a stereotaxic frame the rough coordinates of −0.8 mm posterior, and 0.8 mm lateral were measured for a skull-drilling perforation. After completing the 1% lysolecithin solution was loaded into a pulled glass micropipette and the coordinates were recalculated then the micropipette was slowly inserted 1.3 mm deep (corpus callosum below cingulum bundle). The 2 µL injection was performed using an injection speed of 5 nL s^−1^. After the injection was complete the micropipette was kept in place for an additional 5 minutes to prevent efflux and then slowly pulled out of the brain. Monitoring of reflexes, respiratory rate and breathing pattern was performed during the entire surgical procedure. The skin was sutured with a non-absorbable suture (Vicryl, Ethicon), and the mouse was observed until regaining full consciousness and mobility. Post-operative care was administered daily for the next 72 hours. Mice were sacrificed at 5 and 21 days after surgery using the sampling procedure and sample collection method as the P21 mice (above).

### Tris-mediated retention of *in situ* hybridization signal during clearing DISCO

Three *Sox10:Cre-RCE:LoxP* (eGFP) mice, aged 8-10 weeks, were stereotactically injected with 1% LPC and sacrificed 5 days post-procedure, as described above. One exception in the sacrifice procedure was including a 4% PFA perfusion directly after the aCSF. Each brain was carefully extracted and then stored overnight in 4% PFA at 4℃ before processing through the full TRIC-DISCO protocol^70^. Briefly, a methanol gradient (60%, 80%, 100%,100%) was done and then stored overnight at −20℃. Successive steps to delipidate (100% dichloromethane), wash, and bleach (100% methanol + 5% hydrogen peroxide) were done with overnight incubations at 4℃. Whole brain in situ hybridization chain reaction protocol sequence was performed with 50% formamide wash buffer incubation until brain sinking followed by a three-day incubation (at 37℃) in 50% hybridization buffer with *Csf1r* DNA probe (1.6 pmol of the probe in 400 µL). Samples were then washed three times in 5X saline-sodium citrate (5X SSC) buffer with Tween20 (5X SSCT), incubated in amplification buffer overnight (room temperature), and then washed three times in 5XSSCT buffer. Tris-mediated signal retention involved three washing steps (for 1 hour at room temperature) in 500 mM of Tris-HCl, sample dehydration in 100% methanol (1-hour at room temperature) then dehydration and delipidation in 66% DCM and 33% methanol (3 hours) and finally two washes with 100% DCM. To promote tissue transparency an overnight dibenzl ether (DBE) incubation was performed with one DBE solution change. Whole brain light sheet microscopy using LaVision Ultramicroscope II (MiltenyBiotec) provided images that were used in the study.

### Human samples and slide preparation

De-identified human tissue samples from the second trimester were collected at Zuckerberg San Francisco General Hospital (ZSFGH), with the acquisition approved by the UCSF Human Gamete, Embryo and Stem Cell Research Committee (study number 10-05113). All procedures adhered to protocol guidelines, and informed consent was obtained prior to sample collection and use for this study. The de-identified third trimester and early postnatal tissues were acquired from the University of Maryland Brain and Tissue Bank via the NIH NeuroBioBank. For all samples, coronal cryosections of 10 μm were prepared on poly-L-lysine-coated glass slides (no. 63478-AS, Electron Microscopy Sciences) or 50×75 mm poly-L-lysine-coated glass slides, and stored at −80 °C for subsequent use.

### Multiplex immunofluorescence imaging (CODEX)

Immunofluorescence imaging was conducted using the Akoya Biosciences PhenoCycler^TM^ system, which is integrated with a Keyence BZ-X800 epifluorescence microscope. The protocol for staining and imaging fresh frozen tissue, as detailed in the manufacturer’s user manual, was strictly adhered to.

### DNA barcodes sequences, DNA oligos, and other key reagents

DNA oligos used for transposome assembly, PCR, and library construction are shown in Extended Data Table 1. All the DNA barcode sequences were provided in Extended Data Tables 2,3, and all other chemicals and reagents were listed in Extended Data Table 6.

### Spatial ATAC–RNA–Protein-seq (DBiT ARP-seq)

The fresh-frozen tissue slide was thawed for 10 minutes before fixing with 0.2% formaldehyde with 0.05 U/μL RNase Inhibitor (RI) and quenching with 1.25 M glycine for 5 minutes, respectively. After washing with 0.5× DPBS-RI three times, the tissue section was permeabilized with 0.5% Triton X-100 for 20 minutes and washed again with 0.5× DPBS-RI three times.

The ADT cocktails (diluted 20 times from the original stock) from BioLegend and self-conjugated antibodies (including intracellular and surface proteins for mouse brain) (Supplementary Table 4,5) were added and incubated for 30 minutes at 4 °C. The ADT cocktail was removed by washing three times with 0.5× DPBS-RI. The tissue was then washed for 15 minutes with a lysis buffer containing 0.001% Digitonin, 10 mM Tris-HCl (pH 7.4), 3 mM MgCl_2_, 0.01% NP-40, 1% BSA, 10 mM NaCl, 0.05 U/μL RNase Inhibitor, and 0.01% Tween-20. Subsequently, it was washed twice for 5 minutes each with a wash buffer composed of 0.1% Tween-20, 1% BSA, 10 mM Tris-HCl (pH 7.4), 3 mM MgCl_2_, and 10 mM NaCl. A transposition mix was prepared consisting of 5 μL homemade transposome, 50 μL 2× Tagmentation buffer, 33 μL 1× DPBS, 1 μL 1% Digitonin, 0.05 U/μL RNase Inhibitor, 10 μL nuclease-free H_2_O, and 1 μL 10% Tween-20. This mixture was then added to the sample and incubated at 37 °C for 30 minutes. Following this, 40 mM EDTA containing 0.05 U/μL RNase Inhibitor was added and incubated for 5 minutes at room temperature to halt the transposition process. Subsequently, the tissue section was washed three times with 0.5× PBS-RI for 5 minutes each. The RT mixture consisting of 3.1 μL 10 mM dNTPs each, 4.5 μL RNase-free water, 0.4 μL RNase Inhibitor, 12.5 μL 5× RT Buffer, 6.2 μL Maxima H Minus Reverse Transcriptase, 0.8 μL SuperaseIn RNase Inhibitor, 10 μL RT primer, and 25 μL 0.5× PBS-RI was added to the tissue. The tissue was then incubated for 30 minutes at room temperature and subsequently at 42°C for 90 minutes in a humidified chamber. After the reverse transcription reaction, the tissue was washed for 5 minutes with 1× NEBuffer 3.1 containing 1% RNase Inhibitor.

For the first *in situ* ligation of barcodes (barcodes A), the PDMS chip was positioned over the region of interest (ROI) on the tissue slide. To ensure proper alignment, a brightfield image was captured using a 10× objective on the Thermo Fisher Scientific EVOS FL Auto microscope (AMF7000) with EVOS FL Auto 2 Software Revision 2.0.2094.0. The PDMS device and tissue slide were then securely fastened using a homemade clamp. Initially, barcode A was combined with ligation linker 1: 10 μL of 100 μM ligation linker, 10 μL of 100 μM each barcode A, and 20 μL of 2× annealing buffer (2 mM EDTA, 100 mM NaCl, 20 mM Tris, pH 7.5-8.0) were thoroughly mixed. For each channel, a ligation master mixture was prepared, containing 2 μL of ligation mix (11 μL T4 DNA ligase, 72.4 μL RNase-free water, 27 μL T4 DNA ligase buffer, 5.4 μL 5% Triton X-100), 2 μL of 1× NEBuffer 3.1, and 1 μL of each annealed DNA barcode A (A1−A100, 25 μM). The ligation master mixture was loaded into 100 channels of the device using a vacuum. The device was then incubated at 37 °C for 30 minutes in a humidified chamber. After the incubation, the PDMS chip and clamp were removed following a 5-minute wash with 1× NEBuffer 3.1. The slide was subsequently washed with water and air-dried.

For the second *in situ* ligation of barcodes (barcodes B), the second PDMS chip was placed over the slide. A brightfield image was captured using a 10× objective for alignment purposes. The PDMS device and tissue slide were securely clamped together using a clamp. The annealing of barcodes B (B1−B100, 25 μM) and the preparation of the ligation master mix were performed identically to the procedure for barcodes A. The device was then incubated at 37°C for 30 minutes in a humidified chamber. After incubation, the PDMS chip and clamp were removed following a 5-minute wash with 1× DPBS containing SUPERase In RNase Inhibitor. The slide was then rinsed with water and air-dried. A bright-field image was taken afterwards to assist with further alignment.

After the barcoding process, the ROI on the tissue was treated with a reverse crosslinking mixture containing 1 mM EDTA, 0.4 mg/mL proteinase K, 50 mM Tris-HCl (pH 8.0), 1% SDS, and 200 mM NaCl. This mixture was incubated at 58°C for 2 hours in a humidified chamber. Following this, the lysate was transferred to a 1.5 mL tube and incubated at 65°C overnight.

To separate DNA and cDNA, the lysate was purified using the Zymo DNA Clean & Concentrator-5 and then eluted into 100 μL of RNase-free water. The Dynabeads MyOne Streptavidin C1 beads (40 μL) were washed three times using 1× B&W buffer containing 0.05% Tween-20. Afterward, the beads were resuspended in 100 μL of 2× B&W buffer with 2.5 μL of SUPERase In RNase Inhibitor. This bead suspension was then mixed with the lysate and allowed to bind at room temperature for 1 hour with agitation. Finally, a magnet was used to separate the beads from the supernatant in the lysate.

The supernatant was applied for ATAC-seq library construction and purified using Zymo DNA Clean & Concentrator-5, then eluted into 20 μL of RNase-free water. For the PCR, 2.5 μL of 25 μM indexed i7 primer, 25 μL of 2× NEBNext Master Mix, and 2.5 μL of 25 μM P5 PCR primer were added and thoroughly mixed. The initial PCR program was set as follows: 72°C for 5 minutes, 98°C for 30 seconds, followed by 5 cycles of 98°C for 10 seconds, 63°C for 30 seconds, and 72°C for 1 minute. To determine the need for additional cycles, 5 μL of the pre-amplified mixture was combined with a qPCR solution consisting of 0.5 μL of 25 μM new P5 PCR primer, 5 μL 2× NEBNext Master Mix, 3.76 μL nuclease-free water, 0.24 μL 25× SYBR Green, and 0.5 μL of 25 μM indexed i7 primer. The qPCR reaction was then performed using the following program: 98°C for 30 seconds, followed by 20 cycles of 98°C for 10 seconds, 63°C for 30 seconds, and 72°C for 1 minute. Based on the qPCR results, the remaining 45 μL of pre-amplified DNA was further amplified for additional cycles until it reached one-third of the saturation signal. The final PCR product was then purified using 45 μL of 1× Ampure XP beads and eluted in 20 μL of nuclease-free water.

The beads were employed to construct cDNA libraries. Initially, they were washed twice with 400 μL of 1× B&W buffer containing 0.05% Tween-20 and then once with 10 mM Tris (pH 8.0) with 0.1% Tween-20. The Streptavidin beads, with cDNA molecules bound, were resuspended in a TSO solution composed of 44 μL of 5× Maxima RT buffer, 22 μL of 10 mM dNTPs, 44 μL of 20% Ficoll PM-400 solution, 5.5 μL of RNase Inhibitor, 5.5 μL of 100 μM template switch primer, 88 μL of RNase-free water, and 11 μL of Maxima H Minus Reverse Transcriptase. The beads were incubated at room temperature for 30 minutes and then at 42°C for 90 minutes with gentle shaking. Following this, the beads were washed once with 10 mM Tris containing 0.1% Tween-20 and once with water, then resuspended in a PCR solution consisting of 91.9 μL of RNase-free water, 8.8 μL of 10 μM each of primers 1 and 2, 0.5 μL of 1 μM primer 2-citeseq, and 110 μL of 2× Kapa HiFi HotStart Master Mix. PCR thermocycling was conducted with an initial denaturation at 95°C for 3 minutes, followed by 5 cycles of 98°C for 20 seconds, 65°C for 45 seconds, and 72°C for 3 minutes. After the initial five cycles, the Dynabeads MyOne Streptavidin C1 beads were removed from the PCR solution and 25× SYBR Green was added at a 1× concentration. The samples were then subjected to further qPCR with the program set at 95°C for 3 minutes, cycled at 98°C for 20 seconds, 65°C for 20 seconds, and 72°C for 3 minutes for 15 cycles, with a final extension at 72°C for 5 minutes. The reaction was terminated once the qPCR signal began to plateau.

To differentiate and purify cDNAs derived from mRNA and ADT, we employed 0.6× SPRI beads according to the standard protocol. We mixed 120 µl of SPRI beads with 200 µl of the PCR product solution and allowed the mixture to incubate for 5 minutes. The supernatant, which contained the ADT-derived cDNAs, was then transferred to a 1.5-ml Eppendorf tube. The beads remaining in the original container were washed with 85% ethanol for 30 seconds and eluted in RNase-free water over 5 minutes. The mRNA-derived cDNAs were subsequently quantified using Qubit and BioAnalyzer. For further purification of the supernatant, we added 1.4× SPRI beads and incubated for 10 minutes. These beads were cleaned once with 80% ethanol and resuspended in 50 µl of water. An additional purification step was conducted by adding 100 µl of 2× SPRI beads and incubating for another 10 minutes. After two washes with 80% ethanol, the ADT-derived cDNAs were finally eluted with 50 µl of RNase-free water.

The sequencing libraries for the two types of cDNA products were constructed separately. For the mRNA library, we used the Nextera XT Library Prep Kit. We began by diluting 1 ng of purified cDNA in RNase-free water to a final volume of 5 μL. We then added 10 μL of Tagment DNA Buffer and 5 μL of Amplicon Tagment Mix, and incubated the mixture at 55°C for 5 minutes. Following this, 5 μL of NT Buffer was added and the mixture was incubated at room temperature for another 5 minutes. Next, the PCR master solution, consisting of 15 μL PCR master mix, 1 μL of 10 μM P5 primer, 1 μL of 10 μM indexed P7 primer, and 8 μL RNase-free water, was added.

The PCR was then conducted with the following program: an initial denaturation at 95°C for 30 seconds, followed by 12 cycles of 95°C for 10 seconds, 55°C for 30 seconds, 72°C for 30 seconds, and a final extension at 72°C for 5 minutes. The PCR product was purified using 0.7× Ampure XP beads to obtain the final library.

For the ADT cDNAs, the library was constructed using PCR. In a PCR tube, 45 µL of the ADT cDNA solution was mixed with 10 µM 2.5 µL of P5 index, 2.5 µL of a 10 µM customized i7 index, and 50 µL of 2× KAPA HiFi PCR Master Mix. The PCR conditions were set as follows: an initial denaturation at 95°C for 3 minutes, followed by six cycles of denaturation at 95°C for 20 seconds, annealing at 60°C for 30 seconds, and extension at 72°C for 20 seconds, with a final extension at 72°C for 5 minutes. The PCR product was then purified using 1.6× SPRI beads.

The size distribution and concentration of the library were analyzed using the Agilent Bioanalyzer High Sensitivity Chip prior to sequencing. Next Generation Sequencing (NGS) was then performed using the Illumina NovaSeq 6000 sequencer in paired-end 150 bp mode.

### Spatial CUT&Tag–RNA–Protein-seq (DBiT CTRP-seq)

The preparation of tissue slides, and antibody incubation followed the same protocol as used in the DBiT ARP-seq method. After these initial steps, the tissue section was washed with an NP40-Digitonin wash buffer (0.01% NP40, 0.01% Digitonin, 0.5 mM Spermidine, 150 mM NaCl, one tablet Protease Inhibitor Cocktail, 20 mM HEPES pH 7.5) for 5 minutes, followed by three washes with 0.5× DPBS-RI. The section was then washed with wash buffer (0.5 mM Spermidine, 150 mM NaCl, one tablet Protease Inhibitor Cocktail, 20 mM HEPES pH 7.5). The primary antibody was diluted 1:50 in antibody buffer (2 mM EDTA, 0.001% BSA, in NP40-Digitonin wash buffer) and applied to the tissue, which was then incubated overnight at 4°C. Following this, the secondary antibody (Guinea Pig anti-Rabbit IgG), diluted 1:50 in NP40-Digitonin wash buffer, was added and incubated for 30 minutes at room temperature. The tissue was washed with wash buffer for 5 minutes. Next, a 1:100 dilution of the pA-Tn5 adapter complex in 300-wash buffer (20 mM HEPES pH 7.5, 300 mM NaCl, one tablet Protease Inhibitor Cocktail, 0.5 mM Spermidine) was added and incubated at room temperature for 1 hour, followed by a 5-minute wash with 300-wash buffer. Tagmentation buffer (10 mM MgCl_2_ in 300-wash buffer) was then added and incubated at 37°C for 1 hour. Finally, to stop the tagmentation, 40 mM EDTA with 0.05 U/μL RNase Inhibitor was added and incubated at room temperature for 5 minutes. The tissue was washed three times with 0.5× DPBS-RI for 5 minutes for further processing.

For the reverse transcription, two ligations, and beads separation, the procedures followed those established in the DBiT ARP-seq protocol. For constructing the CUT&Tag library, the supernatant was purified using Zymo DNA Clean & Concentrator-5 and eluted into 20 µL of RNase-free water. The PCR mix, consisting of 2 µL of 10 µM each of P5 PCR primer and indexed i7 primer along with 25 µL of NEBNext Master Mix, was added and thoroughly mixed. The PCR program included an initial incubation at 58°C for 5 minutes, 72°C for 5 minutes, and 98°C for 30 seconds, followed by 12 cycles of 98°C for 10 seconds and 60°C for 10 seconds, with a final extension at 72°C for 1 minute. The PCR product was then purified using 1.3× Ampure XP beads according to the standard protocol and eluted in 20 µL of nuclease-free water. The construction of the cDNA libraries for mRNA and ADT was carried out following the earlier DBiT ARP-seq protocol.

Prior to sequencing, the size distribution and concentration of the library were assessed using the Agilent Bioanalyzer High Sensitivity Chip. Subsequently, next-generation sequencing (NGS) was performed using the Illumina NovaSeq 6000 sequencer in paired-end 150 bp mode.

### Data preprocessing

For ATAC and RNA, as well as CUT&Tag and RNA data, linker 1 and linker 2 were used to filter Read 2, and the sequences were converted to Cell Ranger ARC format (v.2.0.2, 10x Genomics). The genome sequences were in the newly formed Read 1, barcodes A and barcodes B were included in newly formed Read 2. Human reference (GRCh38) or mouse reference (GRCm38) was used to align the fastq files.

For protein data from ADTs, the raw FASTQ file of Read 2 was reformatted the same way as RNA. Employing default configurations of CITE-seq-Count 1.4.2, we quantified the ADT UMI counts associated with each antibody at various spatial locations.

### Data clustering and visualization

Initially, we determined the locations of pixels on the tissue sections from brightfield images using MATLAB 2020b. Subsequently, we generated spatial data files by employing the codes available here: (https://github.com/di-0579/spatial_tri-omics).

Seurat v.4.1^88^ was loaded in R v.4.1 to construct Seurat objects for each sample using RNA matrices. Attribute information was then added to each object, and the objects were merged using the merge function. Pixel normalization, logarithmic transformation of counts, and gene scaling were performed, followed by PCA, retaining the top 50 PCs. To reduce sample heterogeneity, the ‘RunHarmony’ function was used for integration, and ‘RunUMA’ and ‘FindClusters’ were applied to generate RNA clustering information. Finally, ‘metadata’ was extracted to enable the spatial visualization of ‘RNAcluster’ for each sample.

We utilized Library Signac v1.8^89^ in R v4.1 to read the ATAC data matrices. To integrate multiple datasets, functions from the GenomicRanges package were employed to establish a common peak set. Fragment objects were then created for each sample, peaks were quantified across datasets, and these quantified matrices were used to create a Seurat object for each dataset, with the Fragment object stored within each respective assay. The datasets were merged using the ‘merge’ function. To minimize sample heterogeneity, we used ‘FindIntegrationAnchors’ to identify integration anchors, employing LSI embeddings for integration. Subsequently, we performed ATAC clustering and extracted ‘metadata’ for spatial visualization of ‘ATACcluster’.

We employed the SpatialGlue^28^ package to integrate and cluster ATAC and RNA data, obtaining spatial domain information. The ArchR^90^ package was used to load both ATAC and RNA data, incorporating the spatial domain information from SpatialGlue. This enabled us to identify differential RNA expression and gene activity score (GAS) across various regions. We then extracted RNA expression matrices, ATAC GAS expression matrices, and CUT&Tag CSS expression matrices from ArchR objects for spatial gene visualization.

### Spatiotemporal data analysis

To better identify the spatiotemporal patterns of gene regulation using our developed spatial multi-omics technology, we proposed a computational framework that consists of the following three main steps:

#### (1) Spatiotemporal regression model fitting

For each gene in RNA with expression in more than 2% of cells across time points, we fit a negative binomial generalized additive model (NB-GAM) to account for the effects of library size, time, and spatial location on gene expression levels and smooth terms were used to represent the effects of time and spatial location on gene expression, as well as their interaction^91^. More specifically, for the RNA counts *Y*_*gc*_ for a given gene *g* and pixel *c* with gene-specific means *μgc* and dispersion parameters *ϕ*_*g*_, we have

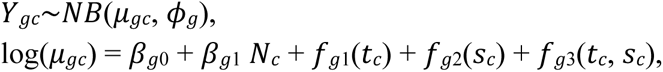

where *N* indicates the sequencing depth; *t*_*c*_ indicates the time point information (postnatal days 0, 2, 5, 10, 21) of pixel *c*; *s*_*c*_ indicates the spatial location of pixel *c*. In the analysis of the corpus callosum, the spatial location was defined by manually dividing the corpus callosum region into ten bins after manually delineating the region. In the analysis of cortical layers, the spatial location was defined using the SpatialGlue results. For a given *g*, *f*_*g*1_() is a smooth function capturing temporal variations, *f*_*g*2_() is a smooth function capturing spatial variations and *f*_*g*3_() is a tensor product smooth function capturing the interaction between time and spatial location. We used the cubie regression spline as the marginal basis of the smooth function. The model is fitted using *gam(),* with *ti()* as the smooth function and family as *nb()* from R package *mgcv*.

For ATAC data, we first derived the gene score matrix by aggregating the peak accessibility for peaks falling within a window ±50 kb of the TSS around each gene and expressed in more than 1% pixels across five time points, using the function *getDORCScores()* from R package *FigR*^92^. We smoothed the sparse gene score matrix per pixel per gene using its 4 nearest neighbors based on the first 50 principal components (PCs) from RNA data, using the function *smoothScoresNN().* We then fitted the GAM on the log-scaled smoothed gene score matrix with the gaussian family.

This model allows us to identify genes with significant associations with either time, spatial location, or their interaction. We determined the significance using Wald tests for the model coefficients and used the Benjamini-Hochberg (BH) procedure to adjust the p-values. Genes with an adjusted p-value of less than 0.01 in any of the spatial, time, or time and spatial interaction terms were selected for further analysis.

#### (2) Gene spatiotemporal clustering

We performed a two-step procedure to group patterns of genes that show significant changes across time and space in either RNA or ATAC. First, we derived pixel-level estimates *μ*_*gc*_ from our statistical model and aggregated them by their respective time points and spatial location regions, calculating RNA and ATAC separately. These aggregated profiles were then scaled. Next, we concatenated the RNA and ATAC aggregated profiles and then performed hierarchical clustering using one minus Pearson correlation coefficient as the distance metric and Ward’s minimum variance as a linkage method^93^, with the number of clusters set as 20 in the analysis of corpus callosum and set as 30 in the analysis of cortical layers. This procedure aims to jointly capture both ATAC and RNA patterns. The number of clusters was intentionally set higher to overestimate the number of clusters, allowing us to capture as many joint patterns of RNA and ATAC as possible.

As we observed that some patterns in RNA and ATAC are similar, we subsequently combined the clusters based on their similarity in RNA and ATAC profiles, respectively. First, we aggregated the RNA and ATAC profiles by their gene clustering labels from the previous step. Then, we performed hierarchical clustering with a smaller number of clusters: 6 for RNA and 4 for ATAC in the corpus callosum analysis, and 15 for RNA and 8 for ATAC in the cortical layers analysis. The number of clusters was determined by assessing whether the main patterns belonged to separate clusters and by evaluating the interpretability of the results. For the previous over-clustered results in the previous step, we assigned labels to represent their RNA and ATAC patterns. If clusters shared the same RNA and ATAC pattern labels, we combined these clusters. Each cluster is then labeled based on its RNA and ATAC pattern, for example, R1-A1 for a cluster where the RNA pattern belongs to R1 and the ATAC pattern belongs to A1. This step summarizes the RNA and ATAC patterns and improves the interpretability of each cluster. By characterizing genes with similar RNA and ATAC patterns separately, we can also jointly examine clusters with different RNA and ATAC patterns across space and time. For example, clusters labeled R1-A1 and R1-A3 share the same RNA pattern but have different ATAC patterns, while clusters labeled R1-A1 and R2-A1 share the same ATAC pattern but have different RNA patterns.

#### (3) Gene Ontology (GO) enrichment analysis of gene spatiotemporal clustering

We performed GO enrichment analysis to further characterize the identified clusters, focusing on the orthogonal ontology of biological processes. The analysis is performed using function *enrichGO()* from the R package *clusterProfiler*^94^ with *minGSSize* set as 20 and *maxGSSize* set as 200.

### CODEX data clustering

Whole cell segmentation was performed with Cellpose^95^ using the Cytoplasm model “cyto”. After obtaining the cell-level protein intensity matrix, we used Seurat v.4.1 to perform the subsequent analysis. For each sample, the data is first normalized using arcsinh transformation of the intensity matrix that scaled with a cofactor of 150^96^ and then scaled using *ScaleData()* and *RunPCA()* using all informative features. We removed the sample effect using the *FindIntegrationAnchors()* function, with Reciprocal PCA (rpca) set as the dimensional reduction method to identify anchors and the first 20 principal components (PCs) used as the dimensions. After integration, we performed clustering using the *FindClusters()* function on the neighbor graphs built based on the first 20 PCs, with the resolution set to 0.4. Each cluster was then manually annotated based on their highly expressed markers.

### Microglia subtype identification using ADT protein data

To determine the local region with enrichment of protein abundance, we calculated the G* local spatial statistics^97^ of the protein abundance of CD11b and CD11c based on the spatial weights derived from 12 spatial nearest neighbors. The higher value of G* indicates the higher possibility of a local cluster of high abundance of a certain protein. In the analysis of mouse development, we considered pixels with G* statistics greater than 4 as regions enriched with CD11b or CD11c. For the analysis of the LPC mouse model, we considered G* values greater than 4 for CD11b and greater than 3 for CD11c as indicative of enrichment.

### Cell2location

We performed cell2location (v0.1.3)^53^ to deconvolute the cell types of our spatial transcriptomics data using public references. We combined all samples in mouse development or in mouse LPC model together. We set the model *cell2location.models.Cell2location()* with the expected average cell abundance as 5 and trained with full data with maximum epochs set as 30000. The estimated cell abundance is based on the posterior mean using *export_posterior()*.

### Reporting summary

Further information on research design is available in the Nature Research Reporting Summary linked to this paper.

## Data availability

Raw and processed data reported in this paper will be deposited in the Gene Expression Omnibus (GEO) for mouse data, and in the European Genome-phenome Archive (EGA) for human data. These datasets are available as a web resource and can be browsed within the tissue spatial coordinates in our own data portal: https://spatial-omics.yale.edu/

The resulting fastq files were aligned to the human reference genome (GRCh38) (https://hgdownload.soe.ucsc.edu/goldenPath/hg38/chromosomes/) or mouse reference genome (GRCm38) (https://hgdownload.soe.ucsc.edu/goldenPath/mm10/chromosomes/). Published data for integration and quality comparison are available online: Atlas of gene regulatory elements in adult mouse cerebrum (http://catlas.org/mousebrain/#!/downloads); atlas of the adolescent mouse brain (http://mousebrain.org/adolescent/downloads.html); developing mouse brain data (https://www.ncbi.nlm.nih.gov/geo/query/acc.cgi?acc=GSE110823); mouse brain (https://assets.nemoarchive.org/dat-qwqfftg); CNS inflammation (https://www.ncbi.nlm.nih.gov/geo/query/acc.cgi?acc=GSE130119); and atlas of adult mouse brain (https://assets.nemoarchive.org/dat-qg7n1b0)

## Code availability

Code for sequencing data analysis is available at Github: https://github.com/di-0579/spatial_tri-omics

## Acknowledgments

We would like to thank Dr. Jens Hjerling-Leffler for the discussion and the Yale Center for Research Computing for guidance and use of the research computing infrastructure. We would also like to thank Dr. Eneritz Agirre for her expert advice and guidance in the computation analysis. The molds for microfluidic devices were fabricated at the Yale University School of Engineering and Applied Science (SEAS) Nanofabrication Center. Next-generation sequencing was conducted at the Yale Center for Genome Analysis (YCGA) as well as the Yale Stem Cell Center Genomics Core Facility which was supported by the Connecticut Regenerative Medicine Research Fund and the Li Ka Shing Foundation. Service provided by the Genomics Core of Yale Cooperative Center of Excellence in Hematology (U54DK106857) was used. We would like to thank the staff at Comparative Medicine-Biomedicum for assistance with animal husbandry and care. Work in G.C.-B.’s research group was supported by the Swedish Research Council (grants 2019-01360 and 2023-00324), the Swedish Cancer Society (Cancerfonden; 23 2945 Pj 01 H), Knut and Alice Wallenberg Foundation (grant 2019-0107,2019-0089 and 2023-0280), Swedish Brain Foundation (grant FO2023-0032), The Swedish Society for Medical Research (SSMF, grant JUB2019), the Göran Gustafsson Foundation for Research in Natural Sciences and Medicine, and Karolinska Institutet. Leslie Kirby was supported by European Committee for Treatment and Research in Multiple Sclerosis (ECTRIMS) and National MS Society (NMSS, USA, TA-2305-41342). Yonglong Dang was supported by Cancerfonden. This research was supported by Packard Fellowship for Science and Engineering (to R.F.), Yale Stem Cell Center Chen Innovation Award (to R.F.), and the U.S. National Institutes of Health (NIH) (RF1MH128876, U54AG076043, U54AG079759, UG3CA257393, UH3CA257393, R01CA245313, U54CA274509 to R.F.).

## Author Contributions

Conceptualization: R.F., D.Z., G.C.-B. and L.R.R-K.; Methodology: D.Z. and L.R.R.-K.; Experimental Investigation: D.Z. and L.R.R.-K.; Model design: D.Z., L.R.R.-K. and Y.Lin; Model development: Y.Lin; Data Analysis: D.Z., L.R.R.-K, Y.Lin., X.L., L.J.W., H.Z., G.C.-B. and R.F.; Data Interpretation: L.R.R.-K,. D.Z., G.C.-B and R.F.; Mouse husbandry: L.R.R.-K; Resources: L.R.R.-K., M.S., L.W., S.K., T.J.-B, Y.D., M.Z., P.K., S.W., X.L.C, F.G., D.W., H.X., Y.Liu. J.C., N.S., P.U. and A.R.K; Data browser: Y.Lin. and L.J.W.; Original Draft: D.Z., L.R.R.-K, Y.Lin., G.C.-B. and R.F. All authors reviewed, edited, and approved the manuscript.

