## Supplementary Information for "Spatial dynamics of mammalian brain development and neuroinflammation by multimodal tri-omics mapping"

[1]Department of Biomedical Engineering, Yale University, New Haven, CT 06520, USA
[2]Yale Stem Cell Center and Yale Cancer Center, Yale School of Medicine, New Haven, CT 06520, USA
[3]Laboratory of Molecular Neurobiology, Department of Medical Biochemistry and Biophysics, Karolinska Institutet, Stockholm, Sweden
[4]Department of Pathology, Yale University School of Medicine, New Haven, CT, USA
[5]Department of Biostatistics, Yale School of Public Health, New Haven, CT 06510, USA
[6]The Eli and Edythe Broad Center of Regeneration Medicine and Stem Cell Research, University of California San Francisco; San Francisco, CA 94143, USA
[7]Department of Neurology, University of California San Francisco; San Francisco, CA 94143, USA
[8]Yale Stem Cell Center and Department of Cell Biology, Yale University School of Medicine, New Haven, CT 06520, USA
[9]Department of Neurology, Yale School of Medicine, New Haven, CT, USA.
[10]Institute of Molecular and Cell Biology (IMCB), Agency for Science, Technology and Research (A*STAR), Singapore, Singapore
[11]Binformatics Institute (BII), Agency for Science, Technology and Research (A*STAR), Singapore, Singapore
[12]Department of Neuroscience, Yale School of Medicine, Yale University, New Haven, CT, USA
[13]Program of Computational Biology and Bioinformatics, Yale University, New Haven, CT, USA
[14]Human and Translational Immunology Program, Yale School of Medicine, New Haven, CT 06520, USA
[15]These authors contributed equally.

### Table of Contents

**3** Extended Data Figs. 1-25
**32** Extended Data Tables 1-6
**50** Extended Data Tables 7-8 (legends included, tables in separate Excel files)
**50** Extended Data Videos 1-3 (legends included, videos in separate .mp4 files)

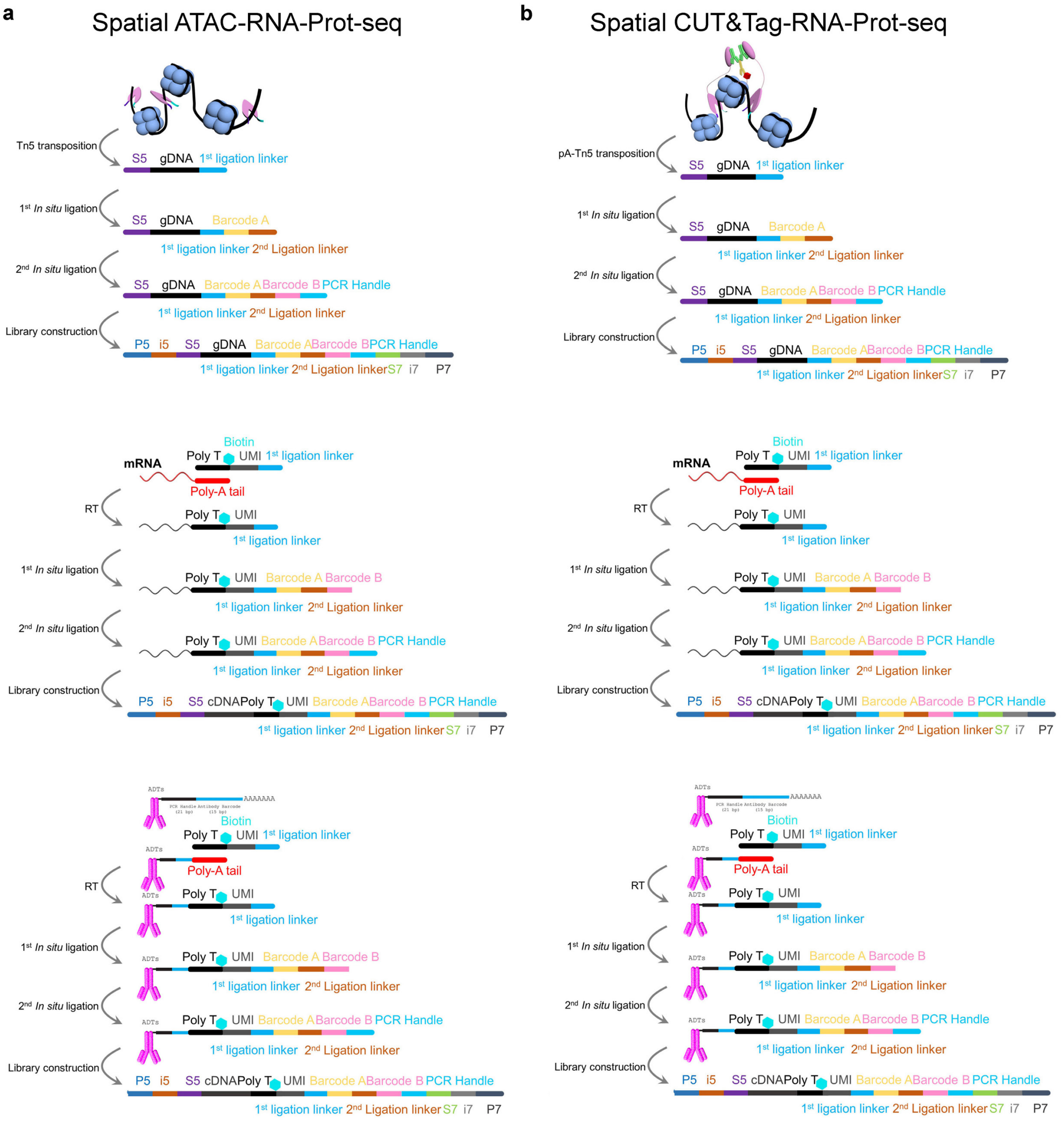


**Extended Data Fig. 1 Workflow of spatial ATAC–RNA–Prot-seq (DBiT ARP-seq) and spatial CUT&Tag–RNA–Prot-seq (DBiT CTRP-seq). a**, Chemistry workflow of ATAC (top), RNA (middle), and protein (bottom) in DBiT ARP-seq. **b**, Chemistry workflow of CUT&Tag (top), RNA (middle), and protein (bottom) in DBiT CTRP-seq.

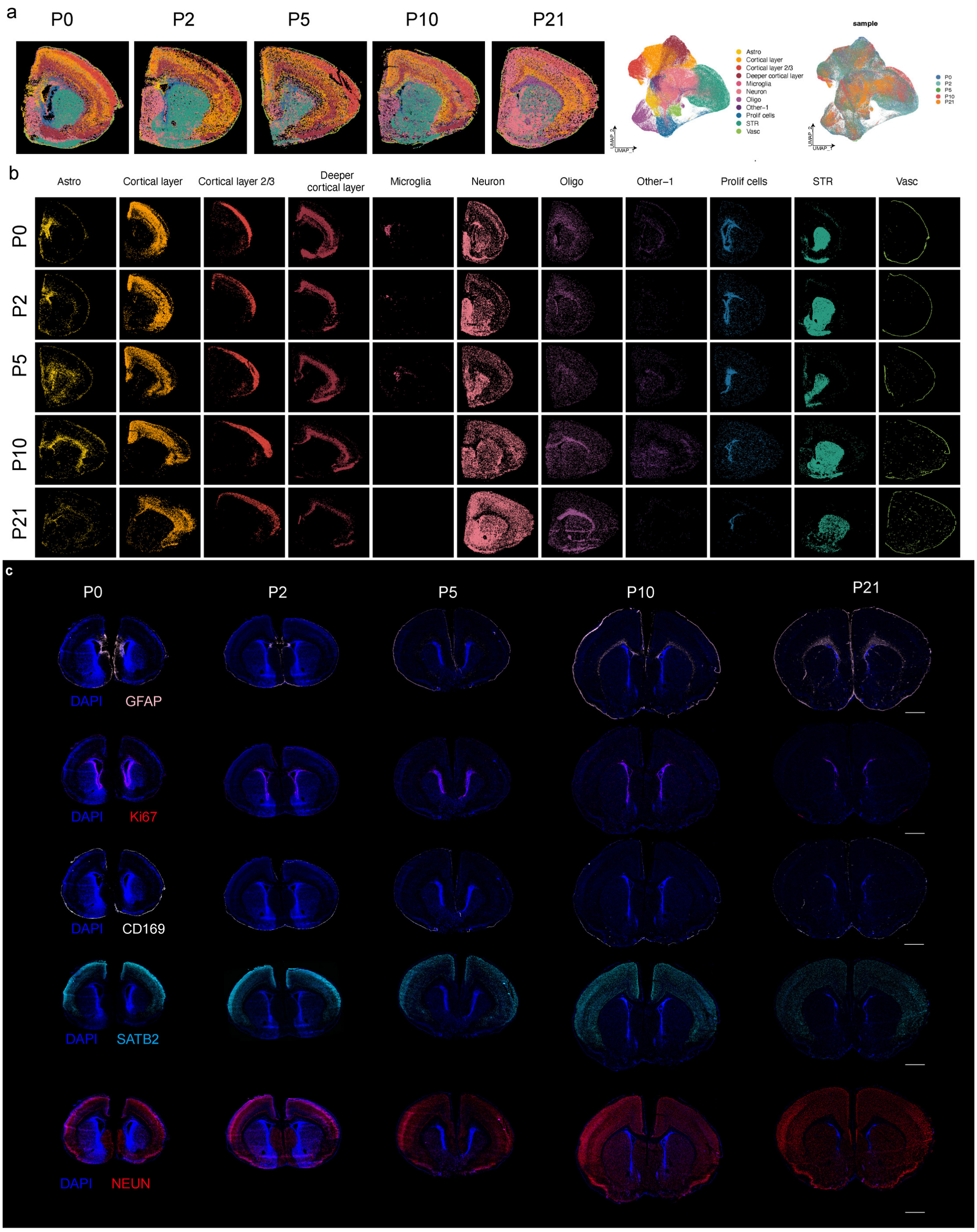


**Extended Data Fig. 2 Further data analysis of CODEX images for postnatal mouse brains.** **a,** Seurat clustering of the CODEX images in **Fig. 1b**. **b,** Spatial map of the cell types from **a**. **c,** CODEX images of GFAP, Ki67, CD169, and SATB2 for postnatal mouse brains. Scale bar, 1 mm.

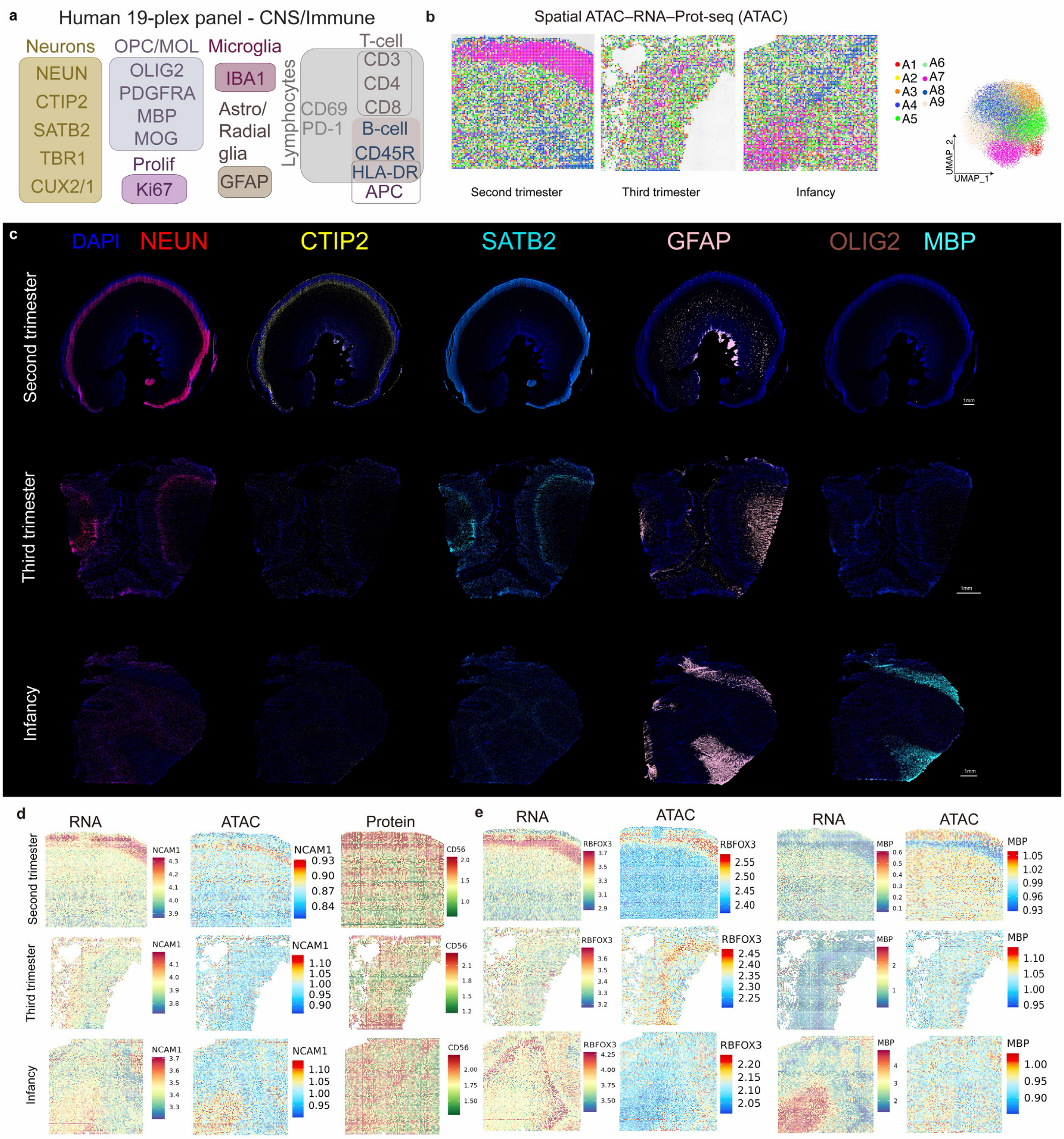


**Extended Data Fig. 3 Further data analysis of human brains in spatial ATAC–RNA–Prot-seq (DBiT ARP-seq).** **a**, CODEX protein panel for human. **b,** ATAC UMAP and spatial distribution of ATAC data of the human brain V1 regions at different stages. **c,** CODEX images of NEUN, CTIP2, SATB2, GFAP, OLIG2, and MBP for human brain V1 region at second trimester, third trimester, and infancy. Scale bar, 1 mm. **d,** Spatial mapping of gene expression, gene activity score (GAS), and ADT protein expression for *NCAM1* (CD56). **e,** Spatial mapping of gene expression and GAS for *RBFOX3* and *MBP*.

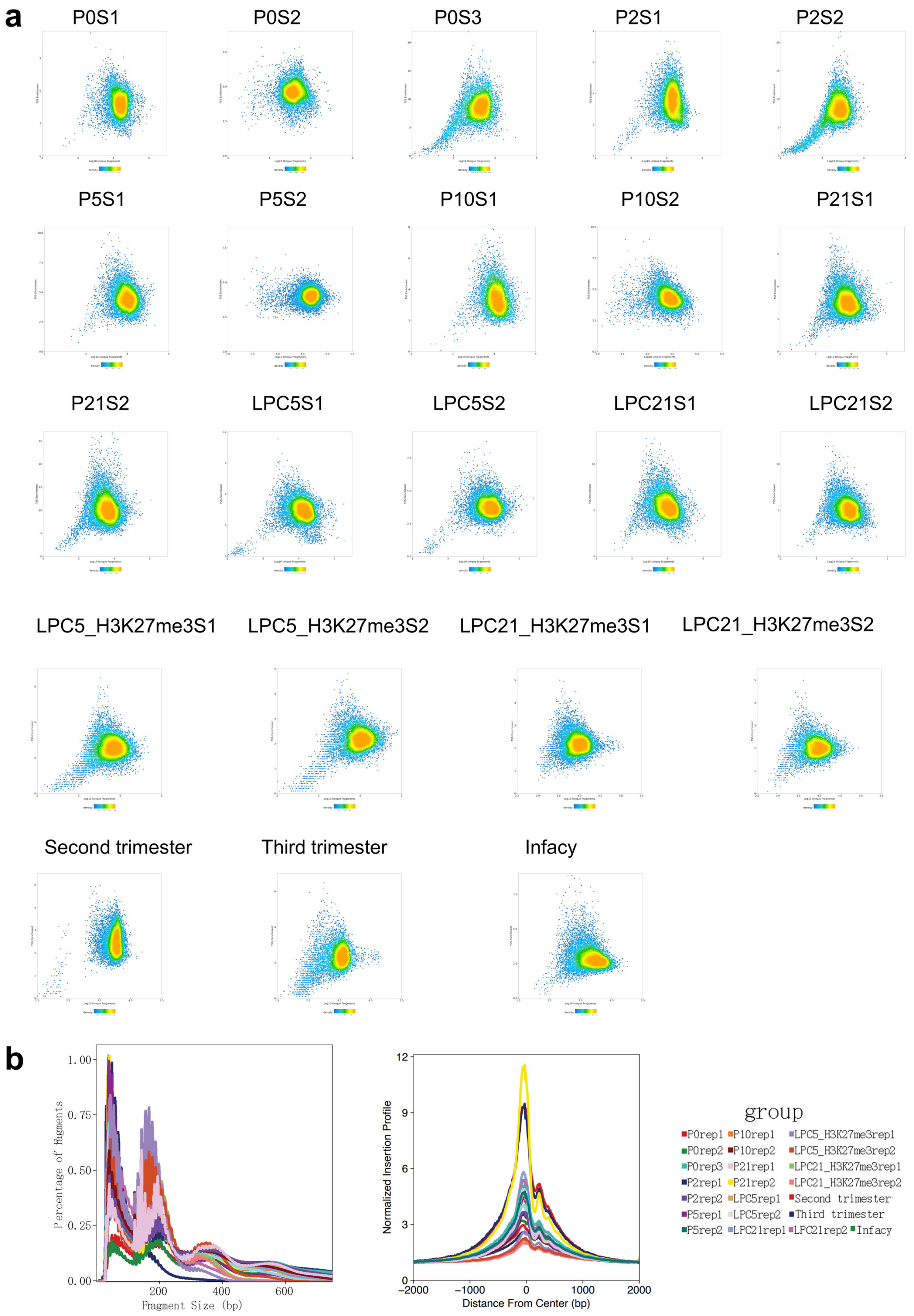


**Extended Data Fig. 4 Quality control metrics for DBiT ARP-seq and DBiT CTRP-seq datasets. a,** Scatterplots showing the TSS enrichment score vs unique nuclear fragments per pixel for all the samples finished. **b,** The insert size distribution of ATAC and CUT&Tag fragments (left) and the enrichment of ATAC or CUT&Tag reads around TSSs (right) in DBiT ARP-seq and DBiT CTRP-seq.

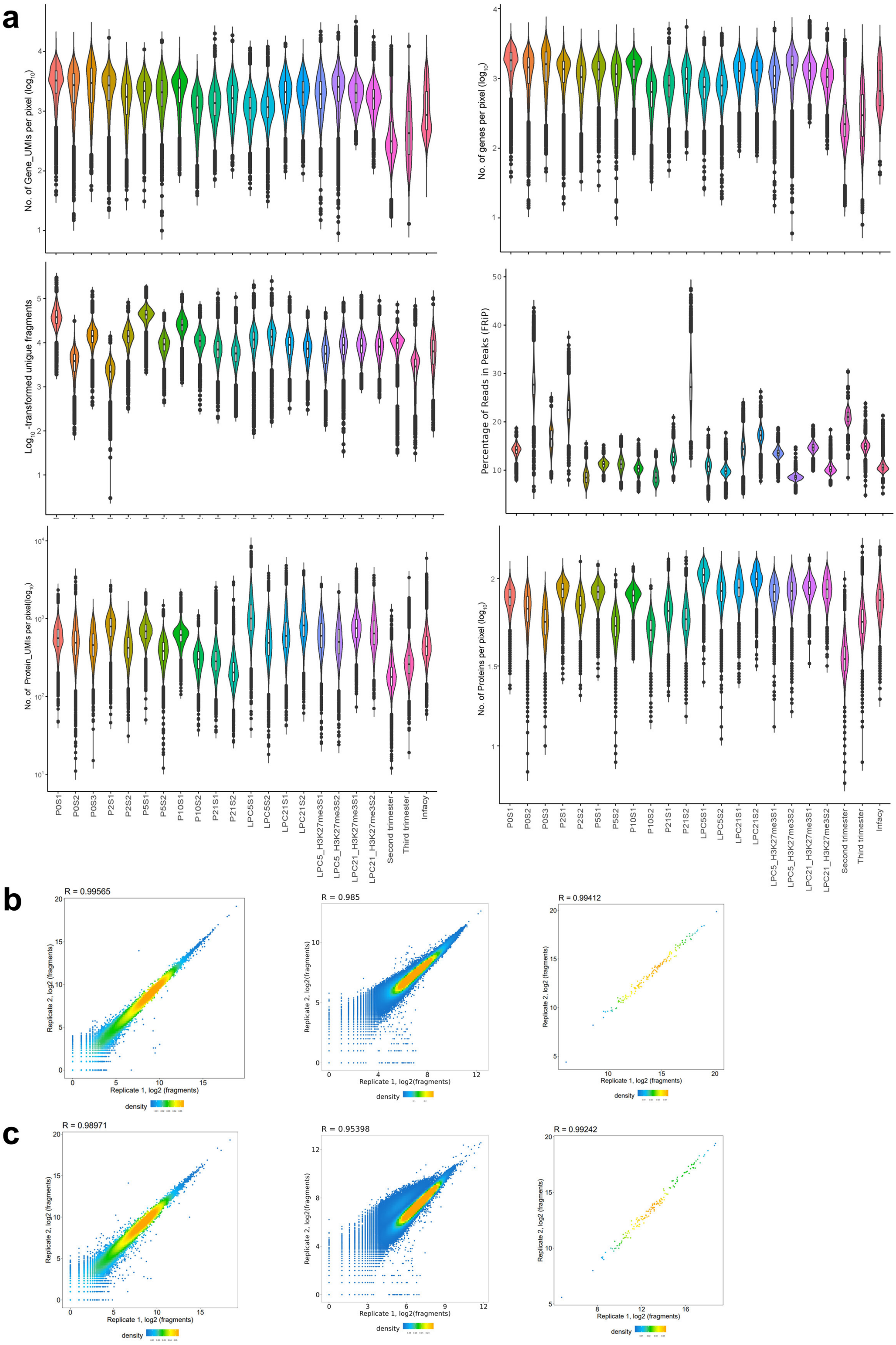


**Extended Data Fig. 5 Quality control metrics for DBiT ARP-seq and DBiT CTRP-seq datasets. a,** Gene and UMI count distribution (upper), comparison of number of unique fragments and fraction of reads in peaks (FRiP) (middle), ADT protein and UMI count (bottom) of processed samples for DBiT ARP-seq and DBiT CTRP-seq (H3K27me3). The box plots show the median (centre line), the first and third quartiles (box limits), and 1.5x the interquartile range (whiskers). **b,** The reproducibility of DBiT ARP-seq between biological replicates on ATAC data (left), RNA data (middle), and ADT protein data (right) for 5DPL mouse brains. **c,** The reproducibility of DBiT CTRP-seq (H3K27me3) between biological replicates on CUT&Tag data (left), RNA data (middle), and ADT protein data (right) for 5DPL mouse brains.

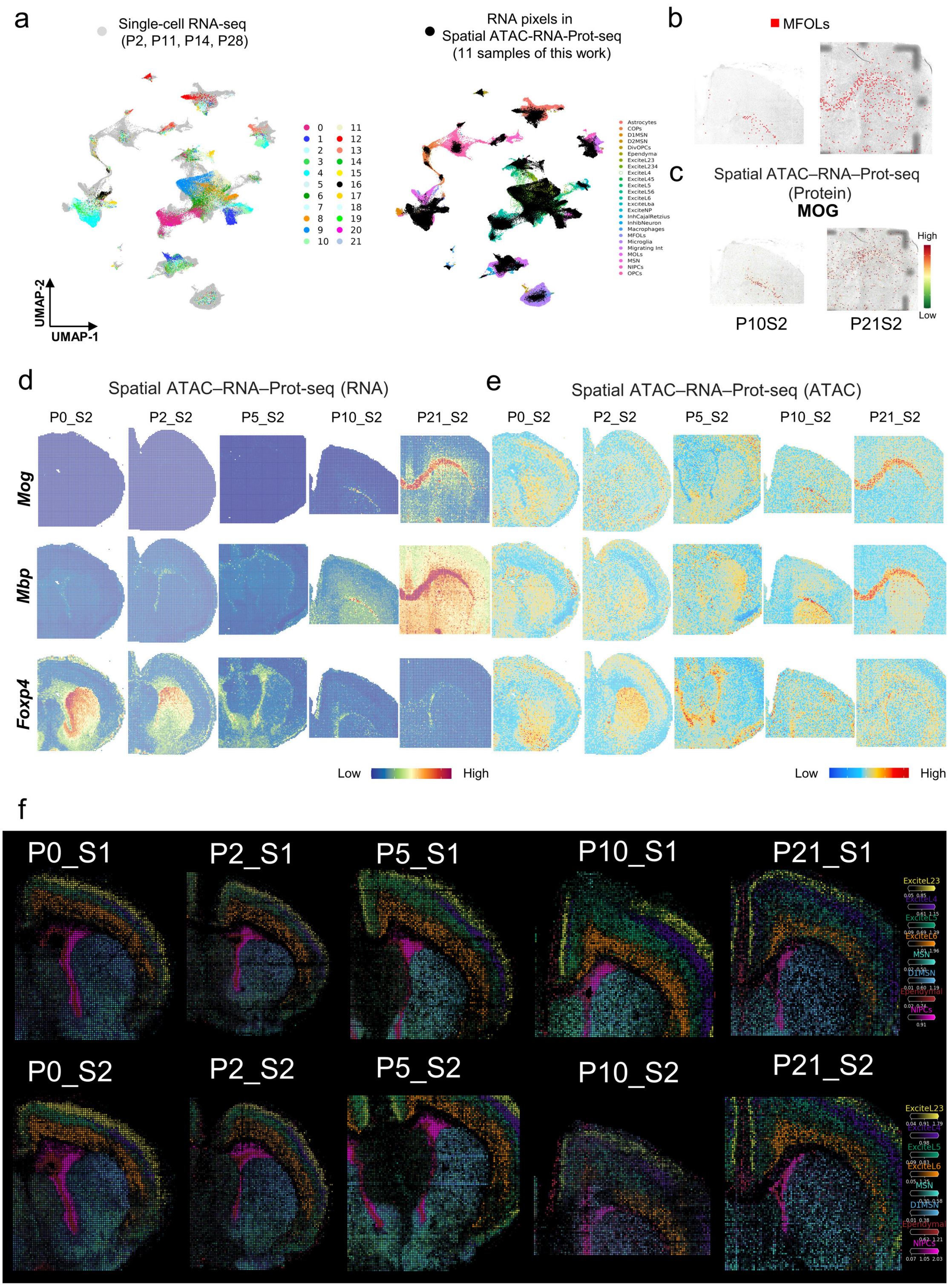


**Extended Data Fig. 6 Further analysis of spatial ATAC–RNA–Prot-seq (DBiT ARP-seq) for postnatal mouse brains. a,** Integration of scRNA-seq data from P2, P11, P14, P28 mouse brains with our spatial RNA data in DBiT ARP-seq. **b,** Spatial mapping of MFOLs identified by label transfer from scRNA-seq to spatial RNA of P10 and P21 mouse brains. **c,** MOG expression in P10 and P21 mouse brain from the ADT protein data in DBiT ARP-seq. **d-e,** spatial mapping of gene expression (**d**) and GAS (**e**) for selected marker genes from replicates in DBiT ARP-seq. **f,** Cell types predicted by cell2location from all processed postnatal mouse brain samples.

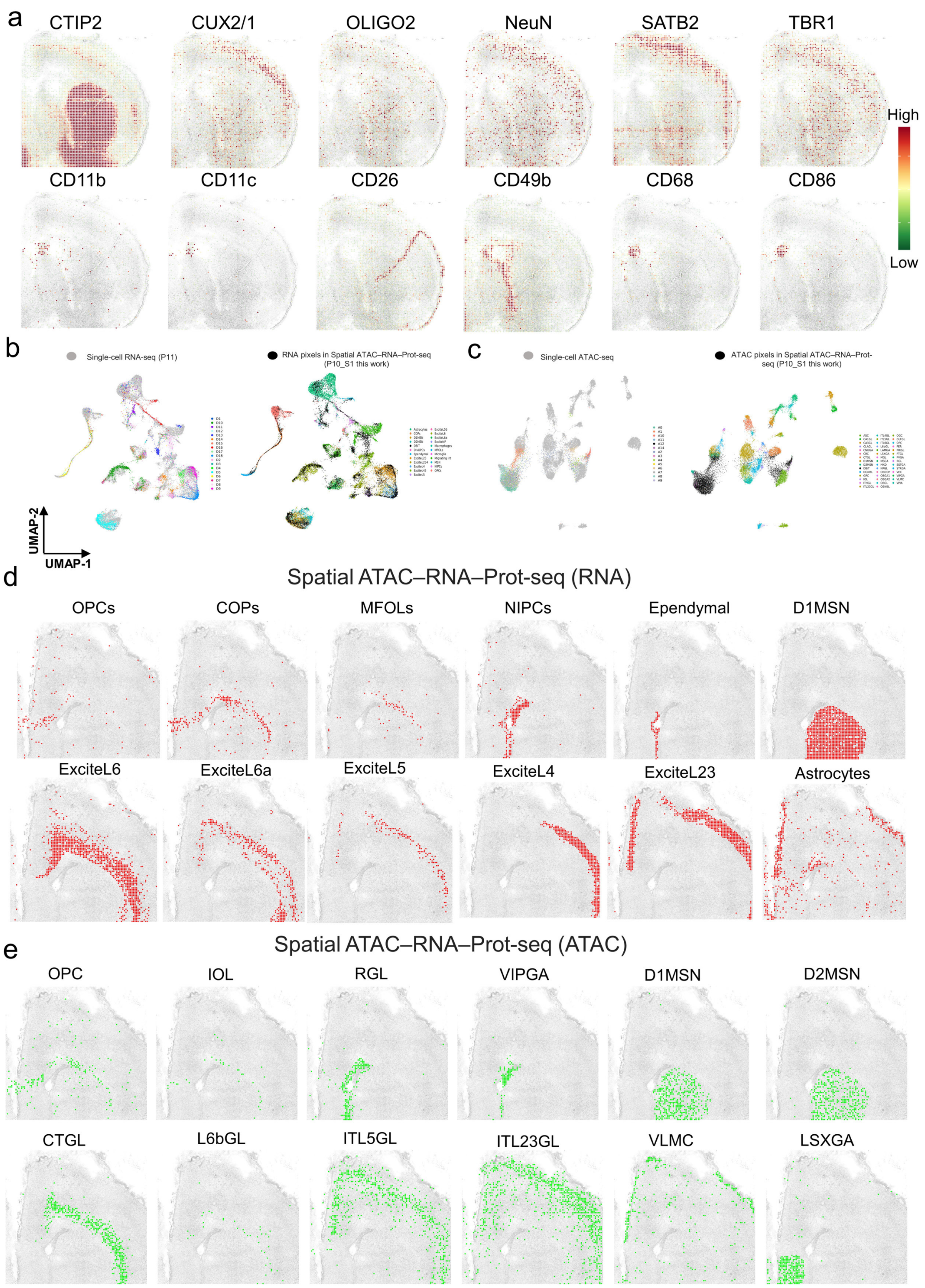


**Extended Data Fig. 7 Further analysis of spatial ATAC–RNA–Prot-seq (DBiT ARP-seq) for postnatal mouse brains. a,** Expression of several ADT proteins in DBiT ARP-seq from P0. **b,** Integration of P10 spatial RNA data and scRNA-seq data from mouse brain. **c**, Integration of P10 spatial ATAC data and scATAC-seq data from mouse brain. **d**, Spatial mapping of cell types identified by label transfer from scRNA-seq to P10 spatial RNA data. **e**, Spatial mapping of cell types identified by label transfer from scATAC-seq to P10 spatial ATAC data.

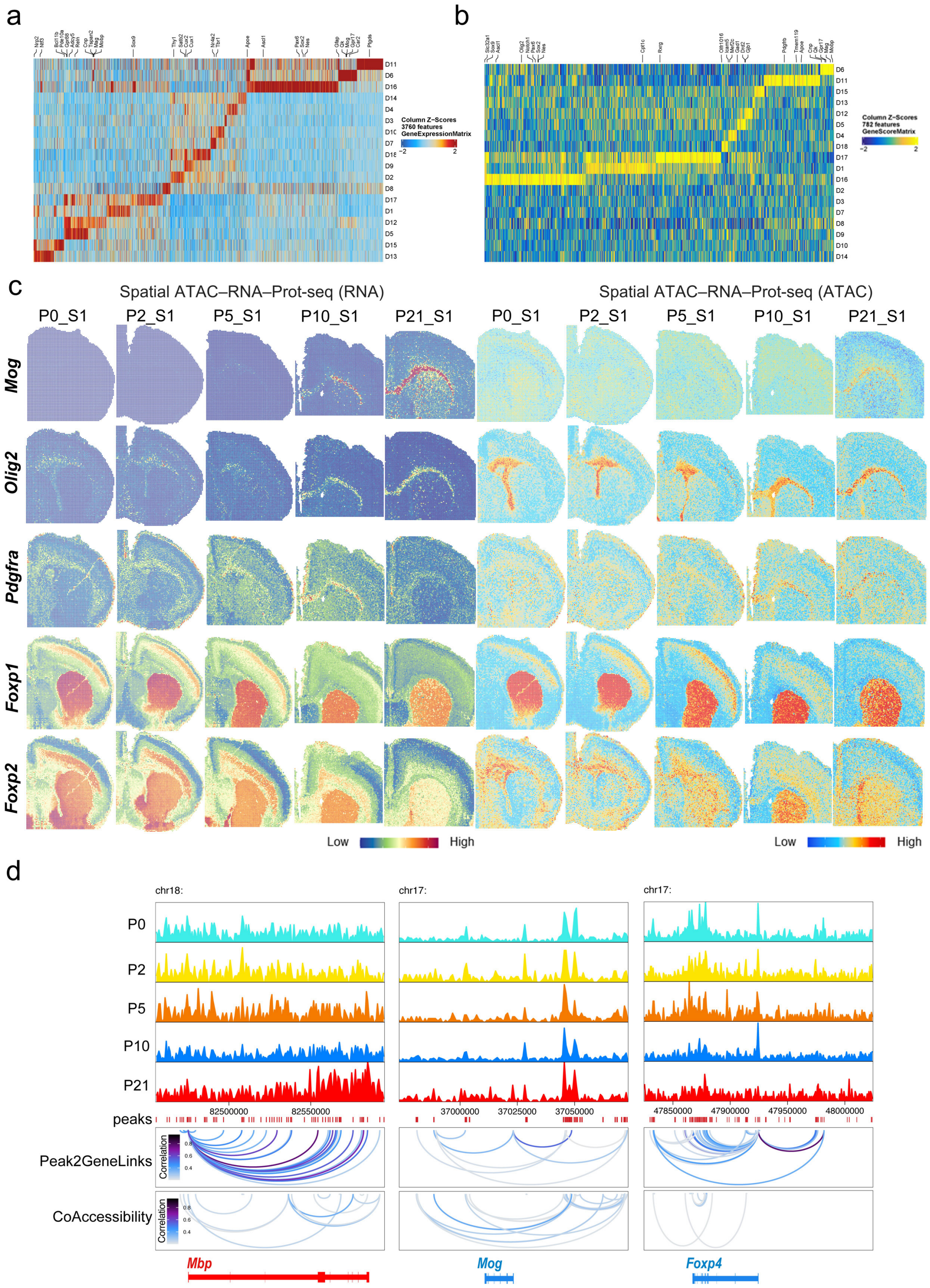


**Extended Data Fig. 8 Further analysis of spatial ATAC–RNA–Prot-seq (DBiT ARP-seq) for postnatal mouse brains. a,** Marker gene expression in each spatial domain in **Fig. 1i**. **b,** Marker GAS in each spatial domain in **Fig. 1i**. **c,** Spatial mapping of gene expression and GAS for selected marker genes in different clusters for RNA and ATAC in DBiT ARP-seq. **d,** Genome track visualization of marker genes with peak-to-gene links for distal regulatory elements and peak co-accessibility.

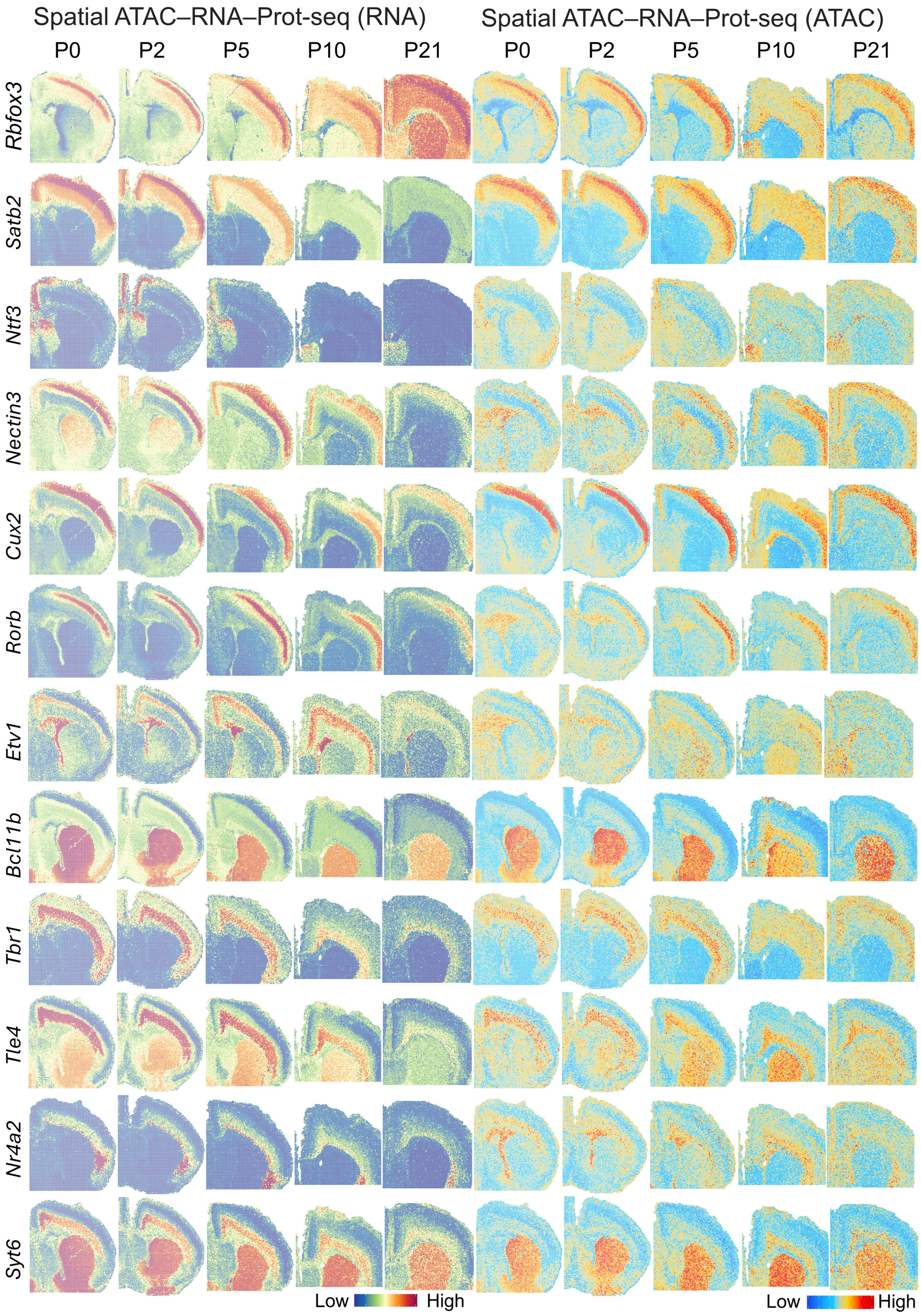


**Extended Data Fig. 9 Gene expression of cortical-layer specific markers for postanal mouse brains.** Spatial mapping of gene expression and GAS for neuronal marker genes from RNA and ATAC in DBiT ARP-seq.

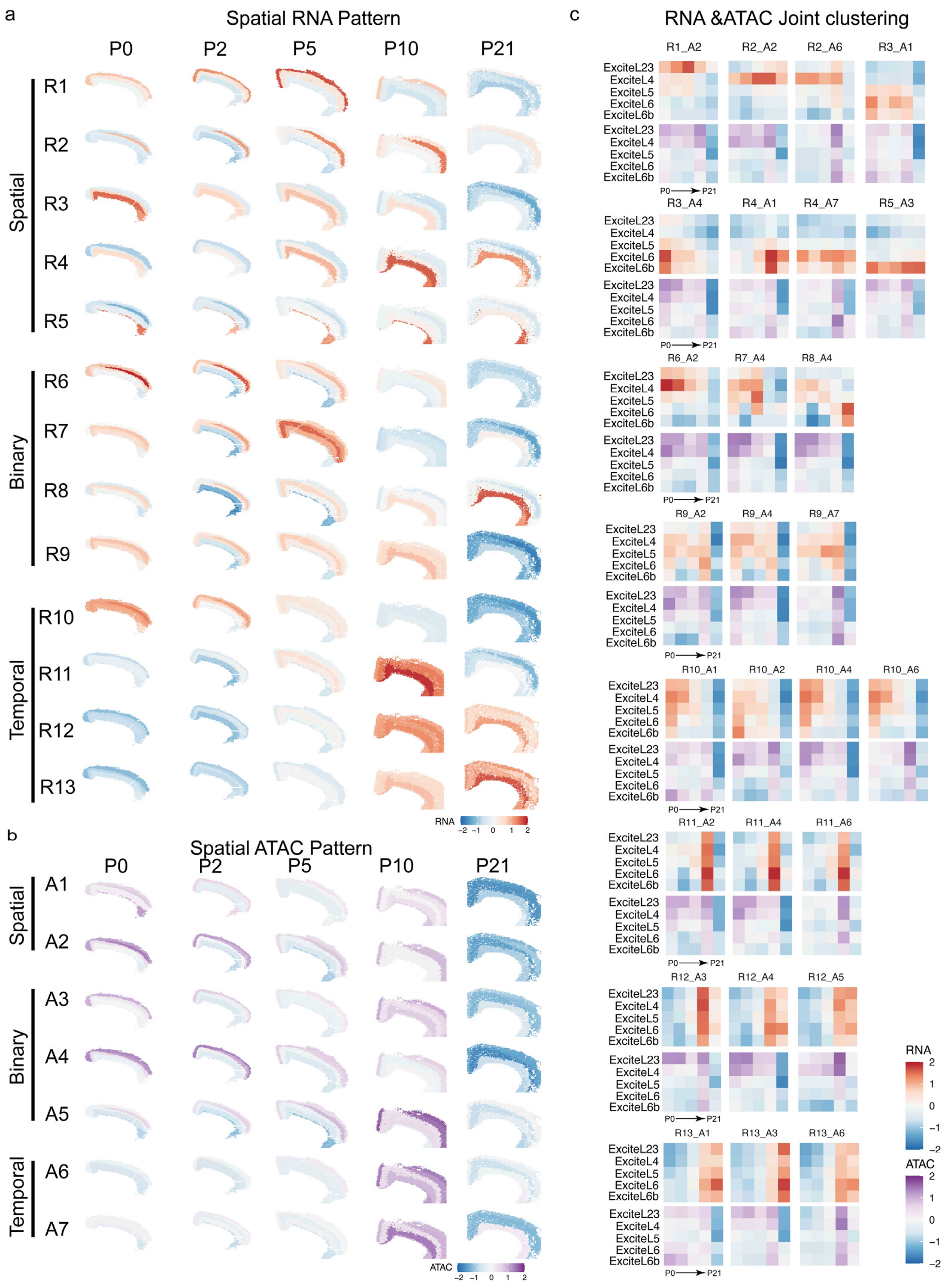


**Extended Data Fig. 10 Further analysis of developing mouse brain cortical layers. a-b,** Spatial RNA **(a)** and ATAC **(b)** patterns generated from the regression model. **c,** Heatmap for each of the 27 clusters from the RNA & ATAC joint clustering analysis.

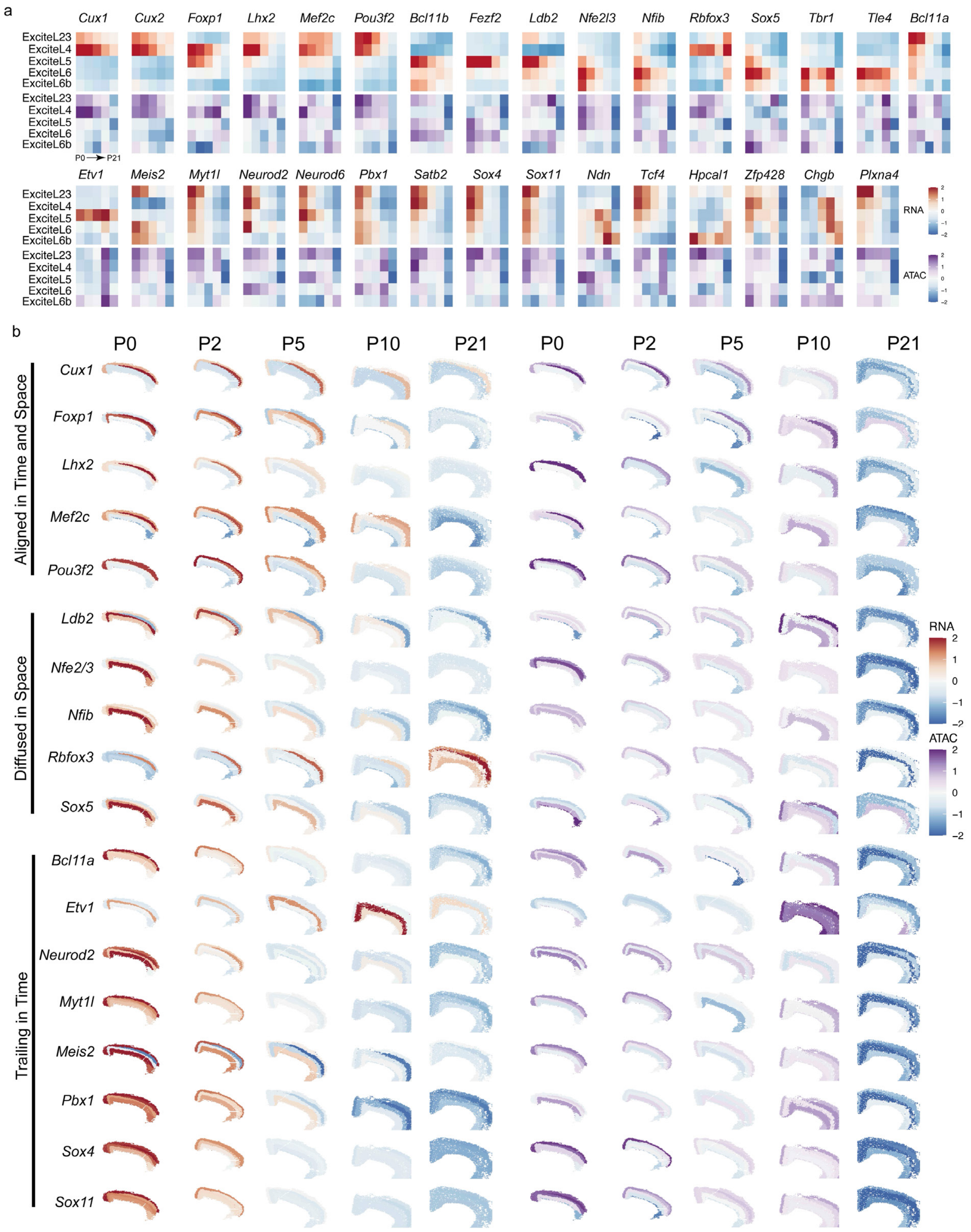


**Extended Data Fig. 11 Further analysis of developing mouse brain cortical layers. a,** Heatmaps of the RNA gene expression (top) and ATAC GAS (bottom) calculated on the basis of the regression model for specific neuronal genes. **b,** The RNA gene expression and ATAC GAS calculated on the basis of the regression model for specific neuronal genes.

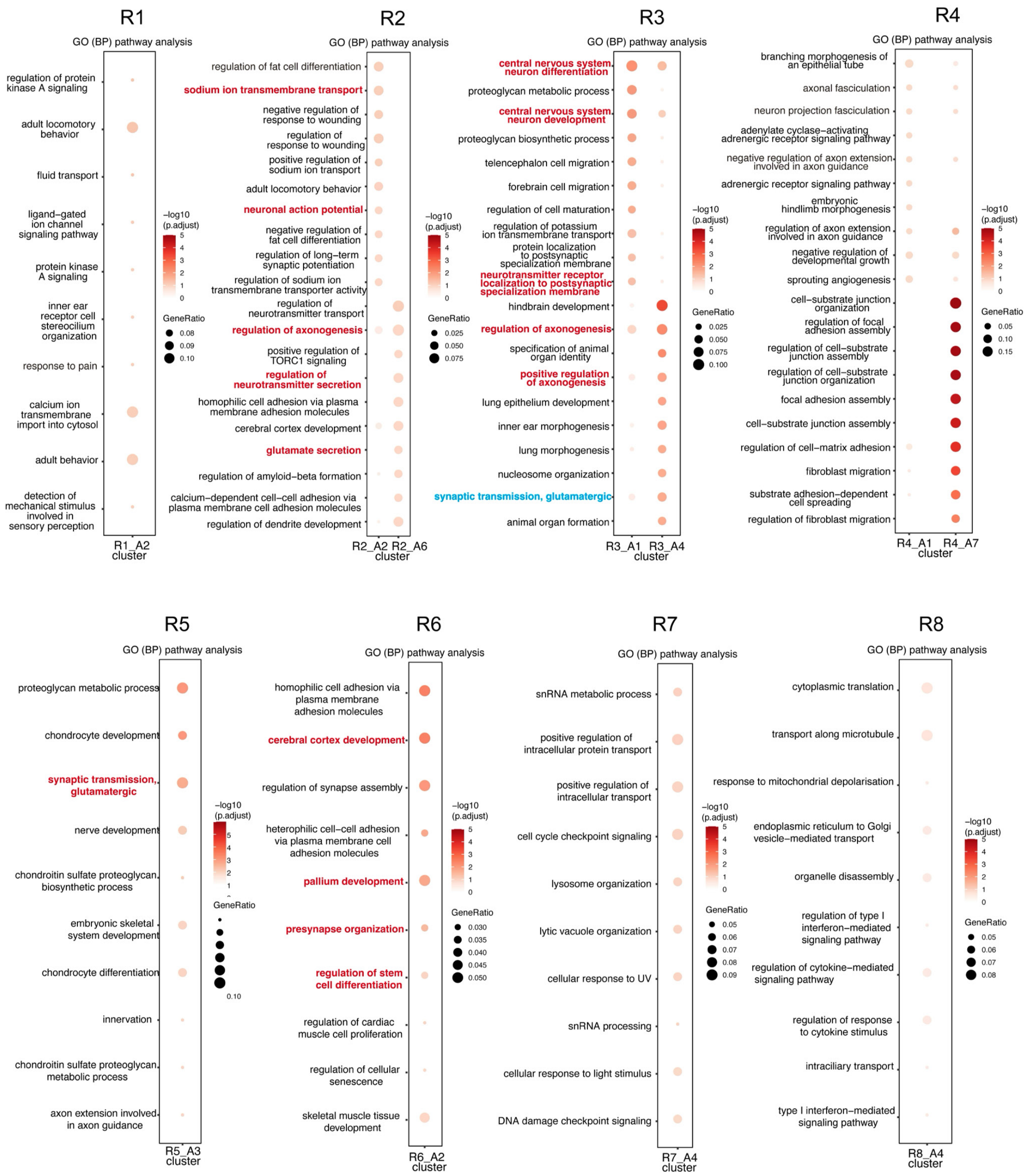


**Extended Data Fig. 12 Further analysis of developing mouse brain cortical layers.** GO analysis for each RNA cluster (R1-R8) generated from the regression model.

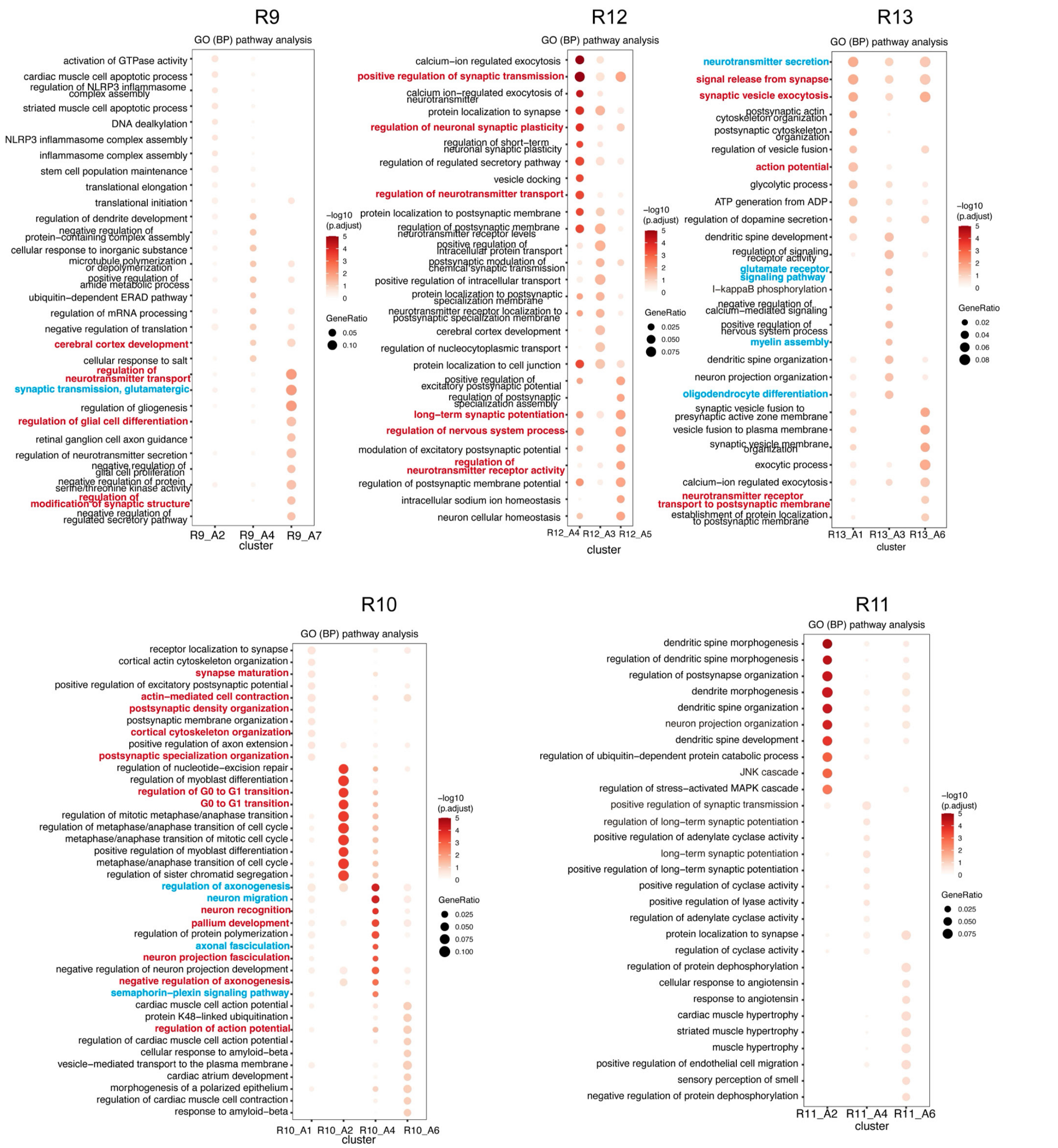


**Extended Data Fig. 13 Further analysis of developing mouse brain cortical layers.** GO analysis for each RNA cluster (R9-R13) generated from the regression model.

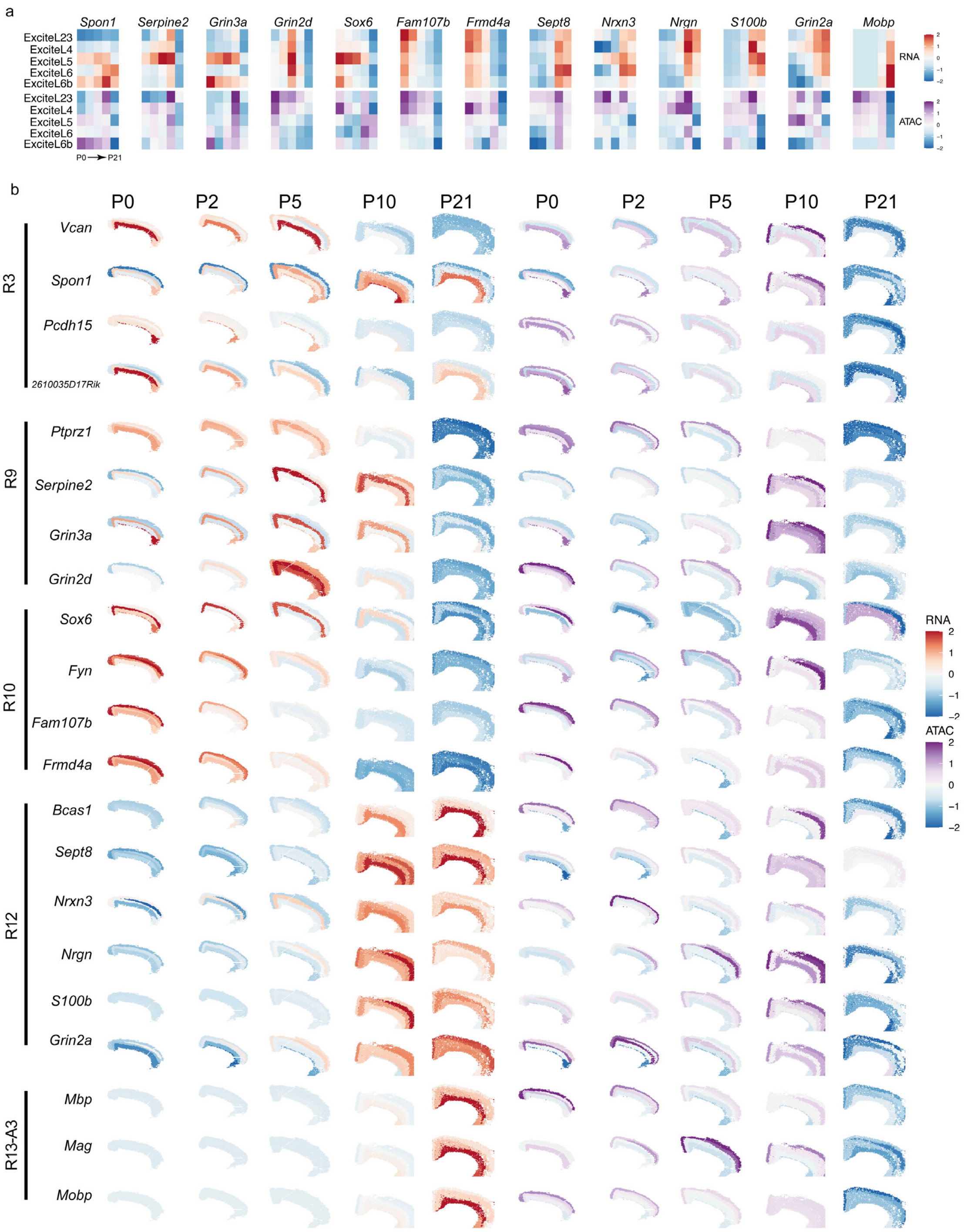


**Extended Data Fig. 14 Further analysis of developing mouse brain cortical layers. a,** Heatmaps of the RNA gene expression (top) and ATAC GAS (bottom) calculated on the basis of the regression model for myelin related genes. **b,** The RNA gene expression and ATAC GAS calculated on the basis of the regression model for myelin related genes.

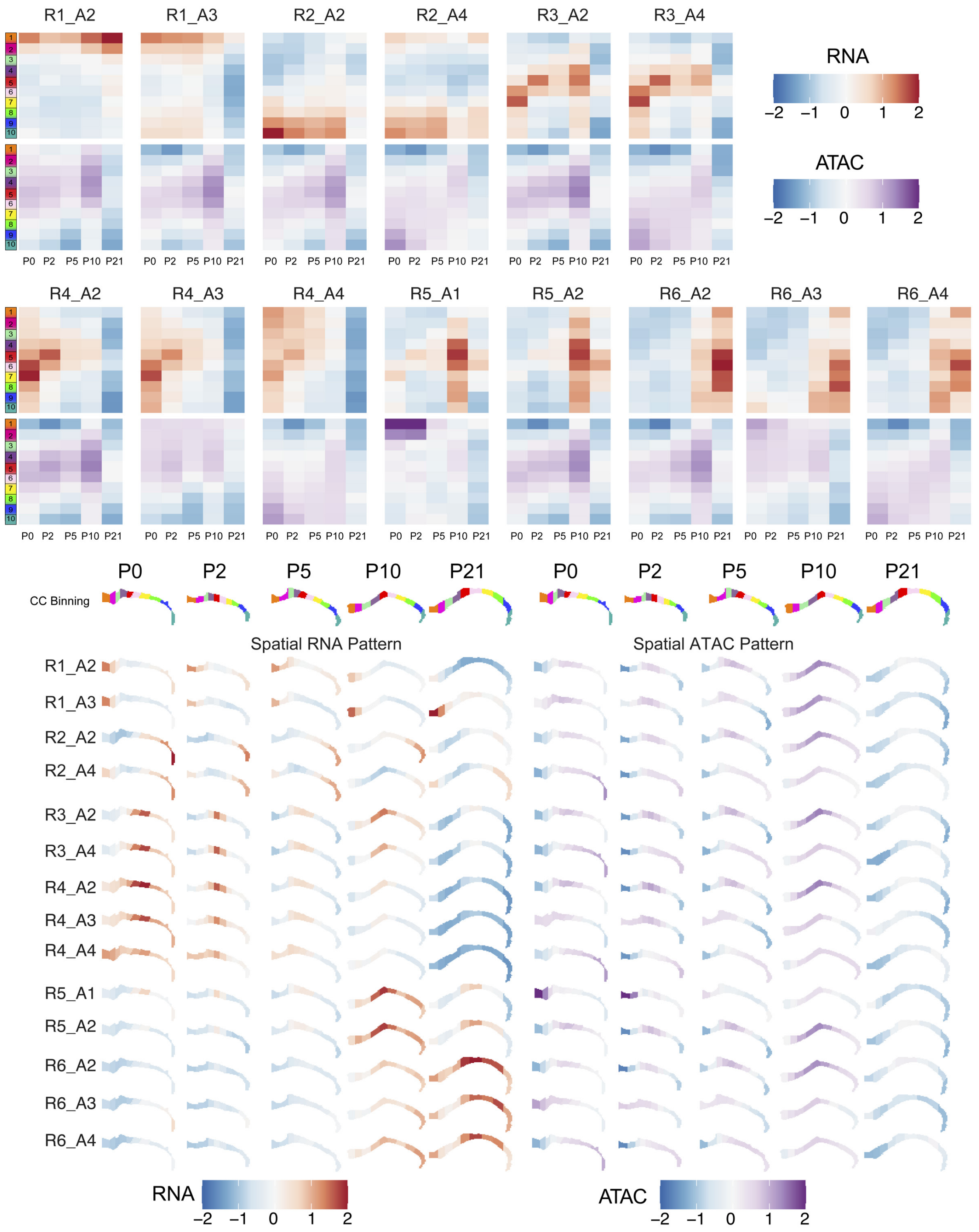


**Extended Data Fig. 15 Further analysis of developing mouse brain corpus callosum.** Heatmaps and spatial patterns of the 14 RNA&ATAC joint clustering.

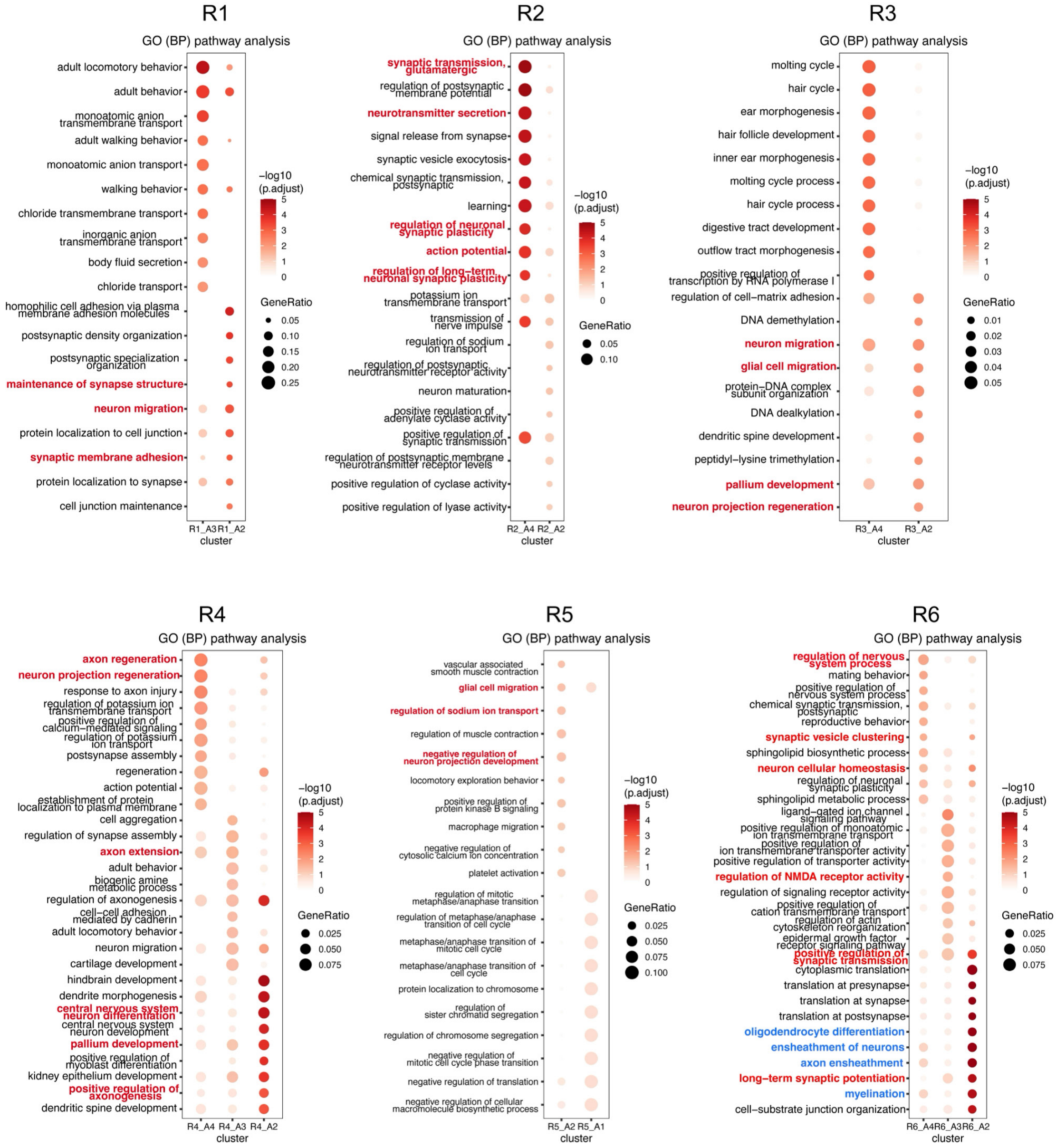


**Extended Data Fig. 16 Further analysis of developing mouse brain corpus callosum.** GO analysis for each RNA cluster (R1-R6) generated from the regression model.

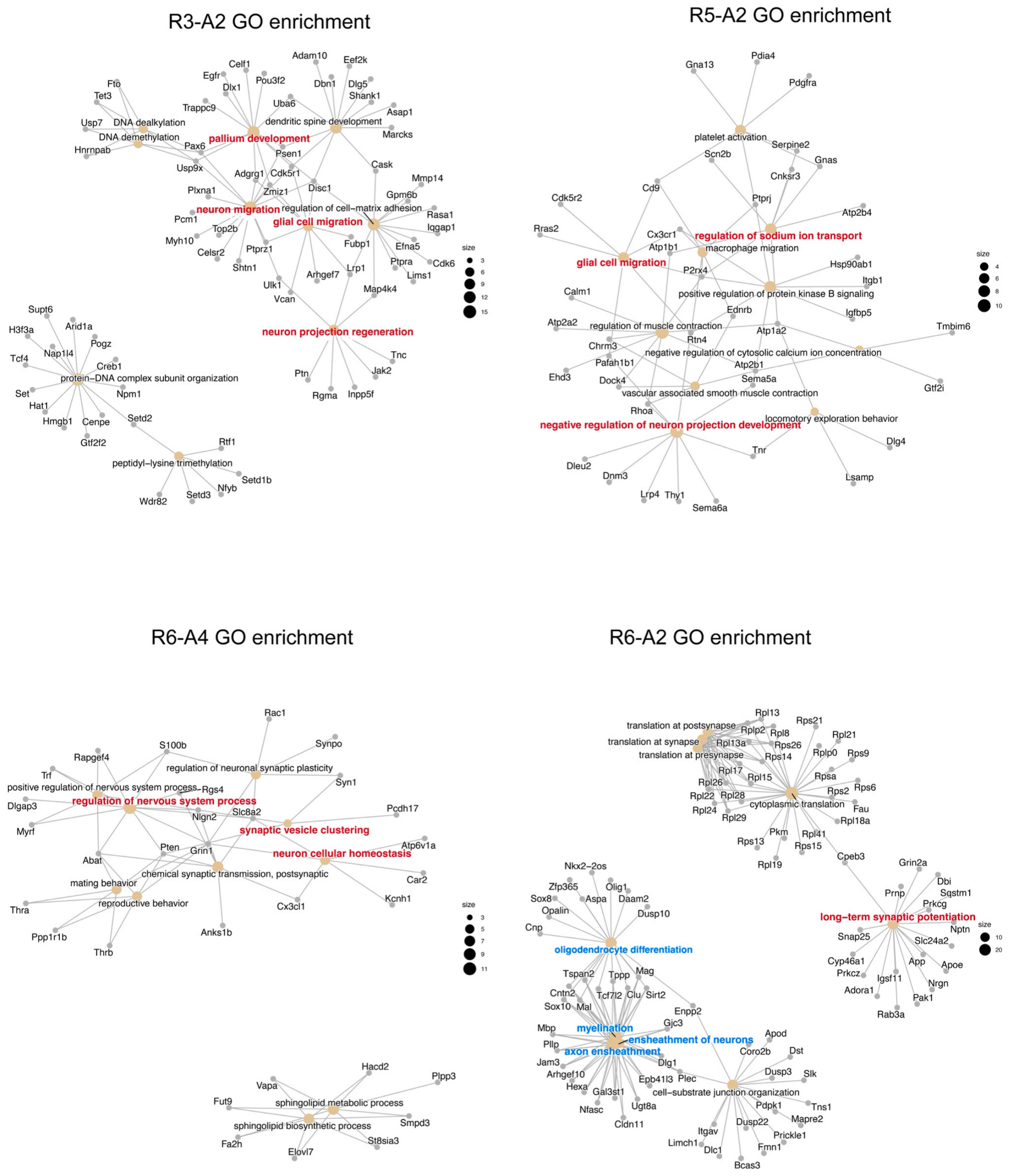


**Extended Data Fig. 17 Further analysis of developing mouse brain corpus callosum.** GO enrichment analysis for cluster R3-A2, R5-A2, R6-A4, and R6-A2.

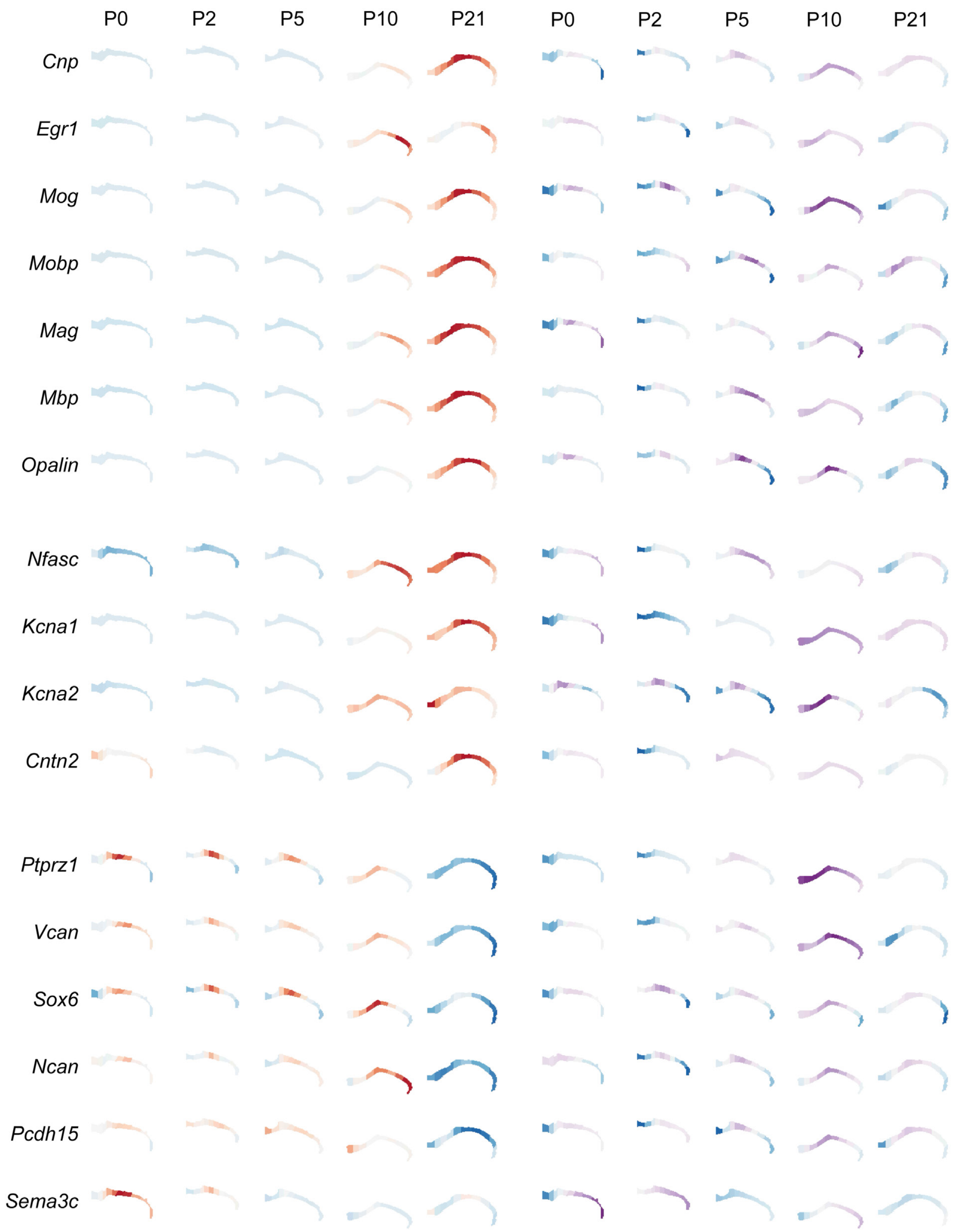


**Extended Data Fig. 18 Further analysis of developing mouse brain corpus callosum.** The RNA gene expression and ATAC GAS calculated on the basis of the regression model for specific genes.

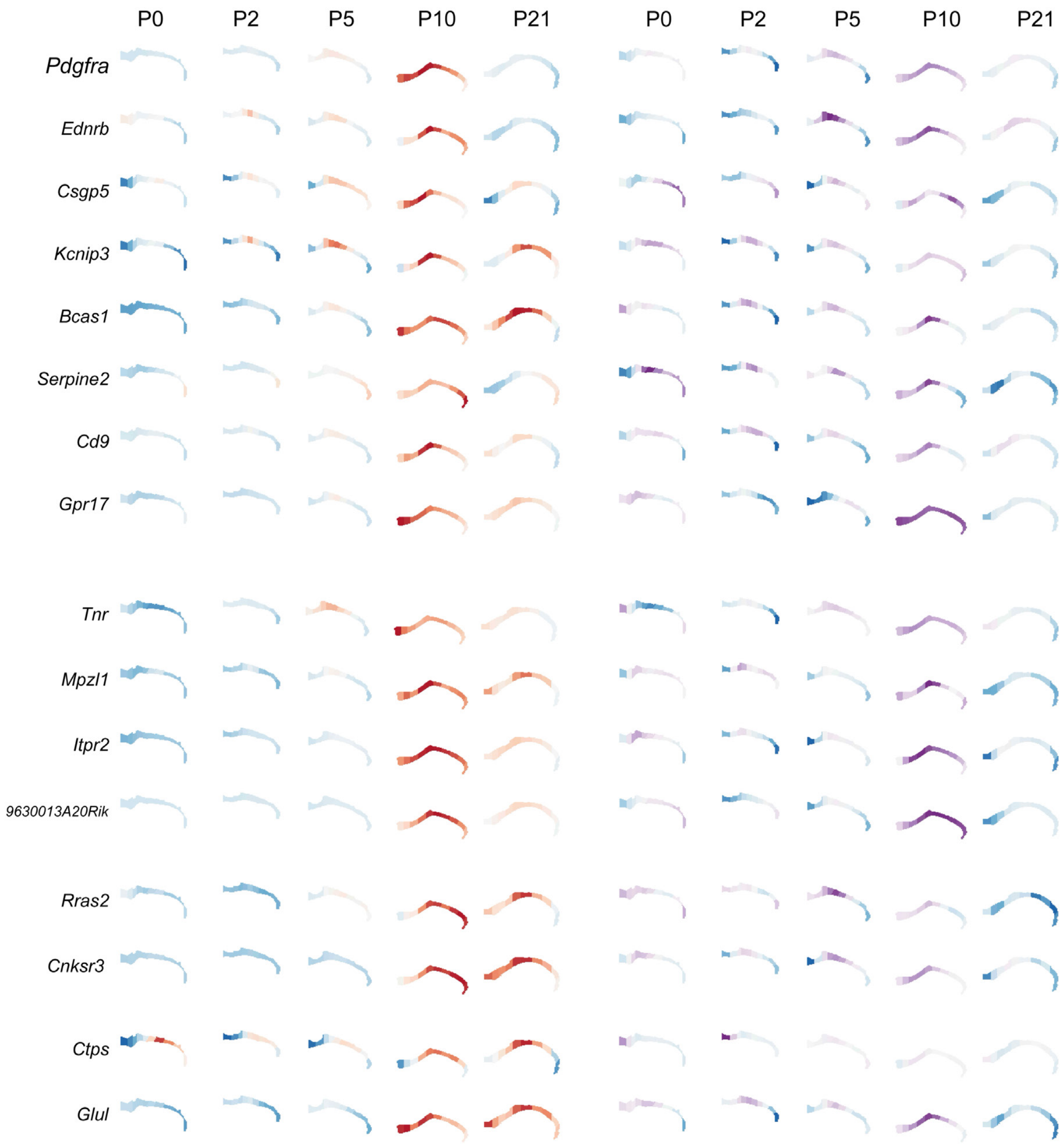


**Extended Data Fig. 19 Further analysis of developing mouse brain corpus callosum.** The RNA gene expression and ATAC GAS calculated on the basis of the regression model for specific genes.

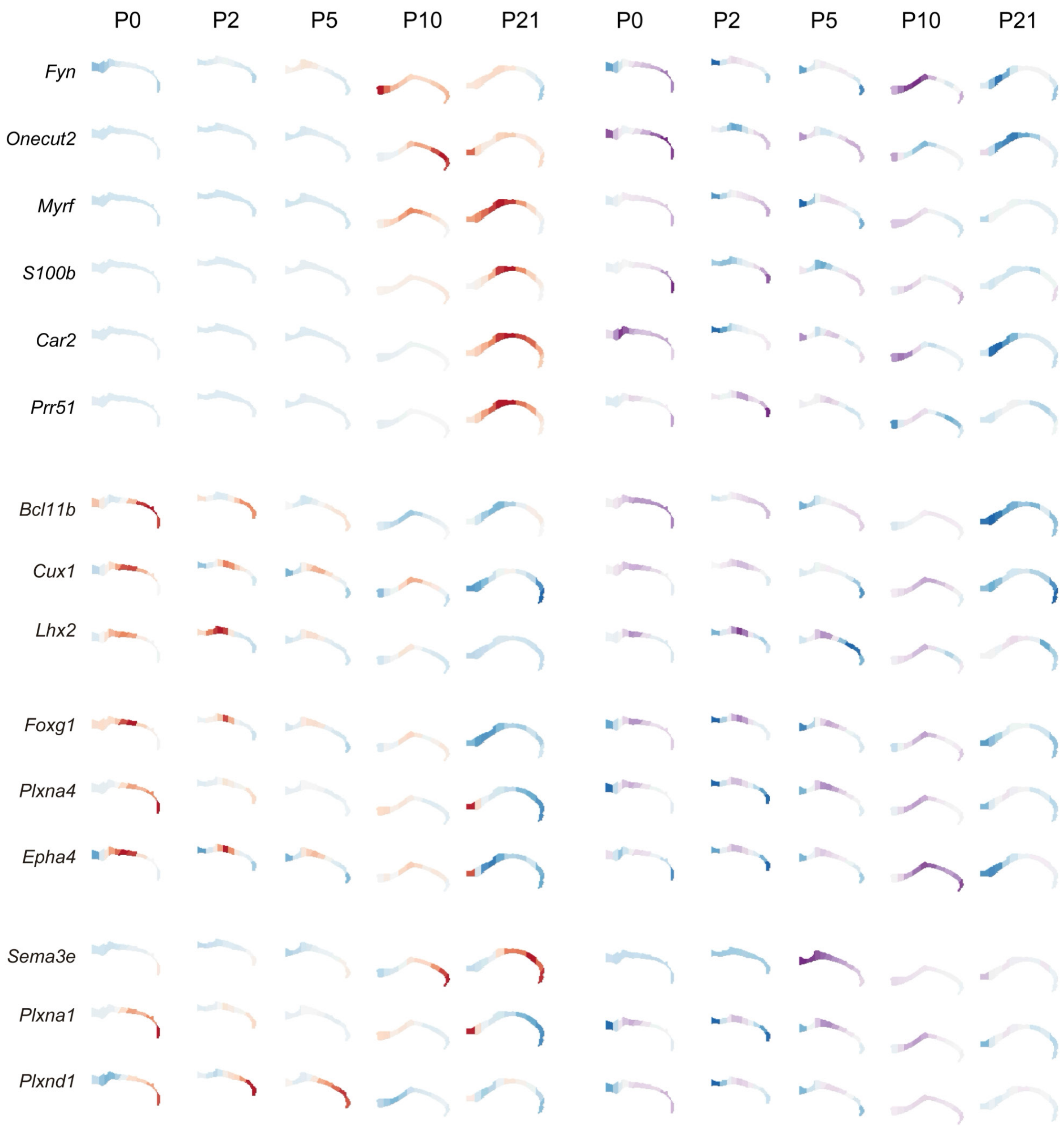


**Extended Data Fig. 20 Further analysis of developing mouse brain corpus callosum.** The RNA gene expression and ATAC GAS calculated on the basis of the regression model for specific genes.

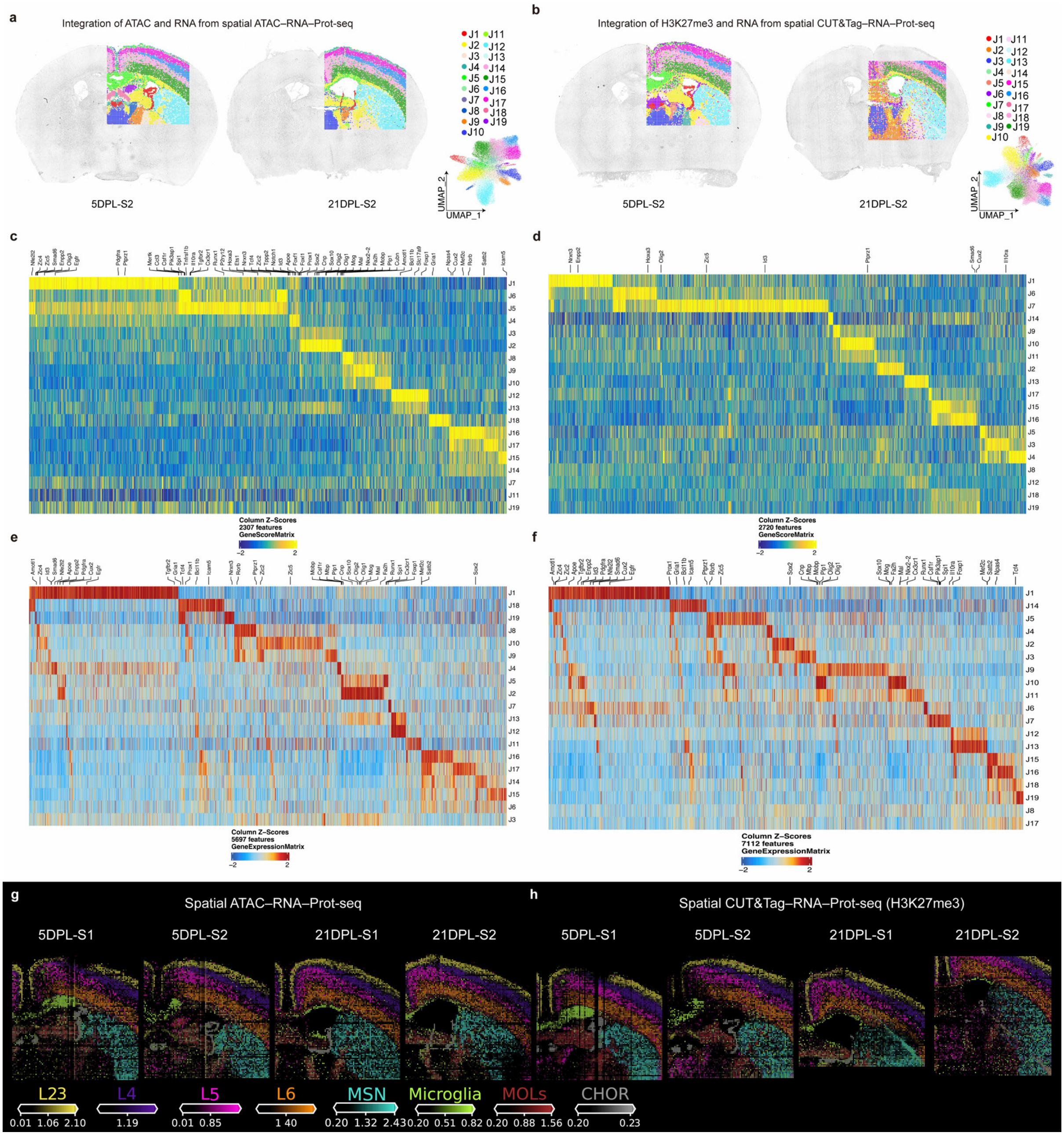


**Extended Data Fig. 21 Further analysis of spatial ATAC–RNA–Prot-seq (DBiT ARP-seq) and spatial CUT&Tag–RNA–Prot-seq (DBiT CTRP-seq, targeting H3K27me3) for LPC mouse model brains at 5 DPL and 21 DPL. a,** Integration of RNA and ATAC data in DBiT ARP-seq for replicate. **b,** Integration of H3K27me3 and RNA data in DBiT CTRP-seq for replicate. **c,** Marker GASs from each joint cluster in DBiT ARP-seq in **a** and **Fig. 4f**. **d,** Marker CSSs from each joint cluster in DBiT CTRP-seq in **b** and **Fig. 4g**. **e**, Marker gene expression from each joint cluster in DBiT ARP-seq in **a** and **Fig. 4f**. **f**, Marker gene expression from each joint cluster in DBiT CTRP-seq in **b** and **Fig. 4g**. **g-h,** Cell types predicted by cell2location from all processed LPC mouse model brains in DBiT ARP-seq (**g**) and DBiT CTRP-seq (**h**).

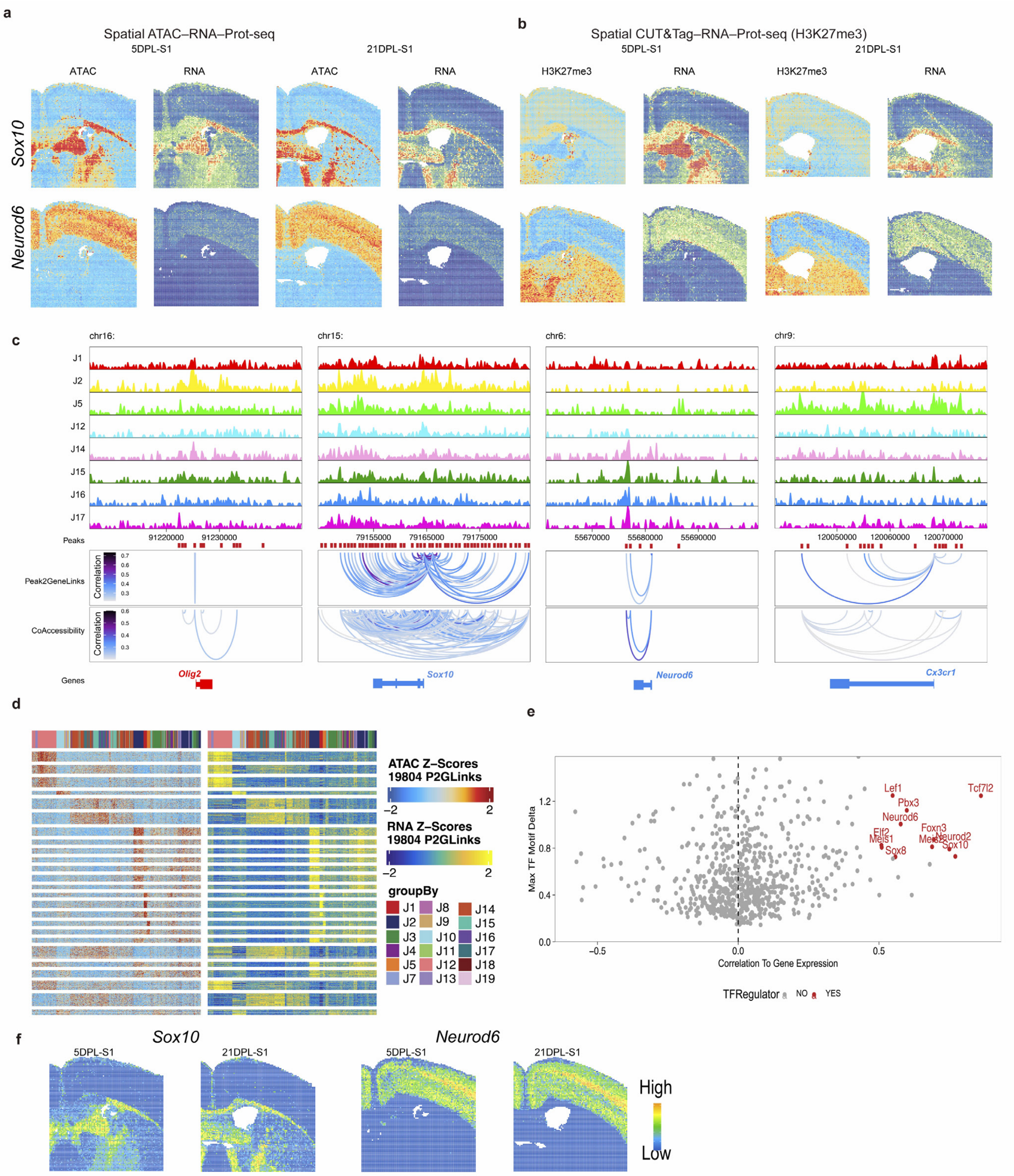


**Extended Data Fig. 22 Further analysis of spatial ATAC–RNA–Prot-seq (DBiT ARP-seq) and spatial CUT&Tag–RNA–Prot-seq (DBiT CTRP-seq, targeting H3K27me3) for LPC mouse model brains at 5 DPL and 21 DPL. a-b,** Spatial mapping of gene expression and GAS (**a**), or gene expression and CSS (**b**) for *Sox10* and *Neurod6* in both DBiT ARP-seq and DBiT CTRP-seq. **c,** Genome track visualization of marker genes with peak-to-gene links for distal regulatory elements and peak co-accessibility. **d,** Heatmaps of peak-to-gene links in DBiT ARP-seq. **e,** Dot plot showing the identification of positive TF regulators. **f,** Spatial mapping of deviation scores for selected TF motifs from DBiT ARP-seq.

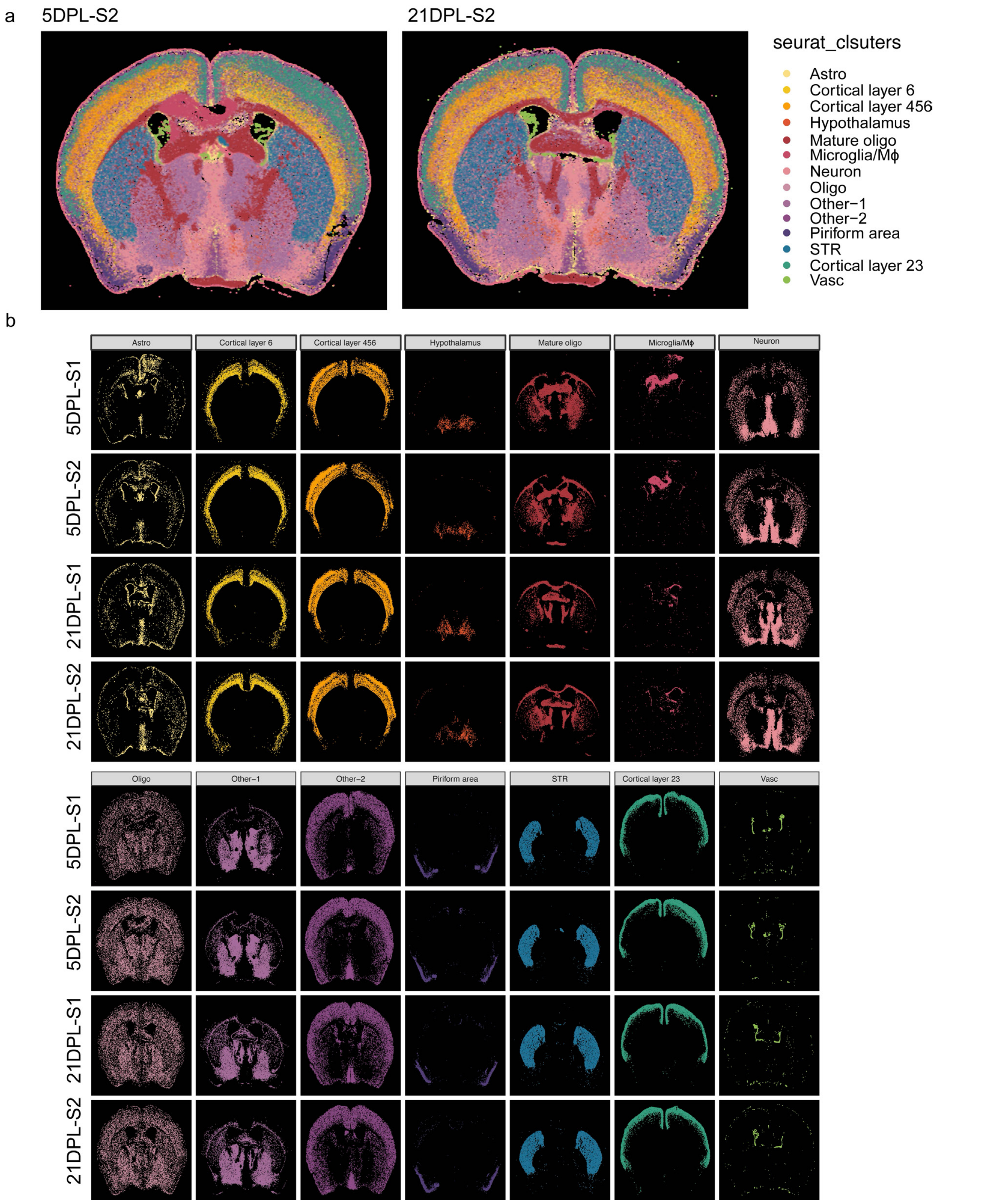


**Extended Data Fig. 23 Further analysis for CODEX images of LPC mouse model brains at 5 DPL and 21 DPL. a,** Seurat clustering of the CODEX images for replicates. **b,** Spatial map of the cell types from **a**.

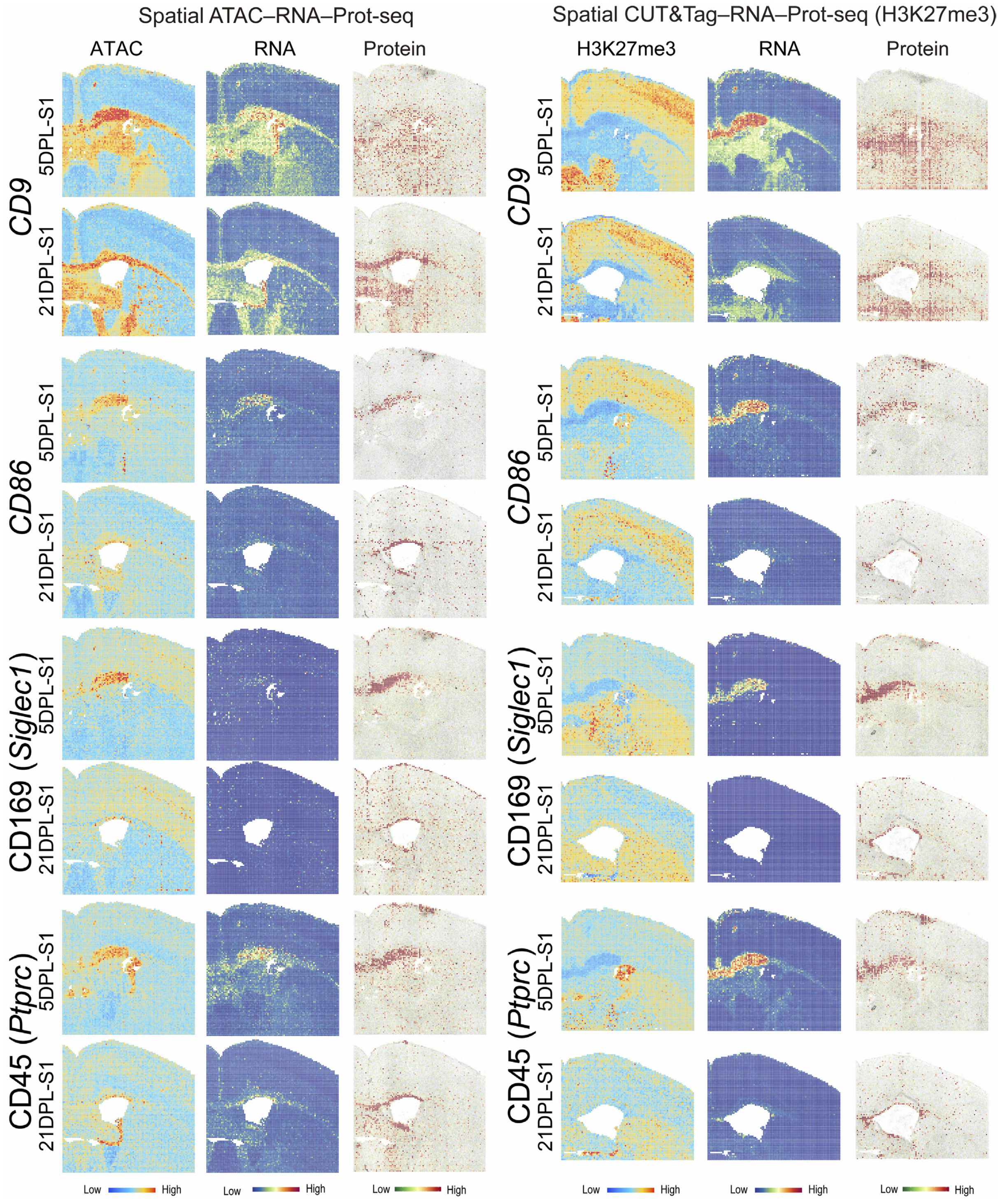


**Extended Data Fig. 24 Further analysis of spatial ATAC–RNA–Prot-seq (DBiT ARP-seq) and spatial CUT&Tag–RNA–Prot-seq (DBiT CTRP-seq, targeting H3K27me3) for LPC mouse model brains at 5 DPL and 21 DPL.** Spatial mapping of gene expression, GAS, CSS, and ADT protein expression for marker genes in both DBiT ARP-seq and DBiT CTRP-seq.

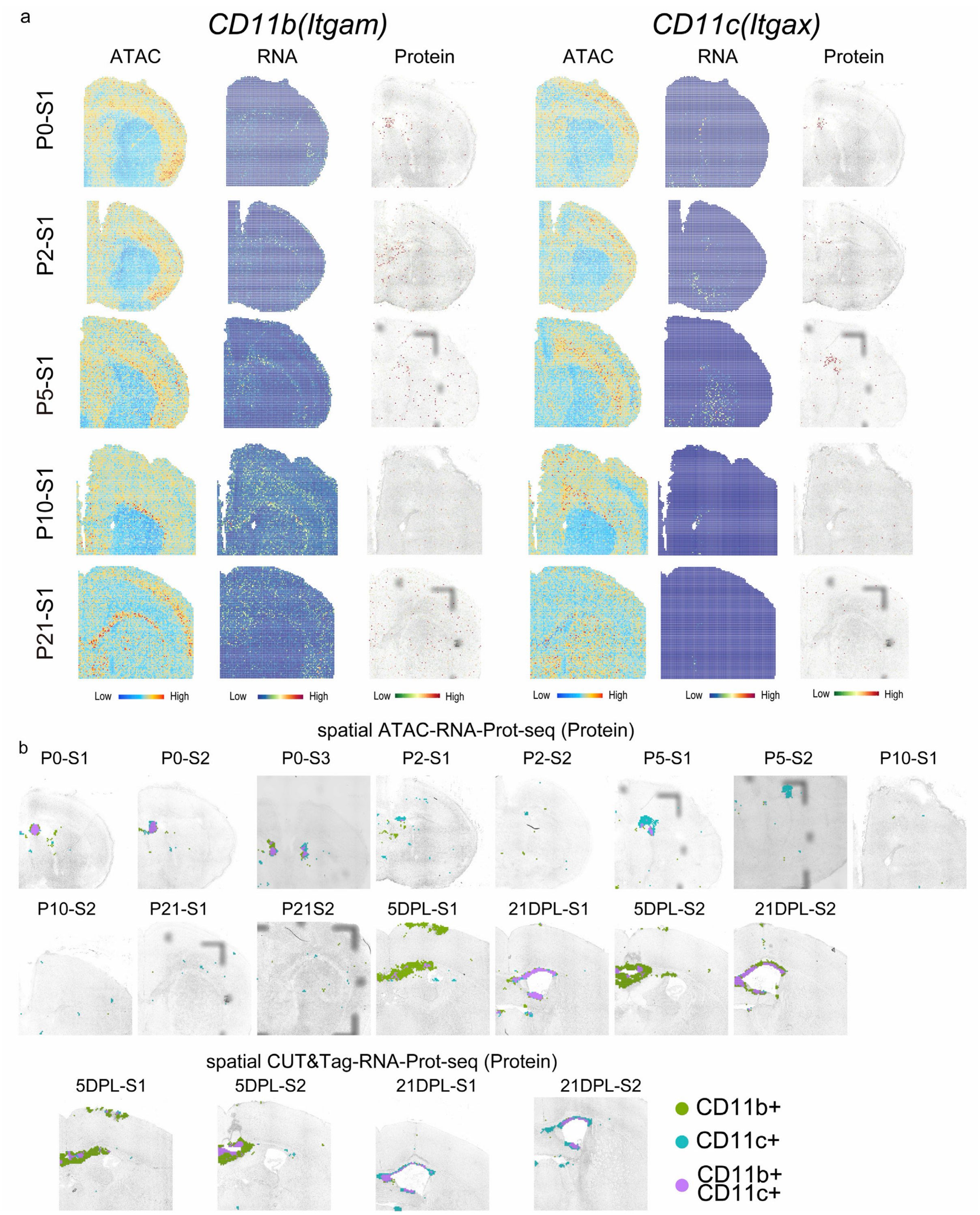


**Extended Data Fig. 25 Further analysis for microglia. a,** Spatial mapping of gene expression, GAS, and ADT protein expression for *Itgam* (CD11b*)* and *Itgax* (CD11c) in both DBiT ARP-seq and DBiT CTRP-seq. **b,** The ADT protein co-expression of CD11b and CD11c for developing and LPC mouse brains (including the replicate).

**Extended Data Table 1. DNA oligos used for transposome assembly, PCR, and preparation of sequencing library.**

| | |
|---|---|
| RT primer | /5Phos/CATCGGCGTACGACTNNNNNNNNNN/iBiodT/TTTTTTTTTTTTTTTVN |
| Ligation linker 1 | AGTCGTACGCCGATGCGAAACATCGGCCAC |
| Ligation linker 2 | CGAATGCTCTGGCCTCTCAAGCACGTGGAT |
| PCR Primer 1 | CAAGCGTTGGCTTCTCGCATCT |
| PCR Primer 2 | AAGCAGTGGTATCAACGCAGAGT |
| N501 | AATGATACGGCGACCACCGAGATCTACACTAGATCGCTCGTCGGCAGCGTCAGATGTGTATAAGAGACAG |
| N701 | CAAGCAGAAGACGGCATACGAGATTCGCCTTAGTCTCGTGGGCTCGGAGATGTGTATAAGAGACAGCAAGCGTTGGCTTCTCGCATCT |
| N702 | CAAGCAGAAGACGGCATACGAGATCTAGTACGGTCTCGTGGGCTCGGAGATGTGTATAAGAGACAGCAAGCGTTGGCTTCTCGCATCT |
| N703 | CAAGCAGAAGACGGCATACGAGATTTCTGCCTGTCTCGTGGGCTCGGAGATGTGTATAAGAGACAGCAAGCGTTGGCTTCTCGCATCT |
| N704 | CAAGCAGAAGACGGCATACGAGATGCTCAGGAGTCTCGTGGGCTCGGAGATGTGTATAAGAGACAGCAAGCGTTGGCTTCTCGCATCT |
| N705 | CAAGCAGAAGACGGCATACGAGATAGGAGTCCGTCTCGTGGGCTCGGAGATGTGTATAAGAGACAGCAAGCGTTGGCTTCTCGCATCT |
| N706 | CAAGCAGAAGACGGCATACGAGATCATGCCTAGTCTCGTGGGCTCGGAGATGTGTATAAGAGACAGCAAGCGTTGGCTTCTCGCATCT |
| N707 | CAAGCAGAAGACGGCATACGAGATGTAGAGAGGTCTCGTGGGCTCGGAGATGTGTATAAGAGACAGCAAGCGTTGGCTTCTCGCATCT |
| Tn5ME-A | 5′-TCGTCGGCAGCGTCAGATGTGTATAAGAGACAG-3′ |
| Tn5MErev | 5′-/5Phos/CTGTCTCTTATACACATCT-3′ |
| Tn5ME-B | 5′-/5Phos/CATCGGCGTACGACTAGATGTGTATAAGAGACAG-3′ |

**Extended Data Table 2. DNA barcode A sequences.**

| Barcode A | Sequence |
|---|---|
| Barcode A-1 | /5Phos/AGGCCAGAGCATTCGAACGTGATGTGGCCGATGTTTCG |
| Barcode A-2 | /5Phos/AGGCCAGAGCATTCGAAACATCGGTGGCCGATGTTTCG |
| Barcode A-3 | /5Phos/AGGCCAGAGCATTCGATGCCTAAGTGGCCGATGTTTCG |
| Barcode A-4 | /5Phos/AGGCCAGAGCATTCGAGTGGTCAGTGGCCGATGTTTCG |
| Barcode A-5 | /5Phos/AGGCCAGAGCATTCGACCACTGTGTGGCCGATGTTTCG |
| Barcode A-6 | /5Phos/AGGCCAGAGCATTCGACATTGGCGTGGCCGATGTTTCG |
| Barcode A-7 | /5Phos/AGGCCAGAGCATTCGCAGATCTGGTGGCCGATGTTTCG |
| Barcode A-8 | /5Phos/AGGCCAGAGCATTCGCATCAAGTGTGGCCGATGTTTCG |
| Barcode A-9 | /5Phos/AGGCCAGAGCATTCGCGCTGATCGTGGCCGATGTTTCG |
| Barcode A-10 | /5Phos/AGGCCAGAGCATTCGACAAGCTAGTGGCCGATGTTTCG |
| Barcode A-11 | /5Phos/AGGCCAGAGCATTCGCTGTAGCCGTGGCCGATGTTTCG |
| Barcode A-12 | /5Phos/AGGCCAGAGCATTCGAGTACAAGGTGGCCGATGTTTCG |
| Barcode A-13 | /5Phos/AGGCCAGAGCATTCGAACAACCAGTGGCCGATGTTTCG |
| Barcode A-14 | /5Phos/AGGCCAGAGCATTCGAACCGAGAGTGGCCGATGTTTCG |
| Barcode A-15 | /5Phos/AGGCCAGAGCATTCGAACGCTTAGTGGCCGATGTTTCG |
| Barcode A-16 | /5Phos/AGGCCAGAGCATTCGAAGACGGAGTGGCCGATGTTTCG |
| Barcode A-17 | /5Phos/AGGCCAGAGCATTCGAAGGTACAGTGGCCGATGTTTCG |
| Barcode A-18 | /5Phos/AGGCCAGAGCATTCGACACAGAAGTGGCCGATGTTTCG |
| Barcode A-19 | /5Phos/AGGCCAGAGCATTCGACAGCAGAGTGGCCGATGTTTCG |
| Barcode A-20 | /5Phos/AGGCCAGAGCATTCGACCTCCAAGTGGCCGATGTTTCG |
| Barcode A-21 | /5Phos/AGGCCAGAGCATTCGACGCTCGAGTGGCCGATGTTTCG |
| Barcode A-22 | /5Phos/AGGCCAGAGCATTCGACGTATCAGTGGCCGATGTTTCG |
| Barcode A-23 | /5Phos/AGGCCAGAGCATTCGACTATGCAGTGGCCGATGTTTCG |
| Barcode A-24 | /5Phos/AGGCCAGAGCATTCGAGAGTCAAGTGGCCGATGTTTCG |
| Barcode A-25 | /5Phos/AGGCCAGAGCATTCGAGATCGCAGTGGCCGATGTTTCG |
| Barcode A-26 | /5Phos/AGGCCAGAGCATTCGAGCAGGAAGTGGCCGATGTTTCG |
| Barcode A-27 | /5Phos/AGGCCAGAGCATTCGAGTCACTAGTGGCCGATGTTTCG |
| Barcode A-28 | /5Phos/AGGCCAGAGCATTCGATCCTGTAGTGGCCGATGTTTCG |
| Barcode A-29 | /5Phos/AGGCCAGAGCATTCGATTGAGGAGTGGCCGATGTTTCG |
| Barcode A-30 | /5Phos/AGGCCAGAGCATTCGCAACCACAGTGGCCGATGTTTCG |
| Barcode A-31 | /5Phos/AGGCCAGAGCATTCGGACTAGTAGTGGCCGATGTTTCG |
| Barcode A-32 | /5Phos/AGGCCAGAGCATTCGCAATGGAAGTGGCCGATGTTTCG |
| Barcode A-33 | /5Phos/AGGCCAGAGCATTCGCACTTCGAGTGGCCGATGTTTCG |
| Barcode A-34 | /5Phos/AGGCCAGAGCATTCGCAGCGTTAGTGGCCGATGTTTCG |
| Barcode A-35 | /5Phos/AGGCCAGAGCATTCGCATACCAAGTGGCCGATGTTTCG |

| | |
|---|---|
| Barcode A-36 | /5Phos/AGGCCAGAGCATTCGCCAGTTCAGTGGCCGATGTTTCG |
| Barcode A-37 | /5Phos/AGGCCAGAGCATTCGCCGAAGTAGTGGCCGATGTTTCG |
| Barcode A-38 | /5Phos/AGGCCAGAGCATTCGCCGTGAGAGTGGCCGATGTTTCG |
| Barcode A-39 | /5Phos/AGGCCAGAGCATTCGCCTCCTGAGTGGCCGATGTTTCG |
| Barcode A-40 | /5Phos/AGGCCAGAGCATTCGCGAACTTAGTGGCCGATGTTTCG |
| Barcode A-41 | /5Phos/AGGCCAGAGCATTCGCGACTGGAGTGGCCGATGTTTCG |
| Barcode A-42 | /5Phos/AGGCCAGAGCATTCGCGCATACAGTGGCCGATGTTTCG |
| Barcode A-43 | /5Phos/AGGCCAGAGCATTCGCTCAATGAGTGGCCGATGTTTCG |
| Barcode A-44 | /5Phos/AGGCCAGAGCATTCGCTGAGCCAGTGGCCGATGTTTCG |
| Barcode A-45 | /5Phos/AGGCCAGAGCATTCGCTGGCATAGTGGCCGATGTTTCG |
| Barcode A-46 | /5Phos/AGGCCAGAGCATTCGGAATCTGAGTGGCCGATGTTTCG |
| Barcode A-47 | /5Phos/AGGCCAGAGCATTCGCAAGACTAGTGGCCGATGTTTCG |
| Barcode A-48 | /5Phos/AGGCCAGAGCATTCGGAGCTGAAGTGGCCGATGTTTCG |
| Barcode A-49 | /5Phos/AGGCCAGAGCATTCGGATAGACAGTGGCCGATGTTTCG |
| Barcode A-50 | /5Phos/AGGCCAGAGCATTCGGCCACATAGTGGCCGATGTTTCG |
| Barcode A-51 | /5Phos/AGGCCAGAGCATTCGGCGAGTAAGTGGCCGATGTTTCG |
| Barcode A-52 | /5Phos/AGGCCAGAGCATTCGGCTAACGAGTGGCCGATGTTTCG |
| Barcode A-53 | /5Phos/AGGCCAGAGCATTCGGCTCGGTAGTGGCCGATGTTTCG |
| Barcode A-54 | /5Phos/AGGCCAGAGCATTCGGGAGAACAGTGGCCGATGTTTCG |
| Barcode A-55 | /5Phos/AGGCCAGAGCATTCGGGTGCGAAGTGGCCGATGTTTCG |
| Barcode A-56 | /5Phos/AGGCCAGAGCATTCGGTACGCAAGTGGCCGATGTTTCG |
| Barcode A-57 | /5Phos/AGGCCAGAGCATTCGGTCGTAGAGTGGCCGATGTTTCG |
| Barcode A-58 | /5Phos/AGGCCAGAGCATTCGGTCTGTCAGTGGCCGATGTTTCG |
| Barcode A-59 | /5Phos/AGGCCAGAGCATTCGGTGTTCTAGTGGCCGATGTTTCG |
| Barcode A-60 | /5Phos/AGGCCAGAGCATTCGTAGGATGAGTGGCCGATGTTTCG |
| Barcode A-61 | /5Phos/AGGCCAGAGCATTCGTATCAGCAGTGGCCGATGTTTCG |
| Barcode A-62 | /5Phos/AGGCCAGAGCATTCGTCCGTCTAGTGGCCGATGTTTCG |
| Barcode A-63 | /5Phos/AGGCCAGAGCATTCGTCTTCACAGTGGCCGATGTTTCG |
| Barcode A-64 | /5Phos/AGGCCAGAGCATTCGTGAAGAGAGTGGCCGATGTTTCG |
| Barcode A-65 | /5Phos/AGGCCAGAGCATTCGTGGAACAAGTGGCCGATGTTTCG |
| Barcode A-66 | /5Phos/AGGCCAGAGCATTCGTGGCTTCAGTGGCCGATGTTTCG |
| Barcode A-67 | /5Phos/AGGCCAGAGCATTCGTGGTGGTAGTGGCCGATGTTTCG |
| Barcode A-68 | /5Phos/AGGCCAGAGCATTCGTTCACGCAGTGGCCGATGTTTCG |
| Barcode A-69 | /5Phos/AGGCCAGAGCATTCGAACTCACCGTGGCCGATGTTTCG |
| Barcode A-70 | /5Phos/AGGCCAGAGCATTCGAAGAGATCGTGGCCGATGTTTCG |
| Barcode A-71 | /5Phos/AGGCCAGAGCATTCGAAGGACACGTGGCCGATGTTTCG |
| Barcode A-72 | /5Phos/AGGCCAGAGCATTCGAATCCGTCGTGGCCGATGTTTCG |

| | |
|---|---|
| Barcode A-73 | /5Phos/AGGCCAGAGCATTCGAATGTTGCGTGGCCGATGTTTCG |
| Barcode A-74 | /5Phos/AGGCCAGAGCATTCGACACGACCGTGGCCGATGTTTCG |
| Barcode A-75 | /5Phos/AGGCCAGAGCATTCGACAGATTCGTGGCCGATGTTTCG |
| Barcode A-76 | /5Phos/AGGCCAGAGCATTCGAGATGTACGTGGCCGATGTTTCG |
| Barcode A-77 | /5Phos/AGGCCAGAGCATTCGAGCACCTCGTGGCCGATGTTTCG |
| Barcode A-78 | /5Phos/AGGCCAGAGCATTCGAGCCATGCGTGGCCGATGTTTCG |
| Barcode A-79 | /5Phos/AGGCCAGAGCATTCGAGGCTAACGTGGCCGATGTTTCG |
| Barcode A-80 | /5Phos/AGGCCAGAGCATTCGATAGCGACGTGGCCGATGTTTCG |
| Barcode A-81 | /5Phos/AGGCCAGAGCATTCGATCATTCCGTGGCCGATGTTTCG |
| Barcode A-82 | /5Phos/AGGCCAGAGCATTCGATTGGCTCGTGGCCGATGTTTCG |
| Barcode A-83 | /5Phos/AGGCCAGAGCATTCGCAAGGAGCGTGGCCGATGTTTCG |
| Barcode A-84 | /5Phos/AGGCCAGAGCATTCGCACCTTACGTGGCCGATGTTTCG |
| Barcode A-85 | /5Phos/AGGCCAGAGCATTCGCCATCCTCGTGGCCGATGTTTCG |
| Barcode A-86 | /5Phos/AGGCCAGAGCATTCGCCGACAACGTGGCCGATGTTTCG |
| Barcode A-87 | /5Phos/AGGCCAGAGCATTCGCCTAATCCGTGGCCGATGTTTCG |
| Barcode A-88 | /5Phos/AGGCCAGAGCATTCGCCTCTATCGTGGCCGATGTTTCG |
| Barcode A-89 | /5Phos/AGGCCAGAGCATTCGCGACACACGTGGCCGATGTTTCG |
| Barcode A-90 | /5Phos/AGGCCAGAGCATTCGCGGATTGCGTGGCCGATGTTTCG |
| Barcode A-91 | /5Phos/AGGCCAGAGCATTCGCTAAGGTCGTGGCCGATGTTTCG |
| Barcode A-92 | /5Phos/AGGCCAGAGCATTCGGAACAGGCGTGGCCGATGTTTCG |
| Barcode A-93 | /5Phos/AGGCCAGAGCATTCGGACAGTGCGTGGCCGATGTTTCG |
| Barcode A-94 | /5Phos/AGGCCAGAGCATTCGGAGTTAGCGTGGCCGATGTTTCG |
| Barcode A-95 | /5Phos/AGGCCAGAGCATTCGGATGAATCGTGGCCGATGTTTCG |
| Barcode A-96 | /5Phos/AGGCCAGAGCATTCGGCCAAGACGTGGCCGATGTTTCG |
| Barcode A-97 | /5Phos/AGGCCAGAGCATTCGCGGAAGAAGTGGCCGATGTTTCG |
| Barcode A-98 | /5Phos/AGGCCAGAGCATTCGGTGACAAGGTGGCCGATGTTTCG |
| Barcode A-99 | /5Phos/AGGCCAGAGCATTCGGAACCAGAGTGGCCGATGTTTCG |
| Barcode A-100 | /5Phos/AGGCCAGAGCATTCGTTGCTGGAGTGGCCGATGTTTCG |

**Extended Data Table 3. DNA barcode B sequences.**

| **Barcode B** | **Sequence** |
|---|---|
| Barcode B-1 | CAAGCGTTGGCTTCTCGCATCTAACGTGATATCCACGTGCTTGAG |
| Barcode B-2 | CAAGCGTTGGCTTCTCGCATCTAAACATCGATCCACGTGCTTGAG |
| Barcode B-3 | CAAGCGTTGGCTTCTCGCATCTATGCCTAAATCCACGTGCTTGAG |
| Barcode B-4 | CAAGCGTTGGCTTCTCGCATCTAGTGGTCAATCCACGTGCTTGAG |
| Barcode B-5 | CAAGCGTTGGCTTCTCGCATCTACCACTGTATCCACGTGCTTGAG |
| Barcode B-6 | CAAGCGTTGGCTTCTCGCATCTACATTGGCATCCACGTGCTTGAG |
| Barcode B-7 | CAAGCGTTGGCTTCTCGCATCTCAGATCTGATCCACGTGCTTGAG |
| Barcode B-8 | CAAGCGTTGGCTTCTCGCATCTCATCAAGTATCCACGTGCTTGAG |
| Barcode B-9 | CAAGCGTTGGCTTCTCGCATCTCGCTGATCATCCACGTGCTTGAG |
| Barcode B-10 | CAAGCGTTGGCTTCTCGCATCTACAAGCTAATCCACGTGCTTGAG |
| Barcode B-11 | CAAGCGTTGGCTTCTCGCATCTCTGTAGCCATCCACGTGCTTGAG |
| Barcode B-12 | CAAGCGTTGGCTTCTCGCATCTAGTACAAGATCCACGTGCTTGAG |
| Barcode B-13 | CAAGCGTTGGCTTCTCGCATCTAACAACCAATCCACGTGCTTGAG |
| Barcode B-14 | CAAGCGTTGGCTTCTCGCATCTAACCGAGAATCCACGTGCTTGAG |
| Barcode B-15 | CAAGCGTTGGCTTCTCGCATCTAACGCTTAATCCACGTGCTTGAG |
| Barcode B-16 | CAAGCGTTGGCTTCTCGCATCTAAGACGGAATCCACGTGCTTGAG |
| Barcode B-17 | CAAGCGTTGGCTTCTCGCATCTAAGGTACAATCCACGTGCTTGAG |
| Barcode B-18 | CAAGCGTTGGCTTCTCGCATCTACACAGAAATCCACGTGCTTGAG |
| Barcode B-19 | CAAGCGTTGGCTTCTCGCATCTACAGCAGAATCCACGTGCTTGAG |
| Barcode B-20 | CAAGCGTTGGCTTCTCGCATCTACCTCCAAATCCACGTGCTTGAG |
| Barcode B-21 | CAAGCGTTGGCTTCTCGCATCTACGCTCGAATCCACGTGCTTGAG |
| Barcode B-22 | CAAGCGTTGGCTTCTCGCATCTACGTATCAATCCACGTGCTTGAG |
| Barcode B-23 | CAAGCGTTGGCTTCTCGCATCTACTATGCAATCCACGTGCTTGAG |
| Barcode B-24 | CAAGCGTTGGCTTCTCGCATCTAGAGTCAAATCCACGTGCTTGAG |
| Barcode B-25 | CAAGCGTTGGCTTCTCGCATCTAGATCGCAATCCACGTGCTTGAG |
| Barcode B-26 | CAAGCGTTGGCTTCTCGCATCTAGCAGGAAATCCACGTGCTTGAG |
| Barcode B-27 | CAAGCGTTGGCTTCTCGCATCTAGTCACTAATCCACGTGCTTGAG |
| Barcode B-28 | CAAGCGTTGGCTTCTCGCATCTATCCTGTAATCCACGTGCTTGAG |
| Barcode B-29 | CAAGCGTTGGCTTCTCGCATCTATTGAGGAATCCACGTGCTTGAG |
| Barcode B-30 | CAAGCGTTGGCTTCTCGCATCTCAACCACAATCCACGTGCTTGAG |
| Barcode B-31 | CAAGCGTTGGCTTCTCGCATCTGACTAGTAATCCACGTGCTTGAG |
| Barcode B-32 | CAAGCGTTGGCTTCTCGCATCTCAATGGAAATCCACGTGCTTGAG |
| Barcode B-33 | CAAGCGTTGGCTTCTCGCATCTCACTTCGAATCCACGTGCTTGAG |
| Barcode B-34 | CAAGCGTTGGCTTCTCGCATCTCAGCGTTAATCCACGTGCTTGAG |
| Barcode B-35 | CAAGCGTTGGCTTCTCGCATCTCATACCAAATCCACGTGCTTGAG |

| | |
|---|---|
| Barcode B-36 | CAAGCGTTGGCTTCTCGCATCTCCAGTTCAATCCACGTGCTTGAG |
| Barcode B-37 | CAAGCGTTGGCTTCTCGCATCTCCGAAGTAATCCACGTGCTTGAG |
| Barcode B-38 | CAAGCGTTGGCTTCTCGCATCTCCGTGAGAATCCACGTGCTTGAG |
| Barcode B-39 | CAAGCGTTGGCTTCTCGCATCTCCTCCTGAATCCACGTGCTTGAG |
| Barcode B-40 | CAAGCGTTGGCTTCTCGCATCTCGAACTTAATCCACGTGCTTGAG |
| Barcode B-41 | CAAGCGTTGGCTTCTCGCATCTCGACTGGAATCCACGTGCTTGAG |
| Barcode B-42 | CAAGCGTTGGCTTCTCGCATCTCGCATACAATCCACGTGCTTGAG |
| Barcode B-43 | CAAGCGTTGGCTTCTCGCATCTCTCAATGAATCCACGTGCTTGAG |
| Barcode B-44 | CAAGCGTTGGCTTCTCGCATCTCTGAGCCAATCCACGTGCTTGAG |
| Barcode B-45 | CAAGCGTTGGCTTCTCGCATCTCTGGCATAATCCACGTGCTTGAG |
| Barcode B-46 | CAAGCGTTGGCTTCTCGCATCTGAATCTGAATCCACGTGCTTGAG |
| Barcode B-47 | CAAGCGTTGGCTTCTCGCATCTCAAGACTAATCCACGTGCTTGAG |
| Barcode B-48 | CAAGCGTTGGCTTCTCGCATCTGAGCTGAAATCCACGTGCTTGAG |
| Barcode B-49 | CAAGCGTTGGCTTCTCGCATCTGATAGACAATCCACGTGCTTGAG |
| Barcode B-50 | CAAGCGTTGGCTTCTCGCATCTGCCACATAATCCACGTGCTTGAG |
| Barcode B-51 | CAAGCGTTGGCTTCTCGCATCTGCGAGTAAATCCACGTGCTTGAG |
| Barcode B-52 | CAAGCGTTGGCTTCTCGCATCTGCTAACGAATCCACGTGCTTGAG |
| Barcode B-53 | CAAGCGTTGGCTTCTCGCATCTGCTCGGTAATCCACGTGCTTGAG |
| Barcode B-54 | CAAGCGTTGGCTTCTCGCATCTGGAGAACAATCCACGTGCTTGAG |
| Barcode B-55 | CAAGCGTTGGCTTCTCGCATCTGGTGCGAAATCCACGTGCTTGAG |
| Barcode B-56 | CAAGCGTTGGCTTCTCGCATCTGTACGCAAATCCACGTGCTTGAG |
| Barcode B-57 | CAAGCGTTGGCTTCTCGCATCTGTCGTAGAATCCACGTGCTTGAG |
| Barcode B-58 | CAAGCGTTGGCTTCTCGCATCTGTCTGTCAATCCACGTGCTTGAG |
| Barcode B-59 | CAAGCGTTGGCTTCTCGCATCTGTGTTCTAATCCACGTGCTTGAG |
| Barcode B-60 | CAAGCGTTGGCTTCTCGCATCTTAGGATGAATCCACGTGCTTGAG |
| Barcode B-61 | CAAGCGTTGGCTTCTCGCATCTTATCAGCAATCCACGTGCTTGAG |
| Barcode B-62 | CAAGCGTTGGCTTCTCGCATCTTCCGTCTAATCCACGTGCTTGAG |
| Barcode B-63 | CAAGCGTTGGCTTCTCGCATCTTCTTCACAATCCACGTGCTTGAG |
| Barcode B-64 | CAAGCGTTGGCTTCTCGCATCTTGAAGAGAATCCACGTGCTTGAG |
| Barcode B-65 | CAAGCGTTGGCTTCTCGCATCTTGGAACAAATCCACGTGCTTGAG |
| Barcode B-66 | CAAGCGTTGGCTTCTCGCATCTTGGCTTCAATCCACGTGCTTGAG |
| Barcode B-67 | CAAGCGTTGGCTTCTCGCATCTTGGTGGTAATCCACGTGCTTGAG |
| Barcode B-68 | CAAGCGTTGGCTTCTCGCATCTTTCACGCAATCCACGTGCTTGAG |
| Barcode B-69 | CAAGCGTTGGCTTCTCGCATCTAACTCACCATCCACGTGCTTGAG |
| Barcode B-70 | CAAGCGTTGGCTTCTCGCATCTAAGAGATCATCCACGTGCTTGAG |
| Barcode B-71 | CAAGCGTTGGCTTCTCGCATCTAAGGACACATCCACGTGCTTGAG |
| Barcode B-72 | CAAGCGTTGGCTTCTCGCATCTAATCCGTCATCCACGTGCTTGAG |

| | |
|---|---|
| Barcode B-73 | CAAGCGTTGGCTTCTCGCATCTAATGTTGCATCCACGTGCTTGAG |
| Barcode B-74 | CAAGCGTTGGCTTCTCGCATCTACACGACCATCCACGTGCTTGAG |
| Barcode B-75 | CAAGCGTTGGCTTCTCGCATCTACAGATTCATCCACGTGCTTGAG |
| Barcode B-76 | CAAGCGTTGGCTTCTCGCATCTAGATGTACATCCACGTGCTTGAG |
| Barcode B-77 | CAAGCGTTGGCTTCTCGCATCTAGCACCTCATCCACGTGCTTGAG |
| Barcode B-78 | CAAGCGTTGGCTTCTCGCATCTAGCCATGCATCCACGTGCTTGAG |
| Barcode B-79 | CAAGCGTTGGCTTCTCGCATCTAGGCTAACATCCACGTGCTTGAG |
| Barcode B-80 | CAAGCGTTGGCTTCTCGCATCTATAGCGACATCCACGTGCTTGAG |
| Barcode B-81 | CAAGCGTTGGCTTCTCGCATCTATCATTCCATCCACGTGCTTGAG |
| Barcode B-82 | CAAGCGTTGGCTTCTCGCATCTATTGGCTCATCCACGTGCTTGAG |
| Barcode B-83 | CAAGCGTTGGCTTCTCGCATCTCAAGGAGCATCCACGTGCTTGAG |
| Barcode B-84 | CAAGCGTTGGCTTCTCGCATCTCACCTTACATCCACGTGCTTGAG |
| Barcode B-85 | CAAGCGTTGGCTTCTCGCATCTCCATCCTCATCCACGTGCTTGAG |
| Barcode B-86 | CAAGCGTTGGCTTCTCGCATCTCCGACAACATCCACGTGCTTGAG |
| Barcode B-87 | CAAGCGTTGGCTTCTCGCATCTCCTAATCCATCCACGTGCTTGAG |
| Barcode B-88 | CAAGCGTTGGCTTCTCGCATCTCCTCTATCATCCACGTGCTTGAG |
| Barcode B-89 | CAAGCGTTGGCTTCTCGCATCTCGACACACATCCACGTGCTTGAG |
| Barcode B-90 | CAAGCGTTGGCTTCTCGCATCTCGGATTGCATCCACGTGCTTGAG |
| Barcode B-91 | CAAGCGTTGGCTTCTCGCATCTCTAAGGTCATCCACGTGCTTGAG |
| Barcode B-92 | CAAGCGTTGGCTTCTCGCATCTGAACAGGCATCCACGTGCTTGAG |
| Barcode B-93 | CAAGCGTTGGCTTCTCGCATCTGACAGTGCATCCACGTGCTTGAG |
| Barcode B-94 | CAAGCGTTGGCTTCTCGCATCTGAGTTAGCATCCACGTGCTTGAG |
| Barcode B-95 | CAAGCGTTGGCTTCTCGCATCTGATGAATCATCCACGTGCTTGAG |
| Barcode B-96 | CAAGCGTTGGCTTCTCGCATCTGCCAAGACATCCACGTGCTTGAG |
| Barcode B-97 | CAAGCGTTGGCTTCTCGCATCTCGGAAGAAATCCACGTGCTTGAG |
| Barcode B-98 | CAAGCGTTGGCTTCTCGCATCTGTGACAAGATCCACGTGCTTGAG |
| Barcode B-99 | CAAGCGTTGGCTTCTCGCATCTGAACCAGAATCCACGTGCTTGAG |
| Barcode B-100 | CAAGCGTTGGCTTCTCGCATCTTTGCTGGAATCCACGTGCTTGAG |

**Extended Data Table 4. Mouse Universal cocktail applied in this study.**

| DNA_ID | Name | Barcode sequence |
|---|---|---|
| A0001 | anti-mouse CD4 | AACAAGACCCTTGAG |
| A0002 | anti-mouse CD8a | TACCCGTAATAGCGT |
| A0003 | anti-mouse CD366 (Tim-3) | ATTGGCACTCAGATG |
| A0004 | anti-mouse CD279 (PD-1) | GAAAGTCAAAGCACT |
| A0013 | anti-mouse Ly-6C | AAGTCGTGAGGCATG |
| A0014 | anti-mouse/human CD11b | TGAAGGCTCATTTGT |
| A0015 | anti-mouse Ly-6G | ACATTGACGCAACTA |
| A0070 | anti-human/mouse CD49f | TTCCGAGGATGATCT |
| A0073 | anti-mouse/human CD44 | TGGCTTCAGGTCCTA |
| A0074 | anti-mouse CD54 | ATAACCGACACAGTG |
| A0075 | anti-mouse CD90.2 | CCGATCAGCCGTTTA |
| A0077 | anti-mouse CD73 | ACACTTAACGTCTGG |
| A0078 | anti-mouse CD49d | CGCTTGGACGCTTAA |
| A0079 | anti-mouse CD200 (OX2) | TCAATTCCGGTAGTC |
| A0090 | Mouse IgG1, κ isotype Ctrl | GCCGGACGACATTAA |
| A0091 | Mouse IgG2a, κ isotype Ctrl | CTCCTACCTAAACTG |
| A0092 | Mouse IgG2b, κ isotype Ctrl | ATATGTATCACGCGA |
| A0093 | anti-mouse CD19 | ATCAGCCATGTCAGT |
| A0095 | Rat IgG2b, κ Isotype Ctrl | GATTCTTGACGACCT |
| A0096 | anti-mouse CD45 | TGGCTATGGAGCAGA |
| A0097 | anti-mouse CD25 | ACCATGAGACACAGT |
| A0103 | anti-mouse/human CD45R/B220 | CCTACACCTCATAAT |
| A0104 | anti-mouse CD102 | GATATTCAGTGCGAC |
| A0105 | anti-mouse CD115 (CSF-1R) | TTCCGTTGTTGTGAG |
| A0106 | anti-mouse CD11c | GTTATGGACGCTTGC |
| A0107 | anti-mouse CD21/CD35 (CR2/CR1) | GGATAATTTCGATCC |
| A0108 | anti-mouse CD23 | TCTCTTGGAAGATGA |
| A0110 | anti-mouse CD43 | TTGGAGGGTTGTGCT |
| A0111 | anti-mouse CD5 | CAGCTCAGTGTGTTG |
| A0112 | anti-mouse CD62L | TGGGCCTAAGTCATC |
| A0113 | anti-mouse CD93 (AA4.1, early B lineage) | GGTATTTCCTGTGGT |
| A0114 | anti-mouse F4/80 | TTAACTTCAGCCCGT |
| A0115 | anti-mouse FcεRIα | AGTCACCTCGAAGCT |
| A0117 | anti-mouse I-A/I-E | GGTCACCAGTATGAT |

| | | |
|---|---|---|
| A0118 | anti-mouse NK-1.1 | GTAACATTACTCGTC |
| A0119 | anti-mouse Siglec H | CCGCACCTACATTAG |
| A0120 | anti-mouse TCR β chain | TCCTATGGGACTCAG |
| A0121 | anti-mouse TCR γ/δ | AACCCAAATAGCTGA |
| A0122 | anti-mouse TER-119/Erythroid Cells | GCGCGTTTGTGCTAT |
| A0130 | anti-mouse Ly-6A/E (Sca-1) | TTCCTTTCCTACGCA |
| A0157 | anti-mouse CD45.2 | CACCGTCATTCAACC |
| A0182 | anti-mouse CD3 | GTATGTCCGCTCGAT |
| A0190 | anti-mouse CD274 (B7-H1, PD-L1) | TCGATTCCACCAACT |
| A0191 | anti-mouse/rat/human CD27 | CAAGGTATGTCACTG |
| A0192 | anti-mouse CD20 | TCCACTCCCTGTATA |
| A0193 | anti-mouse CD357 (GITR) | GGCACTCTGTAACAT |
| A0194 | anti-mouse CD137 | TCCCTGTATAGATGA |
| A0195 | anti-mouse CD134 (OX-40) | CTCACCTACCTATGG |
| A0197 | anti-mouse CD69 | TTGTATTCCGCCATT |
| A0198 | anti-mouse CD127 (IL-7Rα) | GTGTGAGGCACTCTT |
| A0200 | anti-mouse CD86 | CTGGATTTGTGTATC |
| A0201 | anti-mouse CD103 | TTCATTAGCCCGCTG |
| A0202 | anti-mouse CD64 (FcγRI) | AGCAATTAACGGGAG |
| A0203 | anti-mouse CD150 (SLAM) | CAACGCCTAGAAACC |
| A0212 | anti-mouse CD24 | TATATCTTTGCCGCA |
| A0214 | anti-human/mouse integrin β7 | TCCTTGGATGTACCG |
| A0226 | anti-mouse CD106 | CGTTCCTACCTACCT |
| A0230 | anti-mouse CD8b (Ly-3) | TTCCCTCTATGGAGC |
| A0236 | Rat IgG1, κ isotype Ctrl | ATCAGATGCCCTCAT |
| A0237 | Rat IgG1, λ Isotype Ctrl | GGGAGCGATTCAACT |
| A0238 | Rat IgG2a, κ Isotype Ctrl | AAGTCAGGTTCGTTT |
| A0240 | Rat IgG2c, κ Isotype Ctrl | TCCAGGCTAGTCATT |
| A0241 | Armenian Hamster IgG Isotype Ctrl | CCTGTCATTAAGACT |
| A0250 | anti-mouse/human KLRG1 (MAFA) | GTAGTAGGCTAGACC |
| A0378 | anti-mouse CD223 (LAG-3) | ATTCCGTCCCTAAGG |
| A0417 | anti-mouse CD163 | GAGCAAGATTAAGAC |
| A0421 | anti-mouse CD49b | CGCGTTAGTAGAGTC |
| A0422 | anti-mouse CD172a (SIRPα) | GATTCCCTTGTAGCA |
| A0429 | anti-mouse CD48 | AGAACCGCCGTAGTT |
| A0431 | anti-mouse CD170 (Siglec-F) | TCAATCTCCGTCGCT |
| A0440 | anti-mouse CD169/Siglec-1 | ATTGACGACAGTCAT |

| | | |
|---|---|---|
| A0441 | anti-mouse CD71 | ACCGACCAGTAGACA |
| A0443 | anti-mouse CD41 | ACTTGGATGGACACT |
| A0450 | anti-mouse IgM | AGCTACGCATTCAAT |
| A0551 | anti-mouse CD301a | TGTATTTACTCACCG |
| A0552 | anti-mouse CD304 (Neuropilin-1) | CCAGCTCATTCAACG |
| A0555 | anti-mouse CD36 | TTTGCCGCTACGACA |
| A0557 | anti-mouse CD38 | CGTATCCGTCTCCTA |
| A0558 | anti-mouse CD55 (DAF) | ATTGTTGTCAGACCA |
| A0559 | anti-mouse CD63 | ATCCGACACGTATTA |
| A0560 | anti-mouse CD68 | CTTTCTTTCACGGGA |
| A0561 | anti-mouse CD79b (Igβ) | TAACTCAGTGCGAGT |
| A0562 | anti-mouse CD83 | TCTCAGGCTTCCTAG |
| A0563 | anti-mouse CX3CR1 | CACTCTCAGTCCTAT |
| A0566 | anti-mouse CD301b | CTTGCCTTGCGATTT |
| A0567 | anti-mouse Tim-4 | TGCTGGAGGGTATTC |
| A0568 | anti-mouse/rat XCR1 | TCCATTACCCACGTT |
| A0570 | anti-mouse/rat CD29 | ACGCATTCCTTGTGT |
| A0571 | anti-mouse IgD | TCATATCCGTTGTCC |
| A0595 | anti-mouse CD11a | AGAGTCTCCCTTTAG |
| A0807 | anti-mouse CD200R (OX2R) | ATTCTTTCCCTCTGT |
| A0809 | anti-mouse CD200R3 | ATCAACTTGGAGCAG |
| A0810 | anti-mouse CD138 (Syndecan-1) | GCGTTTGTATGTACT |
| A0811 | anti-mouse CD317 (BST2, PDCA-1) | TGTGGTAGCCCTTGT |
| A0813 | anti-mouse CD9 | TAGCAGTCACTCCTA |
| A0825 | anti-mouse CD371 (CLEC12A) | GCGAGAAATCTGCAT |
| A0827 | anti-mouse CD22 | AGGTCCTCTCTGGAT |
| A0837 | anti-mouse IL-33Rα (IL1RL1, ST2) | GCGATGGAGCATGTT |
| A0839 | anti-mouse Ly49H | CCAGTAGGCTTATTA |
| A0841 | anti-mouse Ly49D | TATATCCCTCAACGC |
| A0842 | anti-mouse Ly-49A | AATTCCGTCAGATGA |
| A0846 | anti-mouse CD185 (CXCR5) | ACGTAGTCACCTAGT |
| A0850 | anti-mouse CD49a | CCATTCATTTGTGGC |
| A0851 | anti-mouse CD1d (CD1.1, Ly-38) | CAACTTGGCCGAATC |
| A0852 | anti-mouse CD226 (DNAM-1) | ACGCAGTATTTCCGA |
| A0854 | anti-mouse CD199 (CCR9) | CCCTCTGGTATGGTT |
| A0877 | anti-mouse JAML | GTTATGGTTCGTGTT |
| A0881 | anti-mouse CD272 (BTLA) | TGACCCTATTGAGAA |

| | | |
|---|---|---|
| A0882 | anti-mouse PIR-A/B | TGTAGAGTCAGACCT |
| A0883 | anti-mouse CD26 (DPP-4) | ATGGCCTGTCATAAT |
| A0885 | anti-mouse CD270 (HVEM) | GATCCGTGTTGCCTA |
| A0892 | anti-mouse CD2 | TTGCCGTGTGTTTAA |
| A0893 | anti-mouse CD120b (TNF R Type II/p75) | GAAGCTGTATCCGAA |
| A0903 | anti-mouse CD40 | ATTTGTATGCTGGAG |
| A0904 | anti-mouse CD31 | GCTGTAGTATCATGT |
| A0905 | anti-mouse CD107a (LAMP-1) | AAATCTGTGCCGTAC |
| A0910 | anti-mouse/rat CD61 | TTCTTTACCCGCCTG |
| A0915 | anti-mouse VISTA (PD-1H) | ACATTTCCCTTGCCT |
| A0926 | anti-mouse CD186 (CXCR6) | TGTCAGGTTGTATTC |
| A0927 | anti-mouse CD159a (NKG2AB6) | GTGTTTGTGTTCCTG |
| A0930 | anti-mouse Ly108 | CGATTCTTTGCGAGT |
| A1006 | anti-mouse CD160 | GCGTATGTCAGTACC |
| A1007 | anti-mouse CD85k (gp49 Receptor) | ATGTCAACTCTGGGA |
| A1008 | anti-mouse CD51 | GGAGTCAGGGTATTA |
| A1009 | anti-mouse CD94 | CACAGTTGTCCGTGT |
| A1010 | anti-mouse CD205 (DEC-205) | CATATTGGCCGTAGT |
| A1011 | anti-mouse CD155 (PVR) | TAGCTTGGGATTAAG |
| A1064 | anti-mouse/rat CD81 | TTGTCACCAACTTCC |
| | MBP | TAGTACGGATCCAGT |
| | MOG | GGTACTGAACTTTAG |
| | NEUN | CGAGTGCACCTTGAG |
| | PDGFRα | CTTGATCGTTGACGA |
| | SATB2 | GACAAATTGTCAGTT |
| | TBR1 | AATCGAATTGGCATA |
| | CUX2/1 | AAACTAAGATGACGG |
| | CTIP2 | TTAAGATCAGGAATC |

**Extended Data Table 5. Human Universal cocktail applied in this study.**

| ID | Name | Barcode sequence |
|---|---|---|
| ADT_A0006 | Hu.CD86 | GTCTTTGTCAGTGCA |
| ADT_A0007 | Hu.CD274 | GTTGTCCGACAATAC |
| ADT_A0020 | Hu.CD270 | TGATAGAAACAGACC |
| ADT_A0023 | Hu.CD155 | ATCACATCGTTGCCA |
| ADT_A0024 | Hu.CD112 | AACCTTCCGTCTAAG |
| ADT_A0026 | Hu.CD47 | GCATTCTGTCACCTA |
| ADT_A0029 | Hu.CD48 | CTACGACGTAGAAGA |
| ADT_A0031 | Hu.CD40 | CTCAGATGGAGTATG |
| ADT_A0032 | Hu.CD154 | GCTAGATAGATGCAA |
| ADT_A0033 | Hu.CD52 | CTTTGTACGAGCAAA |
| ADT_A0034 | Hu.CD3_UCHT1 | CTCATTGTAACTCCT |
| ADT_A0046 | Hu.CD8 | GCGCAACTTGATGAT |
| ADT_A0047 | Hu.CD56 | TCCTTTCCTGATAGG |
| ADT_A0050 | Hu.CD19 | CTGGGCAATTACTCG |
| ADT_A0052 | Hu.CD33 | TAACTCAGGGCCTAT |
| ADT_A0053 | Hu.CD11c | TACGCCTATAACTTG |
| ADT_A0058 | Hu.HLA.ABC | TATGCGAGGCTTATC |
| ADT_A0063 | Hu.CD45RA | TCAATCCTTCCGCTT |
| ADT_A0064 | Hu.CD123 | CTTCACTCTGTCAGG |
| ADT_A0066 | Hu.CD7 | TGGATTCCCGGACTT |
| ADT_A0070 | HuMs.CD49f | TTCCGAGGATGATCT |
| ADT_A0071 | Hu.CD194 | AGCTTACCTGCACGA |
| ADT_A0072 | Hu.CD4_RPA.T4 | TGTTCCCGCTCAACT |
| ADT_A0073 | HuMs.CD44 | TGGCTTCAGGTCCTA |
| ADT_A0081 | Hu.CD14_M5E2 | TCTCAGACCTCCGTA |
| ADT_A0083 | Hu.CD16 | AAGTTCACTCTTTGC |
| ADT_A0085 | Hu.CD25 | TTTGTCCTGTACGCC |
| ADT_A0087 | Hu.CD45RO | CTCCGAATCATGTTG |
| ADT_A0088 | Hu.CD279 | ACAGCGCCGTATTTA |
| ADT_A0089 | Hu.TIGIT | TTGCTTACCGCCAGA |
| ADT_A0090 | Isotype_MOPC.21 | GCCGGACGACATTAA |
| ADT_A0091 | Isotype_MOPC.173 | CTCCTACCTAAACTG |
| ADT_A0092 | Isotype_MPC.11 | ATATGTATCACGCGA |
| ADT_A0095 | Isotype_RTK4530 | GATTCTTGACGACCT |
| ADT_A0100 | Hu.CD20_2H7 | TTCTGGGTCCCTAGA |

| | | |
|---|---|---|
| ADT_A0101 | Hu.CD335 | ACAATTTGAACAGCG |
| ADT_A0124 | Hu.CD31 | ACCTTTATGCCACGG |
| ADT_A0127 | Hu.Podoplanin | GGTTACTCGTTGTGT |
| ADT_A0134 | Hu.CD146 | CCTTGGATAACATCA |
| ADT_A0136 | Hu.IgM | TAGCGAGCCCGTATA |
| ADT_A0138 | Hu.CD5 | CATTAACGGGATGCC |
| ADT_A0140 | Hu.CD183 | GCGATGGTAGATTAT |
| ADT_A0141 | Hu.CD195 | CCAAAGTAAGAGCCA |
| ADT_A0142 | Hu.CD32 | GCTTCCGAATTACCG |
| ADT_A0143 | Hu.CD196 | GATCCCTTTGTCACT |
| ADT_A0144 | Hu.CD185 | AATTCAACCGTCGCC |
| ADT_A0145 | Hu.CD103 | GACCTCATTGTGAAT |
| ADT_A0146 | Hu.CD69 | GTCTCTTGGCTTAAA |
| ADT_A0147 | Hu.CD62L | GTCCCTGCAACTTGA |
| ADT_A0149 | Hu.CD161 | GTACGCAGTCCTTCT |
| ADT_A0151 | Hu.CD152 | ATGGTTCACGTAATC |
| ADT_A0152 | Hu.CD223 | CATTTGTCTGCCGGT |
| ADT_A0153 | Hu.KLRG1 | CTTATTTCCTGCCCT |
| ADT_A0154 | Hu.CD27 | GCACTCCTGCATGTA |
| ADT_A0155 | Hu.CD107a | CAGCCCACTGCAATA |
| ADT_A0156 | Hu.CD95 | CCAGCTCATTAGAGC |
| ADT_A0158 | Hu.CD134 | AACCCACCGTTGTTA |
| ADT_A0159 | Hu.HLA.DR | AATAGCGAGCAAGTA |
| ADT_A0160 | Hu.CD1c | GAGCTACTTCACTCG |
| ADT_A0161 | Hu.CD11b | GACAAGTGATCTGCA |
| ADT_A0162 | Hu.CD64 | AAGTATGCCCTACGA |
| ADT_A0163 | Hu.CD141 | GGATAACCGCGCTTT |
| ADT_A0165 | Hu.CD314 | CGTGTTTGTTCCTCA |
| ADT_A0167 | Hu.CD35 | ACTTCCGTCGATCTT |
| ADT_A0168 | Hu.CD57 | AACTCCCTATGGAGG |
| ADT_A0170 | Hu.CD272 | GTTATTGGACTAAGG |
| ADT_A0171 | HuMsRt.CD278 | CGCGCACCCATTAAA |
| ADT_A0172 | Hu.CD275_B7.RP1 | GTTAGTGTTAGCTTG |
| ADT_A0174 | Hu.CD58 | GTTCCTATGGACGAC |
| ADT_A0176 | Hu.CD39 | TTACCTGGTATCCGT |
| ADT_A0179 | Hu.CX3CR1 | AGTATCGTCTCTGGG |
| ADT_A0180 | Hu.CD24 | AGATTCCTTCGTGTT |

| | | |
|---|---|---|
| ADT_A0181 | Hu.CD21 | AACCTAGTAGTTCGG |
| ADT_A0185 | Hu.CD11a | TATATCCTTGTGAGC |
| ADT_A0187 | Hu.CD79b | ATTCTTCAACCGAAG |
| ADT_A0189 | Hu.CD244 | TCGCTTGGATGGTAG |
| ADT_A0206 | Hu.CD169 | TACTCAGCGTGTTTG |
| ADT_A0214 | HuMs.integrin.b7 | TCCTTGGATGTACCG |
| ADT_A0215 | Hu.CD268 | CGAAGTCGATCCGTA |
| ADT_A0216 | Hu.CD42b | TCCTAGTACCGAAGT |
| ADT_A0217 | Hu.CD54 | CTGATAGACTTGAGT |
| ADT_A0218 | Hu.CD62P | CCTTCCGTATCCCTT |
| ADT_A0219 | Hu.CD119 | TGTGTATTCCCTTGT |
| ADT_A0224 | Hu.TCR.AB | CGTAACGTAGAGCGA |
| ADT_A0236 | Isotype_RTK2071 | ATCAGATGCCCTCAT |
| ADT_A0237 | Isotype_G0114F7 | GGGAGCGATTCAACT |
| ADT_A0238 | Isotype_RTK2758 | AAGTCAGGTTCGTTT |
| ADT_A0240 | Isotype_RTK4174 | TCCAGGCTAGTCATT |
| ADT_A0241 | Isotype_HTK888 | CCTGTCATTAAGACT |
| ADT_A0242 | Hu.CD192 | GAGTTCCCTTACCTG |
| ADT_A0246 | Hu.CD122 | TCATTTCCTCCGATT |
| ADT_A0247 | Hu.CD267 | AGTGATGGAGCGAAC |
| ADT_A0352 | Hu.FceRIa | CTCGTTTCCGTATCG |
| ADT_A0353 | Hu.CD41 | ACGTTGTGGCCTTGT |
| ADT_A0355 | Hu.CD137 | CAGTAAGTTCGGGAC |
| ADT_A0357 | Hu.CD43 | GATTAACCAGCTCAT |
| ADT_A0358 | Hu.CD163 | GCTTCTCCTTCCTTA |
| ADT_A0359 | Hu.CD83 | CCACTCATTTCCGGT |
| ADT_A0364 | Hu.CD13 | TTTCAACGCCCTTTC |
| ADT_A0367 | Hu.CD2 | TACGATTTGTCAGGG |
| ADT_A0368 | Hu.CD226_11A8 | TCTCAGTGTTTGTGG |
| ADT_A0369 | Hu.CD29 | GTATTCCCTCAGTCA |
| ADT_A0370 | Hu.CD303 | GAGATGTCCGAATTT |
| ADT_A0371 | Hu.CD49b | GCTTTCTTCAGTATG |
| ADT_A0372 | Hu.CD61 | AGGTTGGAGTAGACT |
| ADT_A0373 | Hu.CD81 | GTATCCTTCCTTGGC |
| ADT_A0383 | Hu.CD55 | GCTCATTACCCATTA |
| ADT_A0384 | Hu.IgD | CAGTCTCCGTAGAGT |
| ADT_A0385 | Hu.CD18 | TATTGGGACACTTCT |

| | | |
|---|---|---|
| ADT_A0386 | Hu.CD28 | TGAGAACGACCCTAA |
| ADT_A0389 | Hu.CD38_HIT2 | TGTACCCGCTTGTGA |
| ADT_A0390 | Hu.CD127 | GTGTGTTGTCCTATG |
| ADT_A0391 | Hu.CD45_HI30 | TGCAATTACCCGGAT |
| ADT_A0393 | Hu.CD22 | GGGTTGTTGTCTTTG |
| ADT_A0394 | Hu.CD71 | CCGTGTTCCTCATTA |
| ADT_A0396 | Hu.CD26 | GGTGGCTAGATAATG |
| ADT_A0398 | Hu.CD115 | AATCACGGTCCTTGT |
| ADT_A0404 | Hu.CD63 | GAGATGTCTGCAACT |
| ADT_A0406 | Hu.CD304 | GGACTAAGTTTCGTT |
| ADT_A0407 | Hu.CD36 | TTCTTTGCCTTGCCA |
| ADT_A0408 | Hu.CD172a | CGTGTTTAACTTGAG |
| ADT_A0419 | Hu.CD72 | CAGTCGTGGTAGATA |
| ADT_A0420 | Hu.CD158 | TATCAACCAACGCTT |
| ADT_A0446 | Hu.CD93 | GCGCTACTTCCTTGA |
| ADT_A0447 | Hu.CD200 | CACGTAGACCTTTGC |
| ADT_A0575 | Hu.CD49a | ACTGATGGACTCAGA |
| ADT_A0576 | Hu.CD49d | CCATTCAACTTCCGG |
| ADT_A0577 | Hu.CD73 | CAGTTCCTCAGTTCG |
| ADT_A0579 | Hu.CD9 | GAGTCACCAATCTGC |
| ADT_A0581 | Hu.TCR.Va7.2 | TACGAGCAGTATTCA |
| ADT_A0582 | Hu.TCR.Vd2 | TCAGTCAGATGGTAT |
| ADT_A0586 | Hu.CD354 | TAGCCGTTTCCTTTG |
| ADT_A0590 | Hu.CD305_LAIR1 | ATTTCCATTCCCTGT |
| ADT_A0591 | Hu.LOX.1 | ACCCTTTACCGAATA |
| ADT_A0599 | Hu.CD158e1 | GGACGCTTTCCTTGA |
| ADT_A0817 | Hu.CD109 | CACTTAACTCTGGGT |
| ADT_A0822 | Hu.CD142 | CACTGCCGTCGATTA |
| ADT_A0830 | Hu.CD319 | AGTATGCCATGTCTT |
| ADT_A0845 | Hu.CD99 | ACCCGTCCCTAAGAA |
| ADT_A0853 | Hu.CLEC12A | CATTAGAGTCTGCCA |
| ADT_A0861 | Hu.CD151 | CTTACCTAGTCATTC |
| ADT_A0864 | Hu.CD352 | AGTTTCCACTCAGGC |
| ADT_A0866 | Hu.CLEC1B | TGCCAGTATCACGTA |
| ADT_A0867 | Hu.CD94 | CTTTCCGGTCCTACA |
| ADT_A0868 | Hu.IgE | GGATGTACCGCGTAT |
| ADT_A0870 | Hu.CD150 | GTCATTGTATGTCTG |

| | | |
|---|---|---|
| ADT_A0871 | Hu.CD162 | ATATGTCAGAGCACC |
| ADT_A0872 | Hu.CD84 | CTCCCTAGTTCCTTT |
| ADT_A0894 | Hu.Ig.LightChain.k | AGCTCAGCCAGTATG |
| ADT_A0896 | Hu.CD85j | CCTTGTGAGGCTATG |
| ADT_A0897 | Hu.CD23 | TCTGTATAACCGTCT |
| ADT_A0898 | Hu.Ig.LightChain.l | CAGCCAGTAAGTCAC |
| ADT_A0902 | Hu.CD328 | CTTAGCATTTCACTG |
| ADT_A0912 | Hu.GPR56 | GCCTAGTTTCCGTTT |
| ADT_A0920 | Hu.CD82 | TCCCACTTCCGCTTT |
| ADT_A0923 | Hu.NKp80 | TATAGTTCCTCTGTG |
| ADT_A0931 | Hu.CD131 | CTGCATGAGACCAAA |
| ADT_A0935 | Hu.CD74 | CTGTAGCATTTCCCT |
| ADT_A0940 | Hu.CD116 | ATGGACAGTTCGTGT |
| ADT_A0941 | Hu.CD37 | ACAGTCACTGGGCAA |
| ADT_A0944 | Hu.CD101 | CTACTTCCCTGTCAA |
| ADT_A1018 | Hu.HLA.DR.DP.DQ | AGCTACGAGCAGTAG |
| ADT_A1046 | Hu.CD88 | GCCGCATGAGAAACA |

**Extended Data Table 6. Chemicals and reagents.**

| Name | Catalog number | Vender |
|---|---|---|
| Formaldehyde solution | PI28906 | Thermo Fisher Scientific |
| HEPES pH 7.5 | BBH-75-250 | Boston BioProducts |
| Glycine | 50046 | Sigma-Aldrich |
| NaCl | AM9760G | Thermo Fisher Scientific |
| Digitonin | G9441 | Promega |
| $MgCl_2$ | AM9530G | Thermo Fisher Scientific |
| Spermidine | S0266 | Sigma-Aldrich |
| EDTA-free Protease Inhibitor Cocktail | 11873580001 | Millipore Sigma |
| NP40 | 11332473001 | Sigma-Aldrich |
| EDTA Solution pH 8.0 | AB00502 | AmericanBio |
| Bovine Serum Albumin (BSA) | A8806 | Sigma-Aldrich |
| Anti-H3K27me3 antibody | 9733 | Cell Signaling Technology |
| Secondary antibody (Guinea Pig anti-Rabbit IgG) | ABIN101961 | Antibodies-Online |
| pA-Tn5 Transposase – unloaded | C01070002 | Diagenode |
| Triton X-100 | T8787 | Sigma-Aldrich |
| T4 DNA Ligase | M0202L | New England Biolabs |
| T4 DNA Ligase Reaction Buffer | B0202S | New England Biolabs |
| NEBuffer 3.1 | B7203S | New England Biolabs |
| DPBS | 14190144 | Thermo Fisher Scientific |
| Proteinase K | EO0491 | Thermo Fisher Scientific |
| Ampure XP beads | A63880 | Beckman Coulter |
| NEBNext High-Fidelity 2X PCR Master Mix | M0541L | New England Biolabs |
| SYBR Green I Nucleic Acid Gel Stain | S7563 | Thermo Fisher Scientific |
| DNA Clean & Concentrator-5 | D4014 | Zymo Research |
| Tn5 Transposase - unloaded | C01070010 | Diagenode |
| Tagmentation Buffer (2x) | C01019043 | Diagenode |
| Sodium dodecyl sulfate | 71736 | Sigma-Aldrich |
| Maxima H Minus Reverse Transcriptase (200 U/L) | EP0751 | Thermo Fisher Scientific |
| dNTP mix | R0192 | Thermo Fisher Scientific |
| SUPERased In RNase Inhibitor | AM2694 | Thermo Fisher Scientific |

| | | |
|---|---|---|
| Ampure XP beads | A63880 | Beckman Coulter |
| Dynabeads MyOne C1 | 65001 | Thermo Fisher Scientific |
| RNase Inhibitor | Y9240L | Enzymatics |
| Kapa Hotstart HiFi ReadyMix | KK2601 | Kapa Biosystems |
| Nextera XT DNA Preparation Kit | FC-131-1024 | Illumina |
| PDGFRα | ab234965 | abcam |
| OLIG2 | ab220796 | abcam |
| APC | ab239828 | abcam |
| MBP | ab230378 | abcam |
| IBA1 | ab220815 | abcam |
| GFAP | ab218309 | abcam |
| CTIP2 | ab269367 | abcam |
| CUX1+Cux2 | ab309140 | abcam |
| TBR1 | ab239000 | abcam |
| NeuN | ab209898 | abcam |
| MOG | ab255266 | abcam |
| Satb2 | ab212177 | abcam |
| CD31 | 4250001 | Akoya |
| CD4 | 4250016 | Akoya |
| CD8a | 4250017 | Akoya |
| CD19 | 4250014 | Akoya |
| CD45R/B220 | 4450006 | Akoya |
| CD11c | 4550108 | Akoya |
| Ki67 | 4250019 | Akoya |
| CD11b | 4450015 | Akoya |
| Ly6g | 4550110 | Akoya |
| CD3 | 4550109 | Akoya |
| CD169 | 4550100 | Akoya |
| Mouse Universal Cocktail | 199901 | BioLegend |
| Human Universal Cocktail | 399907 | BioLegend |

**Extended Data Table 7.** The gene list of each cluster generated from the regression model for mouse brain cortical layers.

**Extended Data Table 8.** The gene list of each cluster generated from the regression model for mouse brain corpus callosum.

**Extended Data Video 1.** Video output from TRIC-DISCO, replicate #1.

**Extended Data Video 2.** Video output from TRIC-DISCO, replicate #2.

**Extended Data Video 3.** Video output from TRIC-DISCO, replicate #3.
